# STOCHASTIC ECO-EVOLUTIONARY DYNAMICS OF MULTIVARIATE TRAITS: A Framework for Modeling Population Processes Illustrated by the Study of Drifting G-Matrices

**DOI:** 10.1101/2025.06.07.658444

**Authors:** Bob Week

## Abstract

I derive a novel stochastic equation for the evolution of the additive genetic variance-covariance matrix **G** in response to mutation, selection, drift, and fluctuating population size. Common wisdom holds that the effect of drift on **G** is simply to reduce each of its entries by a common proportional amount while preserving its orientation. In contrast, I find that drift causes significant and directional shifts in the orientation of **G** by driving genetic correlations to their extremes. Biologically, this is a consequence of linkage build-up introduced by drift. I compare these theoretical results to empirical observations based on experiments conducted by Phillips et al., (2001). Additionally, to derive the model of **G**-matrix evolution, I developed a novel synthetic framework for modelling ecological and evolutionary dynamics of populations carrying multivariate traits. This framework is optimized for deriving new models across a wide range of topics in population biology. Foundations of the framework are formalized by the theory of measure-valued processes, but application of the framework only requires multivariate calculus, and heuristics are presented in the main text for making additional calculations involving stochastic processes. Collectively, this work establishes a powerful framework enabling efficient formal analysis of integrated population processes across evolution and ecology, and its potential for making new discoveries is illustrated by novel findings on fundamental aspects of **G**-matrix evolution.

## 1. Introduction

A fundamental principle of evolutionary biology is that random genetic drift erodes heritable variation at a rate inversely proportional to effective population size. At the level of allele frequencies, models for the distribution of genetic variation responding to drift are well-known (Kimura, 1964, 1968; Ewens, 2004; Hill and Robertson, 1966). In contrast, at the level of quantitative characters, models of additive genetic variation responding to drift focus on the average outcome using deterministic models (Latter, 1970; Bulmer, 1972; Lande, 1976, 1980; Chakraborty and Nei, 1982; Turelli and Barton, 1994; Lynch and Hill, 1986; Bürger, 2000; Barton and Turelli, 2004; Débarre and Otto, 2016; Walsh and Lynch, 2018). Empirical work has supported theoretical predictions for the average response of additive genetic variation to drift (Phillips et al., 2001; McGuigan et al., 2005; Whitlock et al., 2002), but has also emphasized the need to predict the distribution of outcomes (Phillips et al., 2001; Whitlock, 1995) especially for multivariate traits and genetic covariances (Phillips and McGuigan, 2006; Mallard et al., 2023a).

In the setting of multivariate traits, a common summary statistic for genetic architecture is the **G**-matrix. This matrix has the additive genetic variance of each trait on the associated diagonal entry. Off-diagonal entries quantify genetic covariances between traits which may be maintained by pleiotropic loci and linkage between loci affecting different traits. The standard view is that drift produces a proportional decrease in **G** (Phillips and McGuigan, 2006; Cano et al., 2004; McGuigan, 2006; Chapuis et al., 2008; Dugand et al., 2021; Mallard et al., 2023a), and thus independent populations that have diverged due to drift should have proportional **G**-matrices (Roff, 2000; Steppan et al., 2002; Aguirre et al., 2013). However, this insight rests on results from a deterministic model for the response of **G**-matrices to drift (Lande, 1979) and a model that assumes recombination happens sufficiently fast to break-up linkage produced by selection (Lande, 1980). Hence, there is a need to develop theoretical predictions for the stochastic evolution of **G**-matrices driven solely by drift, and especially for the effect of drift on genetic covariances (Mallard et al., 2023a,b).

The reason for this gap in evolutionary theory stems, in part, from the lack of formal approaches to place stochastic models of **G**-matrix evolution on a concrete mathematical foundation. In this paper, I aim to fill this gap by leveraging the powerful theory of measure-valued processes, while also keeping the presentation as accessible as possible. Furthermore, taking this approach to formally derive **G**-matrix dynamics revealed a much broader framework for modelling a wide-range of population processes. In particular, given the growing appreciation for the interplay between ecological and evolutionary processes (Reznick, 2015; Hendry, 2017; Kuosmanen et al., 2022), this framework makes an important contribution by enabling the formal interfacing of **G**-matrix evolution with models of eco-evolutionary feedbacks (e.g., Patel et al., 2018). I therefore focus this paper on presenting the framework in its full generality, and return to the study of **G**-matrix evolution as an example to illustrate its utility.

In its full generality, this framework offers tools to model the integrated ecological and evolutionary dynamics of populations with multivariate traits that respond to mutation, selection (including frequency and abundance dependence), demographic stochasticity, and consequential random genetic drift. In particular, this framework can be used to obtain generalizations of many classical models in evolution and ecology, such as Lotka-Volterra dynamics (Cattiaux and Méléard, 2009; Akjouj et al., 2024), coevolution (Gilman et al., 2012; Débarre et al., 2014), and evolutionary rescue (Klausmeier et al., 2020; Xu et al., 2023). Further details on how to apply this framework to generalize known models are given in the discussion section.

As in classical quantitative genetics, the framework here assumes a linear genotype-phenotype map with fixed mutational covariance. Under this modelling choice, development enters implicitly through the genotype-phenotype map rather than via explicit mechanistic developmental feedbacks (see Milocco and Salazar-Ciudad, 2022; González-Forero, 2022, 2024a,b). This paper therefore focuses on mutation-selection-drift dynamics of multivariate traits, not on the evolution of developmental architecture.

To apply this framework, the most important biological details relevant to a modeler are the mechanisms mediating fitness. In contrast, details involving the genetic architecture of traits are abstracted in a way that captures basic biological principles while optimizing analytical tractability. For instance, asexual reproduction is assumed and mutation is modeled following approaches similar to Kimura (1965) and Débarre and Otto (2016) by assuming offspring traits are distributed around their parental traits. This approach to modelling inheritance is similar to the infinitesimal model in which traits are determined by many loci of small effect, which implies offspring are normally distributed around their parental values (Barton et al., 2017). However, an important distinction is that the infinitesimal model restricts the response of allele frequencies to selection such that changes in genetic variance are only temporary and due to build-up of genetic linkage (Bulmer, 1971). Instead, I begin by assuming offspring traits are normally distributed around their parental traits (referred to as the Gaussian descendants model by Turelli, 2017), in which case the response of population variance to selection can be permanent. Abstract approaches similar to this have been successful for obtaining valuable analytical insights into genetic variation maintained by mutation-selection balance (Kimura, 1965; Lande, 1975; Turelli, 1984, 1986) and by mutation-drift balance (Lande, 1976, 1979; Barton, 1989; Débarre and Otto, 2016). Hence, by optimizing a trade-off between genetic detail and analytical tractability, this framework provides an accessible approach for obtaining and communicating a wide array of novel theoretical insights.

To establish this framework, I build on the work of Week et al. (2021), which presented a stochastic differential equation framework focused on modelling the simultaneous dynamics of abundances, 1-dimensional mean traits, and 1-dimensional trait variances responding to mutation, selection, demographic stochasticity, and random genetic drift. This 1-dimensional framework was based on the development of heuristics (i.e., practical rules that allow exact calculations without requiring familiarity with the underlying martingale theory or its formal derivation) for working with stochastic partial differential equations (spde). However, for multivariate traits, the spde approach breaks down (Dawson, 1993; Etheridge, 2000; Perkins, 2002). To overcome this challenge and establish a rigorous analytical framework for deriving population processes, I take an approach based on so-called *martingale problems* (Ethier and Kurtz, 1986; Dawson, 1993; Stroock and Varadhan, 1997; Rogers and Williams, 2000). Mathematical aspects of this approach are provided in Supplementary Material, Section 4. In the main text, I focus on the resulting dynamical equations and heuristics for performing calculations with minimal technical background.

### 1.1. Overview

I begin by outlining the derivation of the deterministic version of the framework without making any assumptions on the shape of trait distributions, which is summarized by a system of ordinary differential equations. This leads to expressions of selection in terms of covariances with fitness, which I refer to collectively as the Deterministic Covariance version (or *DC* for short). By assuming traits follow multivariate normal distributions, covariances with fitness are replaced by multivariate gradients of fitness with respect to mean traits and trait variances, and I refer to the resulting system of differential equations as the Deterministic Gradient version (or *DG* for short). To simplify presentation of the deterministic version of the framework, I assume traits are perfectly heritable. However, because this work is motivated by understanding the consequences of drift for **G**-matrix evolution, I briefly describe an approach to model imperfect heritability after introducing *DC* and *DG*. This model of imperfect heritability is adopted while introducing the stochastic extensions of the framework.

The stochastic extensions of the framework include the effects of demographic stochasticity (i.e., random reproductive output) and random genetic drift (which occurs here as a consequence of demographic stochasticity). I introduce two stochastic extensions. Both build on *DG* by assuming multivariate normal trait distributions and by expressing selection in terms of fitness gradients. The first form expresses dynamics in terms of Brownian motions as drivers of stochasticity (referred to as the Brownian Motion Gradient version, or *BG* for short), which is particularly useful for numerical analysis. The second form expresses dynamics in terms of a more general underlying martingale process (referred to as the Martingale Gradient version, or *MG* for short), and I use this form to introduce heuristics for deriving analytical models. To demonstrate the heuristics (summarized in Table 1), I derive a stochastic equation for the evolution of additive genetic correlations between trait values. To bring this paper full-circle, I then discuss how this exercise provides novel insights into the evolutionary response of **G**-matrices to random genetic drift and compare these theoretical results with observations obtained from experiments (Phillips et al., 2001; McGuigan et al., 2005; Whitlock et al., 2002).

**Table 1.**
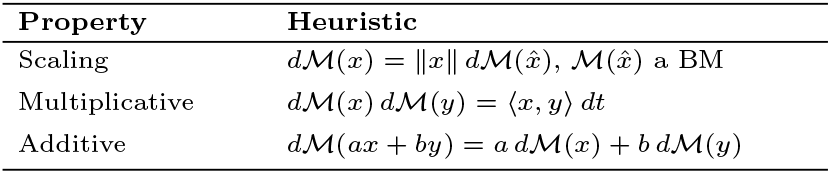
Heuristics summarizing the scaling, multiplicative, and additive properties of the stochastic differentials *d*ℳ(*x*).

The Supplementary Material provides details to support conclusions and arrive at expressions presented throughout the main text. Supplementary Material, Section 1 summarizes numerical implementations of the framework. Detailed derivation of the deterministic versions of the framework are given in Supplementary Material, Section 2. Section 3 of the Supplement formally interfaces the continuous-time framework presented here with discrete time models of classical quantitative genetics. The mathematical foundation of this framework is communicated in Supplementary Material, Section 4. Using this foundation, Section 5 of the Supplement presents detailed calculations for the derivation of the stochastic versions of the framework presented in the main text. In Supplementary Material, Section 6, I derive the dynamics of genetic correlations responding to drift. An outline for one approach to derive this framework from a diffusion-limit is given in Section 7 of the Supplement. Finally, Supplementary Material, Section 8 formalizes a connection with Gillespie’s work on the evolutionary consequences of offspring number variation (Gillespie, 1974, 1975, 1977)

## 2. The Framework

The framework tracks the dynamics of the population density across trait space for asexually reproducing populations. To model *d*-dimensional traits, I assume trait space is the entire Euclidean space ℝ^*d*^. Given the *d*-dimensional trait **z** = (*z*_1_, …, *z*_*d*_)^⊤^ (with ^⊤^ denoting matrix transposition so **z** is a column vector), I write *ν*(**z**) for the population density at **z**. Then the total abundance of the population is given by 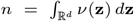. The distribution of the trait **z** is then *p*(**z**) = *ν*(**z**)*/n*, and this is also referred to as the relative abundance of **z**. Using *p*(**z**), the mean trait vector is given by 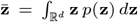, and the trait covariance matrix is 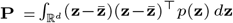, with *P*_*ij*_ being the covariance between *z*_*i*_ and *z*_*j*_. The chosen notation facilitates comparison with classical multivariate quantitative genetic models (Lande, 1980; Lande and Arnold, 1983; Jones et al., 2003; Arnold et al., 2008).

In the following section (2.1) I describe how to obtain the deterministic version of the framework using multivariate calculus. I do this in two parts. The first part (*DC*) makes no assumption on the shape of *p*(**z**). The second part (*DG*) assumes *p*(**z**) is the density of a multivariate normal distribution. I continue to make this assumption in section 2.2 where I introduce the stochastic extension of this framework.

### 2.1. Deterministic Dynamics

#### 2.1.1. The Deterministic Covariance Version (DC)

To establish a flexible, but tractable framework to model the dynamics of *n*, 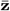, and **P**, I generalize the deterministic model used by Week et al., (2021), referred to there as the Deterministic Asexual Gaussian Allelic model (DAGA), to the multivariate setting. Specifically, DAGA focuses on dynamics due to just mutation and selection for asexually reproducing populations. Mutation is modeled as the variance **M** of a normal distribution determining offspring traits centered on their parental traits, and selection results from the covariance of fitness and phenotype.

In the multivariate trait setting, mutation is modeled as a *d × d* covariance matrix **M**, with the off-diagonal entries contributing to trait covariances. Here, I assume **M** does not depend on **z**. This setup fits within the conceptual framework that has been popular for theoretical and empirical studies of **G**-matrix evolution (Jones et al., 2007; Arnold et al., 2008; Mallard et al., 2023a). Fitness is quantified by a rate *m*(*ν*, **z**), which is the growth rate for the sub-population of individuals with trait value **z** in a population summarized by *ν*. The dependency of *m* on *ν* and **z** allows for the modelling of interwoven ecological and evolutionary dynamics. I will also refer to *m*(*ν*, **z**) as a *fitness function*, and this function may also depend on environmental parameters such as the trait values of individuals in interacting species. However, I omit notation accounting for such possibilities to simplify the frameworks presentation. Putting these components together, the multivariate generalization of DAGA is given by the partial differential equation

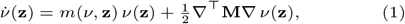

where 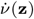 is the instantaneous rate of change of *ν*(**z**) over time, the symbol ∇ := (∂*/*∂*z*_1_, …, ∂*/*∂*z*_*d*_)^⊤^ is the gradient operator with respect to the *d*-dimensional trait **z**, and

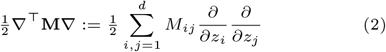

is the mutation operator. Equation (1) can be used to obtain a multivariate generalization of Kimura’s continuum-of-alleles model (Kimura, 1965) that assumes diffusion instead of convolution with an arbitrary mutation kernel. Justification for multivariate DAGA (Equation (1)) as a deterministic approximation of the full stochastic framework is given in Supplementary Material, Section 7.

If *M*_*ij*_ = 0 when *i* ≠ *j* and *M*_*ii*_ = *M*_*jj*_ is constant across all *ij*, then the above mutation operator becomes proportional to the Laplacian operator on ℝ^*d*^, which causes symmetric diffusion of the population density. Hence, unequal entries imply that mutation results in asymmetric diffusion across trait space so that mutation affects some traits more than others. Additionally, if *M*_*ij* ≠_0 when *i* ≠ *j*, mutation contributes to covariance between traits *z*_*i*_ and *z*_*j*_. This model of mutation can be obtained from a diffusion-limit of an individual-based model that assumes independence of reproduction and mutation in which the phenotypic effect size of mutation is small (see Supplementary Material, Section 7). A model where mutation and reproduction interact has been studied by Wickman et al., (2023).

Mathematically, assuming *n* is finite allows us to write 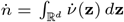. We can then apply integration-by-parts to obtain

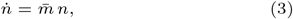

where 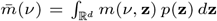 is mean fitness. Biologically, *n* should be very large since drift is being ignored. However, this approach can still be useful for gaining insights when *n* is small, which occurs for example with Lotka-Volterra models (Akjouj et al., 2024). A version of Equation (3) based on a finite number of genotypes (in place of continuous trait variation) is given in Crow and Kimura (1970) on page 10 above their equation 1.2.7.

Dynamics of the mean trait vector are obtained by applying the quotient rule and integration-by-parts to 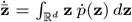, which provides

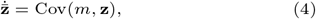

where Cov(*m*, **z**) is a *d*-dimensional vector with *i*-th entry given by

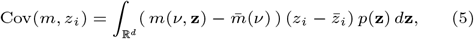

where 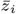 is the *i*-th entry of 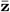. Equation (4) is similar to the Price equation without transmission bias in that evolutionary change is expressed as a covariance between fitness and trait value (Price, 1970, 1972). However, unlike the Price equation (which is a general identity) the present result is derived from an explicit population dynamical model and therefore depends on specific biological and analytical assumptions.

The same techniques can also be applied to obtain 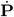 as

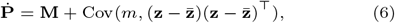

where 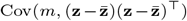 is a *d × d* matrix with *ij*-th entry given by Cov(*m*, 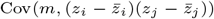). The response of the trait covariance matrix to selection as a covariance of fitness with 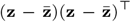 was previously found by Lande and Arnold (1983), without an explicit mutation term. An expression including **M** was later presented by Phillips and McGuigan (2006), but not derived from an underlying dynamical model. The steps required to arrive at equations (3)-(6) are detailed in the Supplementary Material, Section 2.

Equations (3), (4), and (6) follow from the underlying trait-structured dynamical model and make no assumptions on the form of the trait distribution. Consequently, the model can be used to study populations with non-trivial higher moments, such as skewed or otherwise non-Gaussian distributions. However, this flexibility of accommodating general trait distributions comes at the cost of a limited range of fitness functions that are amenable to study. This is due to i) the challenge of calculating covariances between arbitrary fitness functions and phenotypic moments, and ii) moment-closing issues that often arise during these calculations (Barton and Turelli, 1987; Gilpin and Feldman, 2019; Guerand et al., 2023). In spite of this limitation, a family of biologically important fitness functions are tractable to study using *DC*. These function take the form

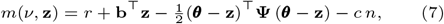

where **b** = (*b*_1_, …, *b*_*d*_)^⊤^ (which does not depend on **z**) confers directional selection (positive or negative for trait *z*_*i*_ depending on the sign of *b*_*i*_), and **Ψ** is a symmetric matrix that mediates stabilizing or disruptive selection around the vector ***θ***. Classically, **Ψ** is denoted ***ω***^−1^ in quantitative genetic theory (e.g., Lande, 1979; Jones et al., 2012). The absence of stabilizing selection under the classical notation occurs when all eigenvalues of ***ω*** diverge towards +∞. To avoid this technicality, I use **Ψ** so that the absence of stabilizing selection is associated with the zero matrix **Ψ** = **0**. Additionally, the parameter *c* ≥ 0 is the strength of competition, which here acts globally between individuals regardless of trait, and *r* is the growth rate in the absence of selection and competition (i.e., when **b, Ψ**, and *c* are all zero). It is possible to generalize this family so that competition is non-global, but this requires additional technical details as the fitness function becomes operator-valued (which is treated, for example, by Volpert, 2014; and by Etheridge et al., 2024). Supplementary Material, Section 3 connects these fitness functions to those known from classical discrete time quantitative genetic models.

Combining equations (3), (4), and (6) with Equation (7) provides (derivation given in Supplementary Material, Section 2.3)

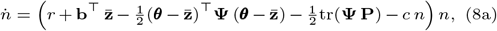

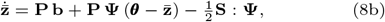

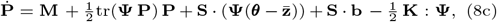

where **S** is the third-order skew tensor of **z** defined by 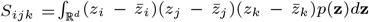. Products with **S** are given by (**S** : **Ψ**)_*i*_ = ∑_*jk*_ *S*_*jki*_Ψ_*jk*_, (**S** · **Ψ**)_*ij*_ = ∑_*k*_ *S*_*ijk*_Ψ_*kj*_, and (**S** · **b**)_*ij*_ = ∑_*k*_ *S*_*ijk*_*b*_*k*_. In addition, **K** is the fourth-order kurtosis tensor of **z** defined by 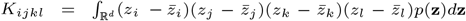. The product with **K** is given by (**K** : **Ψ**)_*ij*_ = ∑_*kl*_ *K*_*ijkl*_Ψ_*kl*_. Equation (8) collects the abundance, mean, and covariance dynamics (including skewness and kurtosis contributions) into a single system for clarity. While related formulations exist, this particular consolidation does not appear to be standard in the literature.

Equations (8) demonstrate that, for fitness functions taking the form of (7), the dynamics of abundance, mean trait, and trait covariance matrix depend on higher phenotypic moments for non-normal trait distributions. In fact, a complete description requires an infinite number of equations. However, by combining Equation (7) with multivariate DAGA (i.e., with Equation (1)), I demonstrate in Supplementary Material, Section 2.5 that, when **Ψ** is positive definite, *ν*(**z**) has a globally asymptotically stable equilibrium proportional to the density of a multivariate normal distribution with covariance matrix

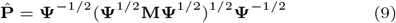

(with matrix square roots defined using eigenvalue decomposition: **Q** = **UΛU**^−1^ implies **Q**^1*/*2^ = **UΛ**^1*/*2^**U**^−1^) and equilibrium mean vector

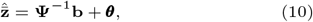

and with a total abundance

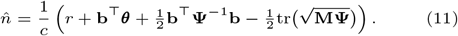

Derivations for these equilibrium expressions are given in Supplementary Material, Section 2.4.

The mutation-selection balance of phenotypic variance 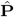 is a multivariate generalization of the univariate classical result obtained from multi-locus models (e.g., Bulmer, 1972) and continuum-of-alleles models (e.g., Bürger, 1986). Lande (1980) found a similar result using a multilocus approach that included recombination. In particular, 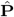 has the same matrix-square-root structure as the equilibrium found by Lande (1980), and the two coincide when linkage is negligible and when **M** corresponds to the sum of Lande’s locus-specific mutation matrices. Further work is needed to extend these results for more general fitness functions and higher phenotypic moments, which are already known to have important evolutionary consequences (Débarre et al., 2015). A new approach to study the dynamics of higher moments for univariate traits was recently introduced by Gilpin and Feldman (2019).

#### 2.1.2. The Deterministic Gradient Version (DG)

As noted above, deriving population dynamics from arbitrary fitness functions can be challenging in the more general framework based on covariances between fitness and phenotype. To overcome this we can make the useful simplifying assumption that traits follow a multivariate normal distribution. Under this assumption, all phenotypic moments (such as skew and kurtosis) can be expressed in terms of the mean and variance which greatly simplifies expressions such as equations (8). While deviations from normality can have consequences for ecological and evolutionary processes (Turelli, 1988; Débarre et al., 2015), normality has been an important initial assumption for studying a wide range of topics such as reaction-norm evolution (Lande, 2014), coevolving mutualistic networks (Nuismer et al., 2018), and niche construction (Fogarty and Wade, 2022). Furthermore, normality is a well-established approximation that holds under many genetic and selective scenarios (Turelli and Barton, 1994). Then, as a first pass, I assume traits are multivariate normally distributed for the rest of this paper.

In the context of this framework, the assumption of multivariate normality is particularly useful because it allows us to rewrite covariances between fitness and phenotype as gradients of fitness functions with respect to moments of the trait distribution. Such gradients can be analytically calculated for a broad range of fitness functions. The calculations to obtain these expressions begin with the definitions of covariances between fitness and phenotype, and then apply properties of the multivariate Gaussian function and integration-by-parts (see Supplementary Material, Section 2.3). As a result, the deterministic *DG* version of the framework is given by

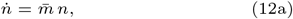

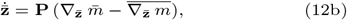

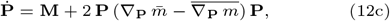

where 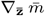 and 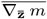 are *d*-dimensional vectors that respectively capture the effects of frequency independent and frequency dependent selection on mean trait evolution. More precisely, writing 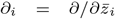 as the partial derivative operator with respect to the *i*-th mean trait, the *i*-th entry of 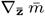 and 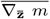 mare respectively given by 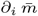 and 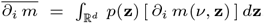. Similarly, 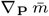 and 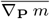 are *d × d* matrices that respectively quantify the dependence of fitness on trait variances and covariances. Writing ∂_*ij*_ = ∂*/*∂*P*_*ij*_ as the partial derivative operator with respect to the covariance between trait components *i* and *j*, the *ij*-th entries of 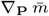 and 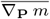 are respectively given by 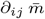 and 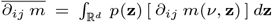. Although *m* is not written explicitly as a function of 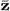 or **P**, it depends on them through the underlying trait distribution. The derivatives above are therefore understood as sensitivities of *m* to perturbations of the distribution that alter 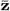 or **P**, and are not zero in general.

In Supplementary Material, Section 2.3, I show how to apply integration-by-parts to obtain the alternative expressions 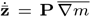 and 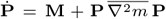 where (∇)_*i*_ = ∂*/*∂*z*_*i*_ (so 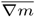 is the trait averaged gradient of fitness with respect to phenotype **z**) and 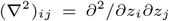 (so 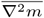 is the trait averaged Hessian of fitness with respect to phenotype **z**). A discrete-time counterpart to 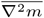 is given by equation (14b) in Lande and Arnold (1983).

The Hessian form for the response of **P** to selection is similar to previously known discrete-time expressions (Lande and Arnold, 1983; Débarre et al., 2014; Phillips and Arnold, 1989; Mullon and Lehmann, 2019). An important difference is that the discrete-time expressions include an additional term equal to the outer product of the per-generation response of the mean trait to selection, commonly denoted 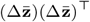, which appears with a negative sign in the discrete-time update of **P**. However, this extra term disappears under weak selection approximations often made in discrete-time quantitative genetic models (Lande, 1976, 1982; Turelli, 1984; Barton and Turelli, 1989; Turelli and Barton, 1990; Lynch and Walsh, 1998; Bürger, 2000; Walsh and Lynch, 2018). Furthermore, under a continuous-time scaling, the mean changes only *O*(Δ*t*) over a small interval, so its contribution to covariance is *O*(Δ*t*^2^). Dividing by Δ*t* to form a rate equation and letting Δ*t* → 0 removes this quadratic term, leaving only the curvature-driven (Hessian) part of selection in the continuous-time covariance dynamics, in agreement with equation (12c).

Mean trait and trait covariance dynamics can be expressed in index form as

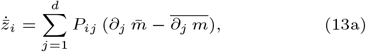

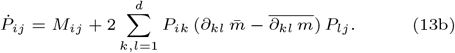

Inclusion of the terms 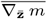 and 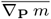 in equations (12b) and (12c) make it clear that this framework accounts for frequency-dependent selection. The decomposition of selection into components arising from frequency-dependent and frequency-independent sources has been made explicit in previous quantitative genetic treatments of mean trait dynamics (see eqn. (9) of Lande, 1976), but was only implicit in models of trait variance evolution due to their expressions in terms of Hessians (Lande and Arnold, 1983; Débarre et al., 2014; Phillips and Arnold, 1989; Mullon and Lehmann, 2019). A decomposition of the Hessian of fitness into frequency-dependent and frequency-independent components does not appear to be standard in existing treatments of trait-covariance evolution. Additionally, the approach outlined here can make a useful complement to adaptive dynamics and evolutionary game-theoretic treatments of frequency-dependent selection (Dieckmann and Law, 1996; Hofbauer and Sigmund, 1998; Traulsen et al., 2005).

An important caveat of both *DC* and *DG* versions of the framework taking the form presented above is that they rely on perfect heritability of trait values. However, traits are in general not perfectly heritable, and this is of fundamental importance in evolutionary biology. I therefore address this additional complexity in the following sub-subsection. I note here that extending *DC* and *DG* to include imperfect heritability results in nearly identical equations, the important difference is that average growth rates must additionally be averaged over a phenotypic residual term. I state this explicitly for the *DG* version below.

#### 2.1.3. Imperfect Inheritance

Following classical quantitative genetics, imperfect inheritance of trait values can be accounted for by assuming traits decompose into an additive genetic component and a residual component: **z** = **g** + **e** (Lynch and Walsh, 1998).

Biologically, this decomposition assumes that non-additive genetic effects and genotype-environment interactions are absent, so that **g** reflects only additive genetic effects, while **e** reflects the remaining non-heritable factors. This justifies interpreting **e** as the residual from fitting a linear statistical model of trait values to genetic predictors (Falconer and Mackay, 1996; Lynch and Walsh, 1998). The absence of non-additive genetic effects is especially important in this asexual framework, where the lack of recombination would otherwise allow multilocus allelic associations to render such effects heritable (Barton and Turelli, 1991). It is further assumed that the only trait-mediating factors that are heritable are additive genetic factors (i.e., cytoplasmic inheritance, persistent epigenetic marks, maternal effects, microbiome transmission, and cultural inheritance are all excluded). Under these assumptions, **e** is uncorrelated with **g** and is uncorrelated among individuals, consistent with interpreting **g** and **e** as the genetic predictor and residual in a linear model relating trait values to genetic variation.

In contrast to **e**, the additive genetic component **g** follows the same Gaussian mutation model described above. In particular, given that **g** is the additive genetic component of the trait of a parent, an offspring will have an additive genetic component that is multivariate normally distributed with mean **g** and covariance matrix **M**.

Assuming the residual **e** is independent and identically distributed for all individuals with mean zero and covariance matrix **E**, and denoting *γ*(**g**) the population density at genetic value **g**, and *ε*(**e**) the distribution of the residuals, the population density of trait values is given by 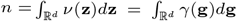. This preserves total abundance so that 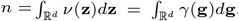. These assumptions also imply that the growth rate for the sub-population of individuals carrying genetic value **g** is given by 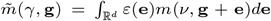. Furthermore, the fitness function for trait values *m*(*ν*, **z**) and the fitness function for additive genetic values 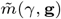 have the same mean value across the population

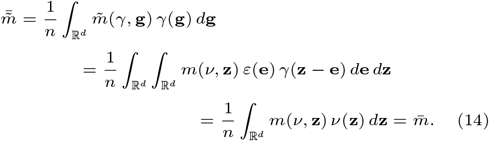

This approach to deriving the fitness associated with an additive genetic value given the fitness associated with a trait value and the distribution of residuals was originally introduced by Kimura and Crow (1978). It also appears in Equation (4) of Lande (1979). The dynamics of *γ* are given in analogy to 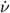 as

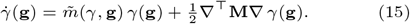

The distribution of additive genetic values in the population is given by 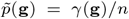. In combination with the model for phenotypic residuals, the mean trait vector is calculated as 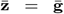 and the trait covariance matrix is **P** = **G** + **E**. To simplify calculating fitness, and to accommodate the assumption of multivariate normal traits, I assume *ε*(**e**) is the density of a multivariate normal distribution (with mean zero and covariance matrix **E**).

Under these assumptions the expression for abundance dynamics does not change, but the mean trait dynamics can be calculated as

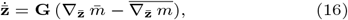

and **G**-matrix dynamics are given by

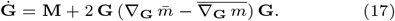

I apply this model of imperfect inheritance while describing the stochastic extensions of the framework below. Further details about how imperfect inheritance is interfaced with the stochastic extension of this framework, and how to obtain equations (16) and (17) are provided in the Supplementary Material, Section 5.

In summary, this section derives deterministic equations for abundance, mean traits, and trait covariances from the DAGA formulation (Equation (1)). It presents both the non-closed moment system for general trait distributions (equations (3)– (8)) and the closed gradient/Hessian form obtained under normality with imperfect inheritance (equations (12a), (16), and (17)), which the stochastic extensions build on below.

### 2.2. Stochastic Dynamics

In this section, I extend the framework to the case where demographic stochasticity induces random genetic drift. This extension introduces the stochastic version of the framework and builds on the Deterministic Gradient (*DG*) formulation developed above. Two closely related stochastic variants are presented.

The first variant, called the Brownian Motion Gradient version (*BG*), expresses stochastic dynamics with respect to Brownian motion processes, as is standard for stochastic differential equations (Øksendal, 2013; Evans, 2012). *BG* is particularly useful for numerical applications, and can be implemented using the Euler-Maruyama algorithm (Bayram et al., 2018). I illustrate this during a brief study of genetic correlations evolving in response to random genetic drift. Further information is provided in Supplementary Material, Section 1. Implementations of this approach using the DifferentialEquations.jl package in Julia (Rackauckas and Nie, 2017), and a manual implementation are provided at the github repository github.com/bobweek/multi-mtgl.

The second variant, called the Martingale Gradient version (*MG*), is useful for deriving the dynamics of more specific quantities. This use-case is illustrated below to formally obtain a stochastic differential equation tracking the evolution of genetic correlations in response to random genetic drift. To do so, I introduce a powerful set of novel heuristics (i.e., practical rules that allow exact calculations without requiring familiarity with the underlying measure-valued martingale theory, see Table 1) that can be used to derive an array of models from this variant of the framework. These heuristics are obtained by examining a so-called martingale process underlying this entire framework, which is also the namesake of *MG*. Mathematical details are provided in Section 4 of the Supplement.

Both stochastic extensions of the framework include an additional variable *v*, which is associated with the variance of lifetime reproductive output of individuals. Supplementary Material, Section 7.1 shows how to derive *v* from an individual-based model. This parameter can be interpreted as the population average of what Gillespie (1974; 1975; 1977) calls the variance in offspring number. Supplementary Material, Section 8 formally demonstrates this interpretation. Importantly, *v* can also be intrepreted as either the rate of reproduction for semelparous populations (demonstrated in Supplementary Material, Section 7) or the sum of birth and death rates in iteroparous populations (discussed in Week et al., 2021). I refer to *v* as the *reproductive variance*.

Previous work has shown that a novel form of selection can emerge when *v* depends on trait value, referred to as *noise-induced selection*, and for which evolutionary responses behave radically different from classical deterministic selection (Gillespie, 1974, 1977; Constable et al., 2016; Parsons et al., 2010; Kuosmanen et al., 2022; Bhat and Guttal, 2025). However, accounting for noise-induced selection leads to significantly more complex expressions for the evolution of mean trait vector and trait covariance matrix (e.g., see Week et al., 2021, for the univariate case). While genotype-dependent differences in the variance of offspring number have been documented in laboratory asexual systems (Turner and Chao, 1999), there is no evidence that such covariance is widespread, and selection acting explicitly on this variance component has not been directly measured in empirical populations. Then, as a first pass, I keep the expressions relatively simple by assuming *v* is constant across additive genetic values.

#### 2.2.1. The Brownian Motion Gradient Version (BG)

Accounting for the effects of demographic stochasticity, the abundance dynamics can now be expressed as the following stochastic differential equation

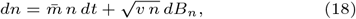

where the noise process *B*_*n*_ is a standard Brownian motion (i.e., *B*_*n*_(*t*) has variance equal to *t* and *B*_*n*_(0) = 0). Assuming multivariate normality, and the above model of imperfect inheritance, the mean trait dynamics (derived in Supplementary Material, Section 5.3) are given by

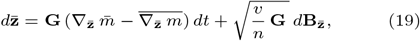

with **B**_**z**_ a *d*-dimensional vector of independent standard Brownian motions, and the matrix square root 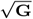 can be any *d × d* matrix **A** that satisfies **AA**^⊤^ = **G** (for examples, see Kessy et al., 2018). Equation (19) provides a continuous-time extension to the framework of multivariate evolution introduced by Lande (1979). Alternatively, the stochastic dynamics of mean traits can be expressed using index notation as

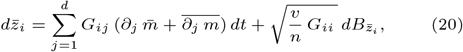

where 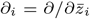 and, for each *i*, 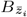 is a standard Brownian motion and non-independence for each *ij* is encoded by the heuristic 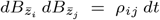, with 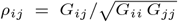 the genetic correlation between traits *i* and *j*. This heuristic is particularly useful when applying Itô’s formula (the stochastic analog of the chain rule, see Klebaner, 1998; Øksendal, 2013; or Evans, 2012) to derive dynamics for functions of mean trait values. This approach may be used, for example, to track the dynamics of interaction coefficients for coevolving species as a multivariate generalization of the approach taken by Week and Nuismer (2021).

In Equation (19) the correlated effects of genetic drift on mean trait evolution are encoded by the product 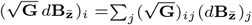. In contrast, for Equation (20), these correlated effects are encoded directly by the non-independence of the Brownian motions 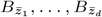. In particular, because 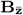 is defined such that its entries are independent, the *i*-th entry of 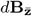 cannot be equal to 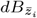 as 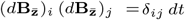, where *δ*_*ij*_ = 1 when *i* = *j* and zero otherwise. These heuristics work in the absence of multivariate normality, but the deterministic component of 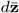 expressed above does depend on multivariate normality.

The expression of 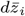 in Equation (20) using non-independent Brownian motions is not standard. However, this form will be useful below to illustrate the relationship between the *BM* (equations (18), (20) and (23)) and *MG* (equations (26)) versions of the framework.

Supplementary Material, Section 5.4 shows that the application of the multivariate normal approximation to the stochastic dynamics of the **G**-matrix leads to the matrix equation

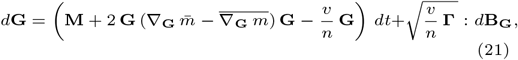

where **Γ** is a fourth-order tensor describing the covariance structure for the response of **G** to drift. Defining products of fourth-order tensors **X** and **Y** by (**X** : **Y**)_*ijkl*_ = ∑_*mn*_ **X**_*ijmn*_**Y**_*mnkl*_, we can write 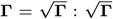. Furthermore, we have 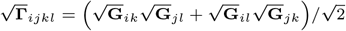, and 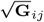 is the *ij*-th entry of 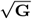 (as mentioned beneath Equation (19) above, 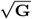 is any *d × d* matrix such that 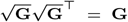). Importantly, this implies 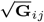 is *not* equal to 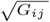.

The product 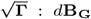 returns a *d × d* matrix with *ij*-th entry 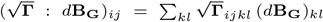. The *d × d* matrix-valued Brownian motion **B**_**G**_ has independent standard Brownian motions along its diagonal entries. The off-diagonal entries are standard Brownian motions scaled by one-half (i.e., the variance of (**B**_**G**_)_*ij*_ (*t*) is *t/*2 when *i ?*= *j*), and symmetric entries are equivalent so that (**B**_**G**_)_*ij*_ = (**B**_**G**_)_*ji*_. The covariance structure of **B**_**G**_ is summarized by the heuristic

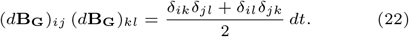

Scaling the off-diagonals by one-half ensures variances and covariances are correctly propagated while calculating the stochastic consequences of drift for **G**-matrix dynamics. Further information on symmetric normal matrices can be found in Gupta and Nagar (2018), particularly Theorem 2.5.1.

Unlike the expression for the stochastic component of 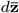, the stochastic component for *d***G** does depend on the assumption of a multivariate normal trait distribution.

Using index notation, the expression (21) simplifies to a *d*(*d* + 1)*/*2-dimensional system of equations summarized by

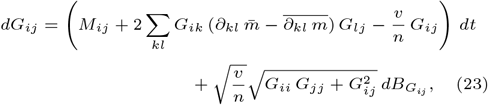

for 1 ≤ *i* ≤ *j* ≤ *d* where ∂_*ij*_ = ∂*/*∂*G*_*ij*_ and for each *ij* we have 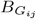 is a standard Brownian motion with 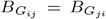. The covariance between 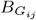 and 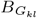 is encoded by the heuristic

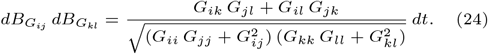

Importantly, this implies that 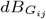 is *not* the *ij*-th entry of *d***B**_**G**_.

Finally, the noise process driving abundance in general does not covary with the noise processes driving mean traits and trait covariances (i.e., 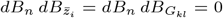), and, under the assumption of multivariate normality, the same holds for trait means and trait covariances (i.e., 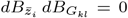). This agrees with the off-diagonal entries in equation (7b) of Barton (1989), but my results for the diagonal entries disagree. See Supplementary Material, Section 5.4 for more details.

The vector-matrix expression of *BG* (equations (19) and (21)) is particularly well-suited for numerical exploration of models because the covariance structure of the noise processes associated with random genetic drift are written explicitly in terms of sums involving the entries of the matrix square root 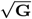. Hence, this version of the framework is easily interfaced with a common algorithm to numerically integrate systems of stochastic differential equations, the Euler-Maruyama method (Bayram et al., 2018). The numerical benefits of the *BG* version of the framework are highlighted in Section 3 below to study **G**-matrix evolution.

The expression of *BG* in index notation (equations 20 and 23), while arguably more complex in appearance, are useful for deriving analytical results, and especially when applying Itô’s formula to derive the dynamics of a quantity depending on 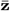 and/or **G** (such as the dynamics of growth rate, 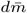, for instance). However, there are limitations with this application of *BG*, particularly for dealing with sums of stochastic differentials such as 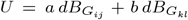. If the Brownian motions 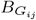 and 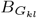 were independent, then we can write 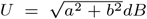 with *B* being another standard Brownian motion. Because 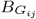 and 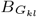 are not independent, which is captured by Equation (24) above, it is not obvious how to properly express *U* in terms of *a, b* and a single standard Brownian motion *B*.

To overcome these challenges, and also to simplify typical calculations, it is useful to rewrite the stochastic components of equations (20) and (23) in terms of an underlying stochastic process ℳ. Because ℳ satisfies a martingale property (roughly, this means 𝔼 [ℳ (*t* + *s*) | ℳ (*t*)] = ℳ (*t*) for all *s >* 0), I refer to this version of the framework as the Martingale Gradient version (or *MG* for short). In the following subsection, I present this version of the framework in a way that minimizes technical pre-requisites, while maintaining the same scope as equations (20) and (23) above. Additionally, heuristics are offered in Table 1 for making calculations, and these are demonstrated by deriving a stochastic differential equation for the correlation between two traits driven purely by random genetic drift.

#### 2.2.2. The Martingale Gradient Version (MG)

In Supplementary Material, Section 4, I show that this framework is justified based on a martingale process ℳ. Brownian motion is a special case of a martingale process, and stochastic equations are often expressed in terms of a Brownian stochastic differential *dB*. However, in this framework, stochastic equations can also be expressed in terms of the stochastic differential with respect to ℳ, denoted by *d*ℳ. This establishes a powerful approach for deriving fundamental insights into evolutionary processes. Further mathematical details are provided in Supplementary Material, Sections 4 and 5, but here I focus on pragmatic aspects regarding calculations that involve *d*ℳ.

I begin by artificially defining symbols related to *d*ℳ in terms of the Brownian motions that appear in equations (18), (19), and (21), and use these definitions to express the framework in terms of *d*ℳ. This is done purely for the sake of motivating the material that follows. After this, I introduce some properties of *d*ℳ and show how these can be used to recover the Brownian motions initially used in the artificial definitions mentioned above. I then provide general heuristics summarized in Table 1 for working with *d*ℳ, and illustrate these heuristics by deriving the response of trait correlations to random genetic drift.

For now, define the symbols 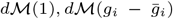 and 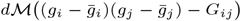 as follows:

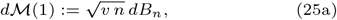

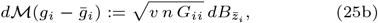

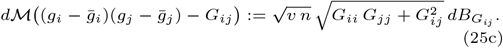

Using this notation, the *MG* version of the framework can be written as

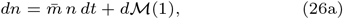

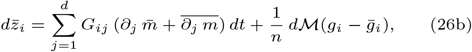

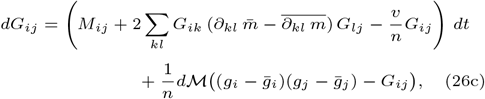

where recall that 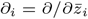 and ∂_*ij*_ = ∂*/*∂*G*_*ij*_.

The covariance structure of the system is maintained through heuristics (summarized in Table 1) for computing products of the above stochastic differentials *d*ℳ(*x*). To understand the heuristics, I introduce some useful notation for keeping track of averages across the distribution of traits in the population. Specifically, for functions *x*(**g**) and *y*(**g**), I define the symbols ∥ *x* ∥ and ⟨ *x, y* ⟩ as follows:

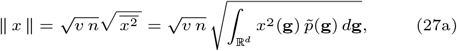

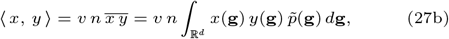

where 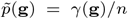 is the distribution of additive genetic values **g** in the population (assumed to be multivariate normal). To provide a few examples, one can calculate 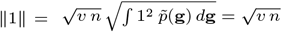 along with

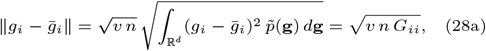

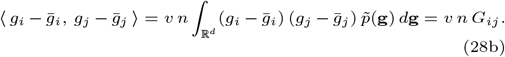

The martingale process ℳ mentioned above can be thought of as a mapping that associates functions with stochastic processes. This idea is formalized in Section 4 of the Supplement. In general, the process that ℳ associates with *x*, denoted ℳ(*x*), will not be a Brownian motion. However, Supplementary Material, Section 4.2 provides justification for what I call the *scaling property*, which states that

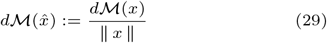

is the stochastic differential of a standard Brownian motion.

Using this heuristic, we can calculate

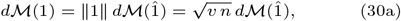

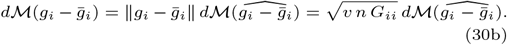

Comparing with equations (18) and (20), these calculations highlight the fact that 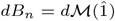 and 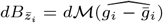. A similar equality holds for 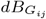, but for the sake of brevity this expression is omitted (see Supplementary Material, Section 5.4 for details).

Informally speaking, this scaling property allows us to “factor out” the standard deviation from the noise process driving the dynamics of a univariate function of the population (such as *p*, 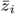, or *G*_*ij*_), and to replace that noise process with the product of the resulting standard deviation with a standard Brownian motion. This is the crucial step for obtaining equations (18), (20), and (23) from the underlying martingale process ℳ, as detailed in Supplementary Material, Section 5 where the heuristics summarized in Table 1 are applied to derive *MG* and *BG*. The calculations shown in Equation (30) are heuristic because they lead to accurate results without a rigorous understanding of their justification.

Just as 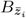 and 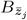 have a covariance encoded by the heuristic 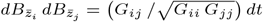, the processes ℳ(*x*) and ℳ(*y*) will also covary following a more general heuristic. More precisely, Supplementary Material, Section 4.2 justifies the *multiplicative property* which states that, for functions *x, y*, we have

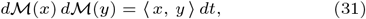

where ⟨ *x, x* ⟩ = ∥ *x* ∥^2^. In addition, we have *dt*^2^ = 0 and *dt d*ℳ(*x*) = *d*ℳ(*x*) *dt* = 0 for any function *x*, which are exact equalities in standard approaches to stochastic differential equations (see Øksendal, 2013; Evans, 2012; Klebaner, 1998). In this context, phenotypic moments *µ*_1_, *µ*_2_ (such as 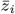 or *G*_*ij*_) behave as constants such that ⟨ *µ*_1_ *x, µ*_2_ *y* ⟩ = *µ*_1_ *µ*_2_ ⟨ *x, y* ⟩ (but see Supplementary Material, Section 4.2 for justification).

Finally, we also have the *additive property*: *d*ℳ(*a x* + *b y*) = *a d*ℳ(*x*) + *b d*ℳ(*y*), for functions *x, y* and constants *a, b* (more generally, *a, b* can also be phenotypic moments such as 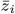 and *G*_*ij*_, see Section 4.2 of the Supplement). This property is particularly useful for computing the correct stochastic term for an equation resulting from the application of Itô’s formula, as illustrated in Section 3 below.

The heuristics associated with the *MG* version of the framework are summarized in Table 1. Example usage of these heuristics are provided through the derivation of this framework in Supplementary Material, Section 5. With these heuristics, it is straightforward to work with equations (26) to rigorously obtain the dynamics of quantities related to the population. For instance, the heuristics summarized in Table 1 may be used to formally derive a stochastic differential equation for the evolution of genetic correlations in response to drift. Indeed, I do this now.

## 3. Genetic Correlations and G-matrix Dynamics

In the following subsection I illustrate how to apply the above framework to obtain new models of evolutionary phenomena. Specifically, I apply the heuristics introduced in the *MG* version of the framework to derive the dynamics of genetic correlations responding only to random genetic drift. Using this derivation, I then provide biological insights into the consequences of drift for genetic correlations in asexually reproducing populations. In section 3.2, I then discuss the significance of these results in the context of empirical research on **G**-matrix dynamics.

### 3.1. Consequences of Drift for Trait Correlations

The additive genetic correlation between traits *z*_*i*_ and *z*_*j*_ is given by 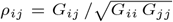. For this subsection, I focus on the single correlation between *z*_*i*_ and *z*_*j*_, and simply write this as *ρ*. To focus on the effects of random genetic drift, and for the sake of simplicity, I ignore mutation, selection, and abundance dynamics in this analysis. For the sake of illustration, I outline only the steps needed to demonstrate how the framework is applied and to obtain the biological result. The full derivation is provided in Supplementary Material, Section 6.1.

Because we have the stochastic equations for *dG*_*ij*_, *dG*_*ii*_, and *dG*_*jj*_ (equation 26c), and because *ρ* can be thought of as a function 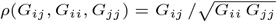, we can apply Itô’s formula, which, in this context, states the following stochastic analog of the chain-rule: *dρ* = *K* + *L*, with *K* = (∂_*ij*_ *ρ*) *dG*_*ij*_ + (∂_*ii*_*ρ*) *dG*_*ii*_ + (∂_*jj*_ *ρ*) *dG*_*jj*_ and

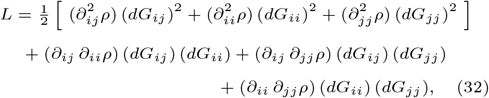

where ∂_*ij*_ *ρ* is the partial derivative of *ρ* with respect to *G*_*ij*_.

With this formula in hand, all that is left is to carry out calculations based on the heuristics from the *MG* version of the framework. First off, we can directly compute the *deterministic* component of the sum *K* and find that this cancels to zero. This part of the calculation does not require the heuristics for *d*ℳ introduced above. However, by using equation (26c) in the expression for *K*, and by leveraging the additive property of *d*ℳ, the *stochastic* component of *K* can be rewritten as

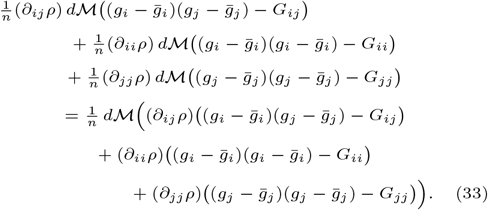

Writing the argument to *d*ℳ on the right-hand-side of Equation (33) as *H*_*ij*_, the scaling property shows the stochastic component of *K* can be written as

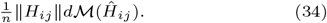

Calculation of ∥*H*_*ij*_ ∥ only requires computation of the first order derivatives ∂_*ij*_ *ρ*, ∂_*ii*_*ρ*, ∂_*jj*_ *ρ* and the fact from multivariate normal distributions that

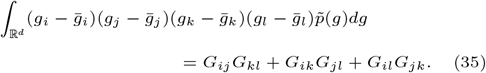

Working through these calculations provides

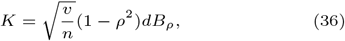

where *B*_*ρ*_ := ℳ(*Ĥ*_*ij*_) is a scalar-valued standard Brownian motion.

To compute *L*, I first rewrite the product (*dG*_*ij*_) (*dG*_*kl*_) using the notation introduced in the *MG* version of the framework above (i.e., using equations (26)), which provides

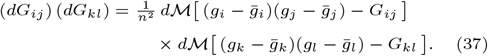

Applying the multiplicative property for *d*ℳ, the product simplifies to

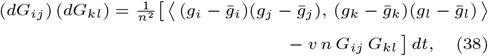

where I made use of 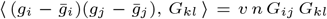. Again applying property (35) from above, I obtain the further simplification 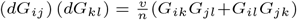. Interfacing this result with derivatives of *ρ*(*G*_*ij*_, *G*_*ii*_, *G*_*jj*_), I arrive at

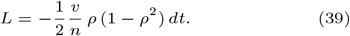

Hence, by applying the *MG* version of the framework, I find that trait correlations driven entirely by drift follow the ordinary stochastic differential equation

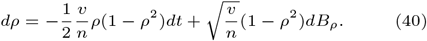

Equation (40) shows that the effect of drift is mediated by the ratio of the variance of lifetime reproductive output *v* (typically set to *v* = 1 in classical quantitative genetic models such as Lande, 1976) to the effective population size *n*. Additionally, one can check that (because mutation and selection are absent) *ρ* = *±*1 are stationary points. Furthermore, leveraging the fact that Equation (40) defines a one-dimensional diffusion, we can in principle solve for its stationary distribution 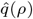 (Karlin and Taylor, 1981; Etheridge, 2010), which is done in Supplementary Material, Section 6.2. However, in attempt to do so we arrive at the non-integrable function:

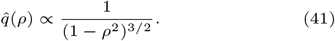

The lack of a formal stationary distribution creates a challenge for understanding the limiting behavior of genetic correlations evolving solely in response to drift. This can be partially overcome by again leveraging the theory of one-dimensional diffusions to demonstrate that the boundaries *±*1 are both attracting and unattainable (Karlin and Taylor, 1981). That is, genetic correlations tend towards their extremes, but never fix at *±*1. This is visualized in Figure 1 by plotting the distribution *q*(*ρ*) of genetic correlations as numerical solutions to the forwards Kolmogorov (i.e., Fokker-Planck) equation associated with (40). Biologically, |*ρ*| increases because drift randomly samples finite numbers of individuals, thereby causing transient correlations among the additive genetic values these individuals carry. Further biological implications are discussed in the following section.

**Fig. 1.**
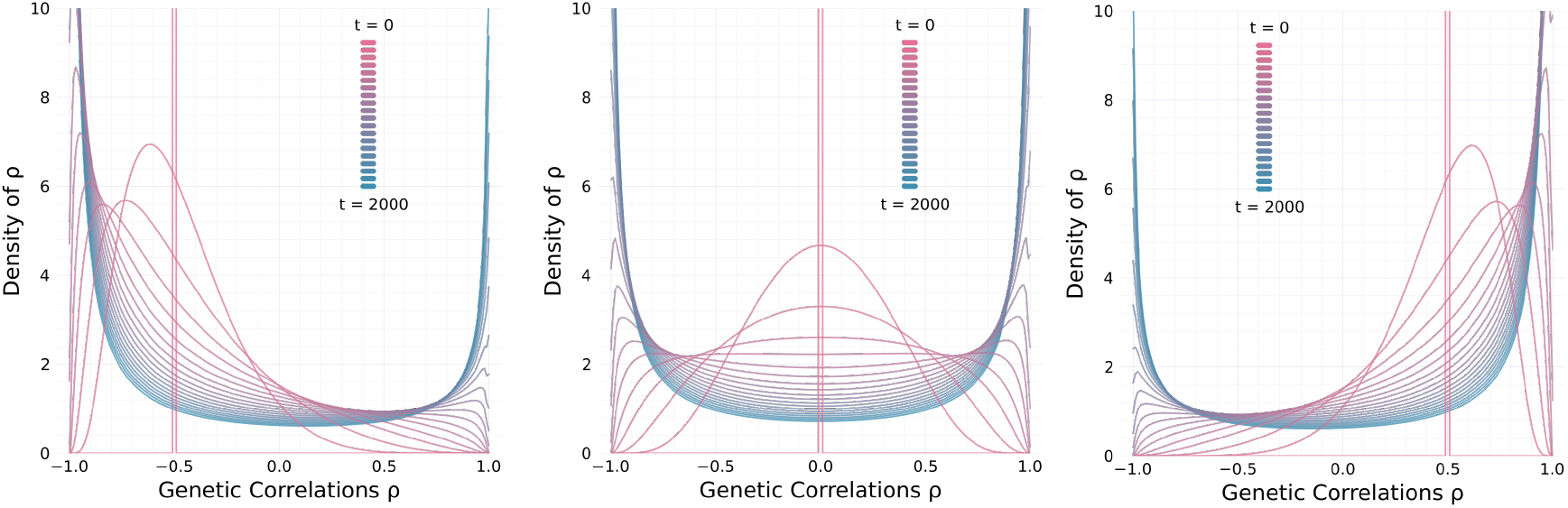
The distribution of genetic correlations evolving under drift alone converges to a non-integrable stationary solution. Shown here are dynamics for initial distributions taking approximate point masses at *ρ*_0_ = −0.5 (left panel), *ρ*_0_ = 0.0 (middle panel), and *ρ*_0_ = 0.5 (right panel). Solutions at earlier times are colored green, and later times are purple which run until *t* = 2000. The rate of drift is set to *v/n* = 0.001.

Another way to view this result is by applying Itô’s formula to *u* = tanh^−1^(*ρ*) (detailed calculations given in Supplementary Material, Section 6.3). Doing so returns

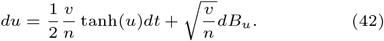

Because *ρ* = tanh(*u*) is a monotone increasing function of *u*, statements about *u* map directly to *ρ*. In particular, *u* = *ρ* = 0 is unstable because tanh(*u*) is positive for positive *u*, and negative for negative *u*. Additionally, if *u* is much greater than 1, then 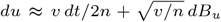 and, writing this point as *u*_0_, we have 𝔼 [*u*_*t*_] ≈ *u*_0_ + *v t/*2*n*. The analogous approximation holds when *u* is much less than −1. Justification for these claims is made in Supplementary Material, Section 6.3. This agrees with the boundary classification result above that demonstrates drift has an overall tendency to drive trait correlations towards *±*1. Numerical estimates for sample paths of the solution to (40), illustrated by Figure 2, support this conclusion.

**Fig. 2.**
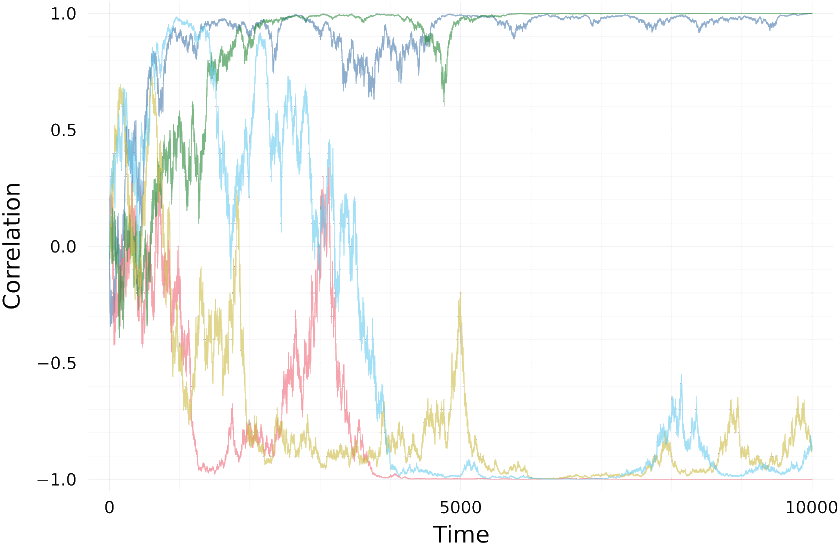
Drift drives trait correlations towards *±*1. Shown here are five replicates illustrating the path-behavior of trait correlations following Equation (40). Each replicate is initiated with *ρ*_0_ = 0 and the rate of drift is set to *v/n* = 0.001.

To confirm the heuristics return correct expressions, I also simulated the evolution of the **G**-matrix driven solely by random genetic drift for a 2-dimensional trait by applying the Euler-Maruyama method (Bayram et al., 2018) to the *BG* version of the framework. I then back-calculated the genetic correlation *ρ* based on the simulated time-series of **G**. Figure 1 in Supplementary Material, Section 1 demonstrates this approach agrees with Equation (40). Further information on the numerical implementation is given in Supplementary Material, Section 1, and associated code is available at the github repository github.com/bobweek/multi-mtgl.

### 3.2. Evolution of The G-Matrix in Response to Drift

The above result demonstrates that drift increases correlations between traits, at least for asexually reproducing populations where recombination is absent. This perspective offers an important refinement of the conventional wisdom that **G**-matrices responding to drift should merely contract proportionally (Lande, 1979; Phillips and McGuigan, 2006; Dugand et al., 2021; Mallard et al., 2023a). In particular, it is classically thought that the orientation of the **G**-matrix should not change due to drift on average, and hence any changes in orientation may be a sign of selection (Roff, 2000; Steppan et al., 2002; Cano et al., 2004; Chapuis et al., 2008; Aguirre et al., 2013).

This idea can be obtained from Equation (21) if the stochastic component is neglected. Then, the deterministic response due to drift is described by **Ġ** = −*v* **G***/n*, which has the solution **G**_*t*_ = **G**_0_*e*^−*vt/n*^, and is a continuous-time equivalent of the result found on page 409 in a paper by Lande (1979). Numerical results displayed in Figure 2 of Supplementary Material, Section 1 indicate agreement with this classical scaling result in expectation. However, when stochastic fluctuations dominate the dynamics over the timescale of interest (so that individual sample paths diverge strongly despite a smooth deterministic trend) the average response to drift may provide little information for within-population dynamics.

Taking an experimental approach, Phillips et al., (2001) established isolated populations of *Drosophila melanogaster* from a common base population and found significant variation across replicates for the response of **G** to drift. This occurs in spite of the fact that *Drosophila melanogaster* is a sexually reproducing organism, and recombination should act to break up correlations caused by linkage (discussed further below) and thereby reduce variation across replicates. Figure 3 emphasizes these variable outcomes theoretically by superimposing expected trait correlations over a collection of individual outcomes. The expected correlations are approximated by averaging over replicates, but it took a very large number of replicates (*>*1000) to obtain a satisfactory deterministic trend. Because population size enters only through the ratio *v/n*, increasing *n* rescales the time axis but does not reduce pathwise variability or improve the predictive power of the average trend. The averages shift towards zero, in the sense of a slow directional trend that is not resolved to convergence over the finite time window shown, but we can see individual replicates are not predicted by this trend. Hence, to gain a more accurate picture of **G**-matrix evolution, there is a need to understand the path behavior of individual outcomes.

**Fig. 3.**
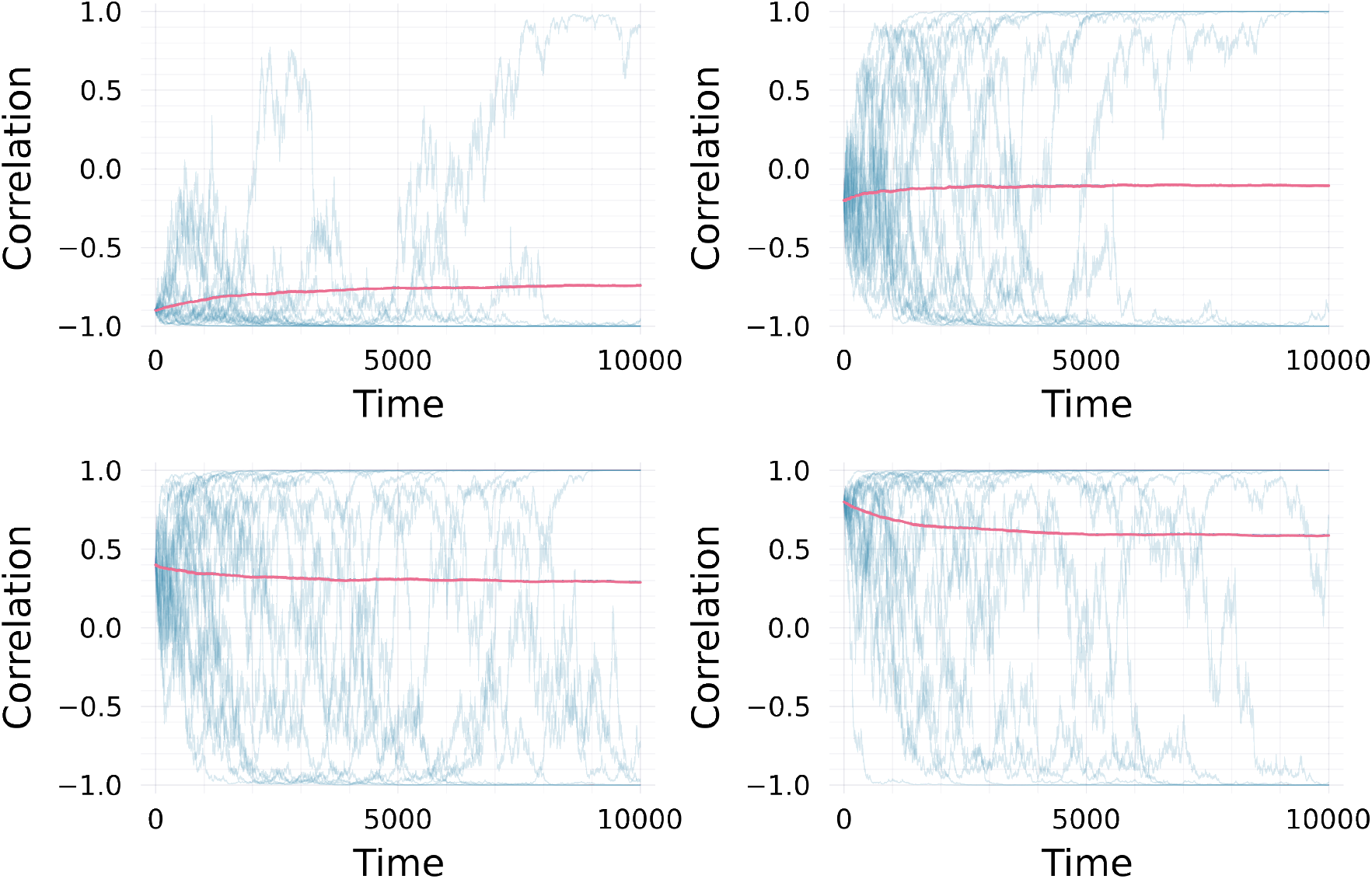
Averages over replicated time-series of trait correlation do not predict within population trait correlations. Averages were computed using 3000 replicates with initial correlations *ρ*_0_ = −0.9 (top-left), *ρ*_0_ = −0.2 (top-right), *ρ*_0_ = 0.4 (bottom-left), and *ρ*_0_ = 0.8 (bottom-right). For the sake of clarity, just 20 random replicates are shown behind the average trends.

One approach to gain insight for the trait correlation of a given replicate, as opposed to the average, is to analyze the proportion of time that correlations spend near *±*1 over the total time the process is running. Figure 4 plots the proportion of time that the magnitude of the trait correlation *ρ* spends above the threshold 1 − *ε*, i.e. with |*ρ*| *>* 1 − *ε*, averaged over 10 replicate trajectories with initial value *ρ*_0_ = 0 and *v/n* = 0.001, for *ε* ∈ {0.20, 0.10, 0.05, 0.02}. Smaller *ε* corresponds to increasingly strict neighborhoods of *±*1. This result shows that trait correlations typically aggregate near *±*1 in drifting, isolated, asexual populations with no mutational input. Hence, interpreting this as a signature of drift, we can say that if trait correlations in a population exhibit significant variation away from *±*1, then either the population has been drifting for only a short time relative to its effective population size, or other processes not captured by this model must be at play, such as ongoing mutation or recombination, which are explicitly excluded here. I therefore discuss mechanisms maintaining genetic correlations before concluding this section.

**Fig. 4.**
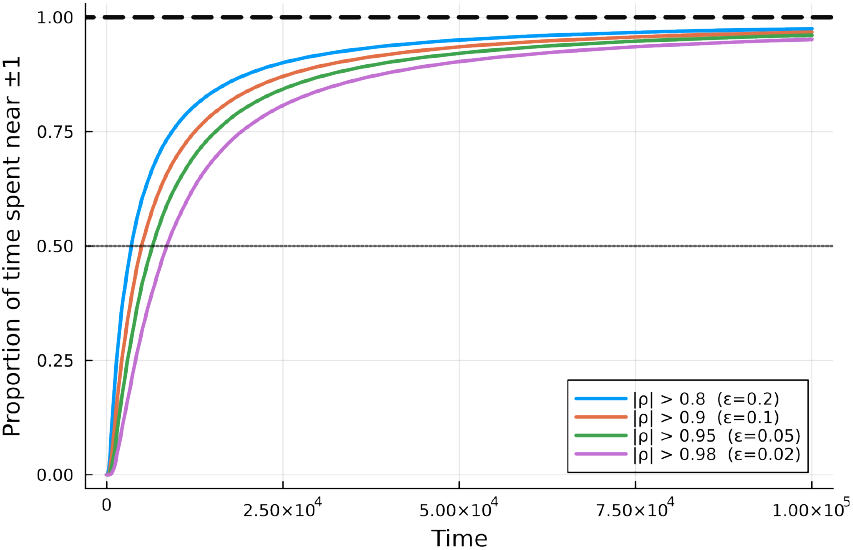
Trait correlations (*ρ*) within populations rapidly evolve towards *±*1 under random genetic drift. Shown is the proportion of time that |*ρ*| spends above a threshold 1 − *ε*, averaged over 10 replicate trajectories, with *ε* ∈ {0.20, 0.10, 0.05, 0.02}. Smaller *ε* corresponds to increasingly strict neighborhoods of *±*1. Simulations start from *ρ*_0_ = 0 with *v/n* = 0.001.

Pleiotropic loci and genetic linkage are two mechanisms that maintain genetic correlations (Lande, 1980). In this framework, pleiotropic mutations would arise from non-zero off-diagonals of the mutation matrix **M**. In contrast, drift facilitates chance correlations of additive genetic values between traits across individuals. Hence, this second mode of genetic correlation accumulation is directly analogous to the build-up of linkage by drift (Hill and Robertson, 1966; Ohta and Kimura, 1969; Lucek and Willi, 2021). For sexually reproducing populations, it is expected that genetic correlations should be maintained by pleiotropic loci as recombination breaks up linkage (Lande, 1980; Jones et al., 2003; Phillips and McGuigan, 2006). However, the results in this section show that drift alone drives genetic correlations toward extreme values. This raises the natural question of how this tendency would be counteracted by recombination, and whether a drift-recombination balance analogous to that studied in classical population genetics (e.g., Ohta and Kimura, 1969) would emerge.

The drift-only example considered here is intended as an illustrative application of the framework rather than a quantitative fit to existing experiments. In a clonal, mutation-free setting, the model predicts that the stochastic dynamics of genetic correlations depend on time only through the rescaled variable *τ* = (*v/n*)*t*. Equivalently, for populations with different effective sizes *n*, the distributions of correlation trajectories become comparable after rescaling time by *n/v*, so that increasing population size slows the dynamics without altering their qualitative path behavior. This prediction can be tested empirically using pathwise statistics such as the distribution of first-passage times *τ*_*c*_ = inf{*t* : |*ρ*(*t*)| *> c*} to a high-correlation threshold *c*, or the long-run fraction of time that correlations satisfy |*ρ*| *> c*; both quantities are predicted to scale with *n/v*. Such tests would, in principle, be feasible in replicated clonal systems with controlled demography and minimal mutational input, including bottlenecked microbial populations (e.g., *Escherichia coli* or asexually propagated *Saccharomyces cerevisiae*) or clonal macrophytes such as duckweed (*Lemna* or *Spirodela*) maintained in controlled microcosms. Extending the framework to incorporate sexual reproduction and recombination would be a natural next step for making quantitative predictions about drift-recombination balance. Nevertheless, the present results highlight that focusing solely on average, deterministic responses can obscure substantial pathwise structure, suggesting that conventional perspectives on **G**-matrix evolution merit closer examination.

## 4. Discussion

The classical quantitative genetic approach has stressed the importance of considering the genetic architecture of traits for understanding evolutionary processes, and has placed particular emphasis on the use of multi-locus models (Bulmer, 1972; Chakraborty and Nei, 1982; Slatkin, 1987; Turelli and Barton, 1994; Barton and Turelli, 2004; Barton et al., 2017; Walsh and Lynch, 2018). The advantage of this approach is its clear connection with explicit genetic details. However, its primary challenge is the manipulation of complex expressions that emerge at such level of detail. To overcome this, and establish a flexible tool for modelling the integrated ecological and evolutionary dynamics of populations carrying multivariate traits, I struck a balance between incorporating genetic detail and analytical tractability. As a consequence, this framework has potential for wide-spread use across topics in evolution, ecology, and population biology.

A central feature of this framework is that a wide range of classical models can be obtained by choosing an appropriate fitness function *m*. For example, stochastic Lotka-Volterra dynamics for a community of species is obtained from this framework by applying the growth rate *m*_*i*_ = *r*_*i*_ + ∑ _*j*_ *α*_*ij*_ *n*_*j*_ for species *i*, with *n*_*j*_ being the abundance of species *j* and *α*_*ij*_ the interaction coefficient. The long time behavior of the resulting stochastic Lotka-Volterra process for two species has been studied by Cattiaux and Méléard (2009). In addition, by making the interaction coefficients *α*_*ij*_ dependent on multivariate trait values **z**_*i*_, **z**_*j*_, models integrating coevolution with abundance feedbacks can be obtained similar to those studied by Gokhale et al., (2013); Cortez and Weitz (2014); and Patel et al., (2018). As a special case, continuous time analogs of multivariate coevolution models are obtained by focusing on two species with fixed (or infinite) abundances, and assuming *α*_*ij*_ (**z**_*i*_, **z**_*j*_) depends on the Euclidean distance between **z**_*i*_ and **z**_*j*_ (Gilman et al., 2012; Débarre et al., 2014). Klausmeier et al. (2020) studied models of evolutionary rescue with univariate traits using growth rates of the forms *m*(*z, t*) = *r* − *ψ*(*θ*(*t*) − *z*)^2^*/*2 and *m*(*z, t*) = *r* + *r*_0_*e*^−*ψ*(*θ*(*t*)−*z*) 2*/*2^, where *θ*(*t*) is a dynamic phenotypic optimum and *ψ* is the strength of stabilizing selection. Applying multivariate generalizations of these growth rates to the above framework leads to extensions of an evolutionary rescue model involving demographic stochasticity studied by Xu et al. (2023). Additionally, Jones et al. (2012) also studied a model of phenotypic adaptation to a dynamic optimum, but in the context of **G**-matrix evolution. This framework can then be used as a bridge between research topics such as evolutionary rescue and **G**-matrix evolution. This list provides a small set of examples for how this framework can be used to derive new models across a broad range of topics in ecology, evolution, and population biology.

Further work is needed to extend this framework in several directions. For instance, it is possible to incorporate sexual reproduction and recombination by assuming each trait value is determined by the sum of additive genetic effects contributed by two gametes, rather than by a single genetic value. In this case, each gamete contributes an additive genetic effect *g*_*i*_, and the trait is encoded as 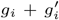, where *g*_*i*_ and 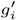 arise from convex combinations of the parental gametic values, similar to the approach taken by Lande (1980). How to formalize this extension using measure-valued processes is not obvious. In another direction, environmental stochasticity (i.e., stochastic growth rates) can be formally incorporated in a measure-valued context following the work of Mytnik (1996). Extending the framework in this direction may yield important additional insights complementing the work of Lande (2007; 2008), who studied the consequences of environmental stochasticity for long-run population growth rates. An extension in this direction may also be useful for studies investigating the consequences of environmental stochasticity on **G**-matrix evolution (e.g., Engen and Sæther, 2024).

I also point to two possible extensions that concern the role of development. The first accounts for the evolution of the mutational variance-covariance matrix **M** within the classical quantitative genetic framework. Allowing **M** to evolve enables analytical and simulation models for the evolution of evolvability, as studied by Jones et al. (2007), including numerical integration of the resulting stochastic differential equations (e.g. via Euler-Maruyama), rather than relying exclusively on individual-based models, and may yield more precise quantitative predictions amenable to experimental study (Mallard et al., 2023a). A second, conceptually distinct extension would be to replace the linear, regression-based genotype-phenotype map and residual model used here with nonlinear or mechanistic developmental maps, as in evo-devo frameworks (e.g., Milocco and Salazar-Ciudad, 2022; González-Forero, 2022, 2024b,a), in which genetic and phenotypic covariation emerge from explicit developmental dynamics rather than being parameterized through a mutational covariance matrix.

The application of this framework to study **G**-matrix evolution uncovered a more nuanced picture for the role of drift than what conventional wisdom suggests (Phillips and McGuigan, 2006). In particular, although drift indeed proportionally reduces **G**-matrices when averaged over many replicated populations, it also drives trait correlations towards their extremes within populations. This agrees with the result that the expected eigenvalues of **G** differ from the eigenvalues of the expectation of **G** under drift (Griswold et al., 2007). In addition, the impact of drift on genetic correlations can be understood as a reduction in the effective dimensionality of **G**-matrices (Hine and Blows, 2006), reflecting an extension of the principle that drift erodes heritable variation. That is, because drift drives pairwise correlations to *±*1, the distribution of multivariate traits in a population that has been evolving solely under drift may be well approximated after projecting onto a lower-dimensional trait space. Conceptually, this is similar to dimensionality reduction techniques such as principal components analysis (Kirkpatrick and Meyer, 2004). Importantly, this effect of drift on trait correlations calls into question comparative quantitative genetic methods utilizing **G**-matrices which routinely treat differences in **G**-orientation as a signal of selection in diverged populations (Roff, 2000; Steppan et al., 2002; Phillips and McGuigan, 2006; Cano et al., 2004; McGuigan, 2006; Chapuis et al., 2008; Aguirre et al., 2013; Dugand et al., 2021; Mallard et al., 2023a). These findings have broad implications for evolutionary biology, highlighting the need for revised theoretical perspectives and further empirical validation.

The study of genetic correlations and **G**-matrix dynamics using this framework can be extended by studying equation with mutation and selection. In the special case of one-dimensional traits, stabilizing selection, and no abundance dynamics, the theory of one-dimensional diffusions (Etheridge, 2010) can be used to show that the stationary distribution of additive genetic variance follows a generalized inverse Gaussian distribution (Jorgensen, 2012). This suggests the stationary distribution of **G** may follow a matrix-variate generalization, such as the matrix generalized inverse Gaussian (MGIG) distribution (Fazayeli and Banerjee, 2016). Proposed distributions may be checked by evaluating the forwards Kolmogorov equation associated with Equation (21) at equilibrium assuming a solution that follows the density of the proposal. Identification of the stationary distribution for (21) may then be used to study the distribution of genetic correlations maintained by interactions between mutation, selection, and random genetic drift.

In summary, this work introduces a versatile framework for modeling the stochastic eco-evolutionary dynamics of multivariate traits, providing a unifying approach that integrates mutation, selection, demographic stochasticity, and drift. By balancing mathematical rigor with accessibility, this framework enables the derivation of new models across a broad spectrum of population biology, making it a valuable tool for both theoretical and applied researchers. With its broad applicability, the framework presented here offers a foundation for future studies investigating the dynamics of populations in both theoretical and empirical contexts.

## 5. Acknowledgements

I thank Jonas Wickman, Hinrich Schulenburg, Brendan J.M. Bohannan, Arne Traulsen, Patrick C. Phillips, Peter L. Ralph, and two anonymous reviewers for their insightful feedback, which has greatly improved this manuscript. Special thanks to Stephen M. Krone, who taught me probability and inspired me to think about populations as random Hilbert spaces. This work is dedicated in honor of his retirement. Additionally, I am thankful for support through KiTE (i.e., Kiel Training for Excellence), which is part of the European Union’s Horizon Europe research and innovation programme under the Marie Sklodowska-Curie grant agreement number No 101081480.

