## Supplementary Material for "STOCHASTIC ECO-EVOLUTIONARY DYNAMICS OF MULTIVARIATE TRAITS: A Framework for Modeling Population Processes Illustrated by the Study of Drifting G-Matrices"

-

Bob Week

#### 1. Numerical Implementation

In the associated github repository (`https://github.com/bobweek/multi-mtgl`), I provide four different numerical implementations for the Brownian Motion Gradient version of the framework described in the main text. Each implementation has been written in Julia. The first three implementations (found in the folders labelled `knw`, `dff`, and `cov`) make use of the DifferentialEquations.jl package (Rackauckas and Nie, 2017). The first implementation, `knw`, works when analytical expressions for mean fitness $\bar{m}$ and fitness gradients $\nabla_{\bar{\mathbf{z}}}\bar{m}$, $\overline{\nabla_{\mathbf{z}} m}$, $\nabla_{\mathbf{G}}\bar{m}$, and $\overline{\nabla_{\mathbf{G}} m}$ have been identified. The second implementation, `diff`, works when an analytical expression for mean fitness has been identified, but the gradients $\nabla_{\bar{\mathbf{z}}}\bar{m}$, $\nabla_{\mathbf{G}}\bar{m}$ are numerically approximated using finite differences. This second implementation assumes frequency-independent selection so that $\overline{\nabla_{\mathbf{z}} m} = \overline{\nabla_{\mathbf{G}} m} = 0$. The third implementation, `cov`, works with just the fitness function $m(\nu, \mathbf{z})$ and numerically approximates integrals using a cubature method to obtain mean fitness and fitness gradients (which are computed using their expressions as covariances, see section 3.2 above). The fourth implementation (found in the folder labelled `diy`) is a manual implementation, which is described in the following paragraphs:

A simple approach to numerically solve stochastic differential equations is the Euler-Maruyama method (Bayram et al., 2018). This method is a natural stochastic extension of the Euler method to solving deterministic differential equations. In particular, for the $d$-dimensional differential equation $\dot{\mathbf{u}} = f(\mathbf{u})$ with $f : \mathbb{R}^d \to \mathbb{R}^d$, the Euler method approximates the solution with the recursion $\mathbf{u}_{t+\Delta t} = \mathbf{u}_t + f(\mathbf{u}_t)\,\Delta t$, for a given fixed time-step $\Delta t$. As an extension, the Euler-Maruyama approach to solving

$$d\mathbf{u} = f(\mathbf{u})\,dt + g(\mathbf{u})\,d\mathbf{B}, \tag{1}$$

with $g : \mathbb{R}^d \to \mathbb{R}^{d\times d}$, is given by the stochastic recursion

$$\mathbf{u}_{t+\Delta t} = \mathbf{u}_t + f(\mathbf{u}_t)\,\Delta t + g(\mathbf{u}_t)\,\mathbf{W}_t \tag{2}$$

where $\mathbf{W}_t$ is a $d$-dimensional multivariate normal random vector with independent entries each having mean zero and variance $\Delta t$, and the product $g(\mathbf{u}_t)\,\mathbf{W}_t$ returns another $d$-dimensional vector defined as usual matrix-vector multiplication

To numerically integrate $\mathbf{G}$-matrix dynamics, this approach is extended for matrix-valued recursions. In this case, we have $\mathbf{u}_t \in \mathbb{R}^{d\times d}$, $f : \mathbb{R}^{d\times d} \to \mathbb{R}^{d\times d}$ $g : \mathbb{R}^{d\times d} \to \mathbb{R}^{d\times d\times d\times d}$, and $\mathbf{W}_t$ a $d \times d$ symmetric normal matrix. In particular, $\mathbf{W}_t$ has independent diagonal entries each with mean zero and variance $\Delta t$, off-diagonals have mean zero and variance $\Delta t/2$, and symmetric pairs are perfectly correlated so that $[\mathbf{W}_t]_{ij} = [\mathbf{W}_t]_{ji}$, but otherwise entries are independent. The product $g(\mathbf{u}_t)\,\mathbf{W}_t$ is replaced by $g(\mathbf{u}_t) : \mathbf{W}_t$, and returns a $d \times d$ matrix following the tensor products described in the main text:

$$\left[g(\mathbf{u}_t) : \mathbf{W}_t\right]_{ij} = \sum_{kl} \left[g(\mathbf{u}_t)\right]_{ijkl} \left[\mathbf{W}_t\right]_{kl}. \tag{3}$$

##### 1.1. Comparing Results Based on Heuristics with Classical Scaling Result

Using the manual implementation described above, I first solved for $\mathbf{G}$ and then back-calculated $\rho$ to check that the solutions match those found using the heuristics presented in the main text to obtain the sde $d\rho$ (i.e., eqn (37)). Results indicating confirmation based on 500 replicates for initial values $\rho_0 = -0.5, 0.0, 0.5$ and rate $v/n = 0.1$ are displayed in Figure 1.

I also used this approach to compare the expected dynamics of $\mathbf{G}$-matrices under drift with the classical scaling result $\mathbb{E}[\mathbf{G}_t] = \mathbf{G}_0 e^{-v\,t/n}$. Using 500 replicates with initial value $G_{11} = G_{22} = 1.0$, $G_{12} = 0.5$ and rate $v/n = 0.1$, I calculated approximate expectations by averaging over replicate runs $(\langle G_{11}\rangle, \langle G_{22}\rangle, \langle G_{12}\rangle)$. Figure 2 demonstrates agreement.

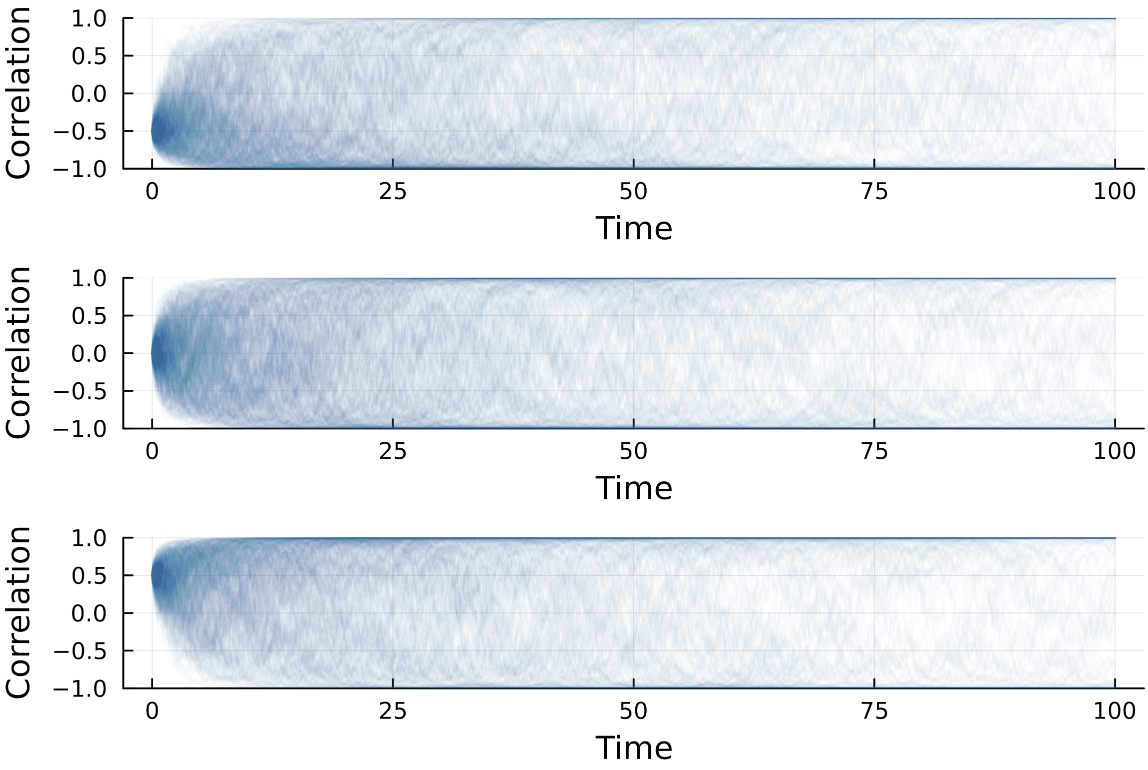


**Fig. 1.** Correlation dynamics obtained by back-calculuting $\rho$ from Euler-Maruyama solutions to $\mathbf{G}$ indicate agreement with heuristics presented in main text. Each panel displays 500 replicates initiated at $\rho_0 = -0.5$ (top), $\rho_0 = 0.0$ (middle), and $\rho_0 = 0.5$ (bottom) with rate $v/n = 0.1$.

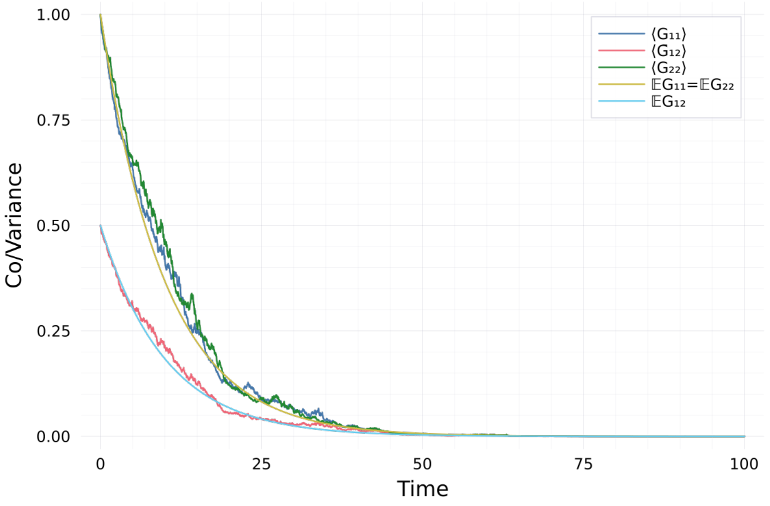


**Fig. 2.** Euler-Maruyama solutions to $\mathbf{G}$ with initial condition $G_{11} = G_{22} = 1.0, G_{12} = 0.5$ and rate $v/n = 0.1$. Each panel displays averages over 500 replicates ($\langle \mathbf{G} \rangle$) compared with the classical scaling result ($\mathbb{E}\mathbf{G}$).

#### 2. Calculation Steps for Expressions in Main Text

##### 2.1. Equation (1)

For the sake of convenience, the multivariate Deterministic Asexual Gaussian Allelic (DAGA) partial differential equation (equation (1) of the main text) is repeated here:

$$\dot{\nu}_t(\mathbf{z}) = m(\nu_t, \mathbf{z})\, \nu_t(\mathbf{z}) + \tfrac{1}{2} \nabla^\top \mathbf{M} \nabla\, \nu_t(\mathbf{z}). \tag{4}$$

This equation is obtained from a deterministic approximation to solutions of the martingale problem described in section 4 below. There are two ways to obtain this approximation: 1) assume the population size is very large or 2) assume the reproductive variance $v$ is very small. Further technical details associated with these approximation schemes are described in section 7.2 below.

---

#### 2.2. The Deterministic Covariance ($DC$) Version: Equations (3), (4), and (6)

##### *2.2.1. Equation (3)*

Given finite population abundance at time $t$ (i.e., $n_t < \infty$) and integrable time derivative of population density at $t$ (i.e., $\int_{\mathbb{R}^d} |\dot{\nu}_t(\mathbf{z})| d\mathbf{z} < \infty$), Lebesgue's Dominated Convergence theorem (see, Axler, 2020) can be applied to the limit definition of the time derivative to justify $\dot{n}_t = \int_{\mathbb{R}^d} \dot{\nu}_t(\mathbf{z}) d\mathbf{z}$. Substituting the right hand side of multivariate DAGA (equation (4) above) for $\dot{\nu}_t$ leads to

$$\dot{n}_t = \int_{\mathbb{R}^d} m(\nu_t, \mathbf{z})\,\nu_t(\mathbf{z}) + \tfrac{1}{2}\nabla^\top \mathbf{M} \nabla\, \nu_t(\mathbf{z}) d\mathbf{z}. \tag{5}$$

Because $\bar{m}_t = \int_{\mathbb{R}^d} m(\nu_t, \mathbf{z}) p_t(\mathbf{z}) d\mathbf{z} = \int_{\mathbb{R}^d} m(\nu_t, \mathbf{z}) \nu_t(\mathbf{z})/n_t d\mathbf{z}$, we have $\int_{\mathbb{R}^d} m(\nu_t, \mathbf{z}) \nu_t(\mathbf{z}) d\mathbf{z} = \bar{m}_t\, n_t$ (recall $p_t(\mathbf{z}) = \nu_t(\mathbf{z})/n_t$ is the trait distribution at time $t$, and $\bar{m}_t$ is mean fitness at time $t$).

By definition of matrix products, we immediately have $\int_{\mathbb{R}^d} \nabla^\top \mathbf{M} \nabla\, \nu_t(\mathbf{z}) d\mathbf{z} = \int_{\mathbb{R}^d} \sum_{ij} M_{ij} \partial_{z_i} \partial_{z_j} \nu_t d\mathbf{z} d\mathbf{z}$. Then, assuming $\nu_t(\mathbf{z}) \to 0$ and $|\nabla \nu_t(\mathbf{z})| \to 0$ as $|\mathbf{z}| \to \infty$ *sufficiently fast*, integration-by-parts with respect to $z_i$ shows $\int_{\mathbb{R}^d} \partial_{z_i} \partial_{z_j} \nu_t(\mathbf{z}) d\mathbf{z} d\mathbf{z} = -\int_{\mathbb{R}^d} \partial_{z_i} \nu_t(\mathbf{z}) \partial_{z_j}(1) d\mathbf{z} d\mathbf{z} = 0$. This holds for each $i = 1, \ldots, d$ which implies $\int_{\mathbb{R}^d} \nabla^\top \mathbf{M} \nabla\, \nu_t(\mathbf{z}) d\mathbf{z} = \int_{\mathbb{R}^d} \sum_{ij} M_{ij} \partial_{z_i} \partial_{z_j} \nu_t(\mathbf{z}) d\mathbf{z} d\mathbf{z} = \sum_{ij} M_{ij} \int_{\mathbb{R}^d} \partial_{z_i} \partial_{z_j} \nu_t(\mathbf{z}) d\mathbf{z} d\mathbf{z} = 0$.

Linearity of integration then implies $\dot{n}_t = \bar{m}_t\, n_t$, which is equation (3) of the main text.

. . . . . . . . . . . . . . . . . . . . . . . . . . . . . . . . . . . . . . . . . . . . . . . . . . . . . . . . . . . .

##### *2.2.2. Equation (4)*

At time $t$ the mean trait vector is defined by $\bar{\mathbf{z}}_t = \int_{\mathbb{R}^d} \mathbf{z}\, p_t(\mathbf{z}) d\mathbf{z}$. Assuming $\int_{\mathbb{R}^d} |\mathbf{z}| \left(p_t(\mathbf{z}) + |\dot{p}_t(\mathbf{z})|\right) d\mathbf{z} < \infty$ (with $|\mathbf{z}|$ being the Euclidean norm of $\mathbf{z}$), we can again apply Lebesgue's Dominated Convergence theorem to the limit definition of the time derivative to justify $\dot{\bar{\mathbf{z}}}_t = \int_{\mathbb{R}^d} \mathbf{z}\, \dot{p}_t(\mathbf{z}) d\mathbf{z}$.

The quotient rule applied to $\dot{p}_t(\mathbf{z}) = \partial_t[\nu_t(\mathbf{z})/n_t]$ provides

$$\dot{p}_t(\mathbf{z}) = \frac{n_t\, \dot{\nu}_t(\mathbf{z}) - \nu_t(\mathbf{z})\, \dot{n}_t}{n_t^2}. \tag{6}$$

Substituting the right-hand-side of DAGA (equation (4) above) for $\dot{\nu}_t(\mathbf{z})$, substituting the right-hand-side of the result $\dot{n}_t = \bar{m}_t\, n_t$ for $\dot{n}_t$, and then substituting $p_t(\mathbf{z}) = \nu_t(\mathbf{z})/n_t$ into equation (6) and simplifying, the dynamics of the trait distribution are then obtained as

$$\dot{p}_t(\mathbf{z}) = \left(m(n_t\, p_t, \mathbf{z}) - \bar{m}_t\right) p_t(\mathbf{z}) + \tfrac{1}{2}\nabla^\top \mathbf{M}\, \nabla\, p_t(\mathbf{z}). \tag{7}$$

with $m(n_t\, p_t, \mathbf{z}) = m(\nu_t, \mathbf{z})$ because $\nu_t(\mathbf{z}) = n_t\, p_t(\mathbf{z})$. Integration-by-parts is not needed to arrive at this result. The deterministic model summarized by DAGA (equation (4) above) is equally summarized by the combination of $\dot{n}_t = \bar{m}_t\, n_t$ and equation (7).

Now substituting the right-hand-side of (7) into $\dot{\bar{\mathbf{z}}}_t = \int_{\mathbb{R}^d} \mathbf{z}\, \dot{p}_t(\mathbf{z}) d\mathbf{z}$, we arrive at

$$\dot{\bar{\mathbf{z}}}_t = \int_{\mathbb{R}^d} \mathbf{z}\left(\left(m(n_t\, p_t, \mathbf{z}) - \bar{m}_t\right) + \tfrac{1}{2}\nabla^\top \mathbf{M}\, \nabla\right) p_t(\mathbf{z}) d\mathbf{z}. \tag{8}$$

The *selection term* involving $m(n_t\, p_t, \mathbf{z}) - \bar{m}_t$ can be rewritten as

$$\int_{\mathbb{R}^d} \mathbf{z}(m(n_t\, p_t, \mathbf{z}) - \bar{m}_t) p_t(\mathbf{z}) d\mathbf{z} = \int_{\mathbb{R}^d} (\mathbf{z} - \bar{\mathbf{z}}_t)(m(n_t\, p_t, \mathbf{z}) - \bar{m}_t) p_t(\mathbf{z}) d\mathbf{z} = \mathrm{Cov}_t(m, \mathbf{z}). \tag{9}$$

because

$$\int_{\mathbb{R}^d} \bar{\mathbf{z}}_t (m(n_t\, p_t, \mathbf{z}) - \bar{m}_t) p_t(\mathbf{z}) d\mathbf{z} = \mathbf{0}. \tag{10}$$

For the *mutation term* involving $\mathbf{M}$, note the $k$-th entry of the $d$-dimensional vector $\int_{\mathbb{R}^d} \mathbf{z} \nabla^\top \mathbf{M} \nabla \nu_t(\mathbf{z}) d\mathbf{z}$ can be written using index notation using the definition of matrix products as $\sum_{ij} M_{ij} \int_{\mathbb{R}^d} z_k\, \partial_{z_i}\, \partial_{z_j}\, p_t(\mathbf{z}) d\mathbf{z}$. Assuming $p_t(\mathbf{z}) \to 0$ and $|\nabla p_t(\mathbf{z})| \to 0$ *sufficiently fast* as $|\mathbf{z}| \to \infty$, integration-by-parts in $z_i$ shows $\int_{\mathbb{R}^d} z_k\, \partial_{z_i}\, \partial_{z_j}\, p_t(\mathbf{z}) d\mathbf{z} = -\int_{\mathbb{R}^d} \partial_{z_i}(z_k) \partial_{z_j} p_t(\mathbf{z}) d\mathbf{z} = -\delta_{ik} \int_{\mathbb{R}^d} \partial_{z_j} p_t(\mathbf{z}) d\mathbf{z}$. Thanks to the rapid decay of $p_t(\mathbf{z})$ for large $|\mathbf{z}|$, we have $\int_{\mathbb{R}^d} \partial_j p_t(\mathbf{z}) d\mathbf{z} = 0$. This holds for each $j, k = 1, \ldots, d$ and thus demonstrates $\dot{\bar{\mathbf{z}}}_t = \mathrm{Cov}_t(m, \mathbf{z})$ in agreement with equation (4) of the main text.

. . . . . . . . . . . . . . . . . . . . . . . . . . . . . . . . . . . . . . . . . . . . . . . . . . . . . . . . . . . .

##### 2.2.3. *Equation (6)*

At time $t$ the trait covariance matrix is defined by $\mathbf{P}_t = \int_{\mathbb{R}^d} (\mathbf{z}-\bar{\mathbf{z}})(\mathbf{z}-\bar{\mathbf{z}})^\top p_t(\mathbf{z})d\mathbf{z}$. Assuming

$$\int_{\mathbb{R}^d} (|\mathbf{z}| + |\mathbf{z}|^2)(p_t(\mathbf{z}) + |\dot{p}_t(\mathbf{z})|)d\mathbf{z} < \infty \tag{11}$$

Lebesgue's Dominated Convergence theorem can again be applied to justify $\dot{\mathbf{P}}_t = \int_{\mathbb{R}^d} \partial_t[(\mathbf{z}-\bar{\mathbf{z}})(\mathbf{z}-\bar{\mathbf{z}})^\top p_t(\mathbf{z})]d\mathbf{z}$. Expanding the partial time derivative inside the integral leads to

$$\int_{\mathbb{R}^d} \partial_t[(\mathbf{z}-\bar{\mathbf{z}})(\mathbf{z}-\bar{\mathbf{z}})^\top p_t(\mathbf{z})]d\mathbf{z} = \int_{\mathbb{R}^d} (\mathbf{z}-\bar{\mathbf{z}})(\mathbf{z}-\bar{\mathbf{z}})^\top \dot{p}_t(\mathbf{z})d\mathbf{z} + \int_{\mathbb{R}^d} p_t(\mathbf{z})\partial_t[(\mathbf{z}-\bar{\mathbf{z}})(\mathbf{z}-\bar{\mathbf{z}})^\top]d\mathbf{z}. \tag{12}$$

The product rule of elementary calculus shows that $\partial_t[(\mathbf{z}-\bar{\mathbf{z}}_t)(\mathbf{z}-\bar{\mathbf{z}}_t)^\top] = -\dot{\bar{\mathbf{z}}}_t(\mathbf{z}-\bar{\mathbf{z}}_t)^\top - (\mathbf{z}-\bar{\mathbf{z}}_t)\dot{\bar{\mathbf{z}}}_t^\top$. Hence, by the definition of $\bar{\mathbf{z}}_t$, we have $\int_{\mathbb{R}^d} p_t(\mathbf{z})\partial_t[(\mathbf{z}-\bar{\mathbf{z}}_t)(\mathbf{z}-\bar{\mathbf{z}}_t)^\top]d\mathbf{z} = 0$ and the dynamics of $\mathbf{P}_t$ simplify to

$$\dot{\mathbf{P}}_t = \int_{\mathbb{R}^d} (\mathbf{z}-\bar{\mathbf{z}}_t)(\mathbf{z}-\bar{\mathbf{z}}_t)^\top \dot{p}_t(\mathbf{z})d\mathbf{z}. \tag{13}$$

Replacing $\dot{p}_t(\mathbf{z})$ in equation (13) with the right-hand-side of equation (7), we arrive at

$$\dot{\mathbf{P}}_t = \int_{\mathbb{R}^d} (\mathbf{z}-\bar{\mathbf{z}}_t)(\mathbf{z}-\bar{\mathbf{z}}_t)^\top \left((m(n_t p_t, \mathbf{z}) - \bar{m}_t) + \tfrac{1}{2}\nabla^\top \mathbf{M} \nabla\right) p_t(\mathbf{z})d\mathbf{z}. \tag{14}$$

The *selection term* involving $m(n_t p_t, \mathbf{z}) - \bar{m}_t$ can be rewritten as

$$\int_{\mathbb{R}^d} (\mathbf{z}-\bar{\mathbf{z}}_t)(\mathbf{z}-\bar{\mathbf{z}}_t)^\top (m(n_t p_t, \mathbf{z}) - \bar{m}_t) p_t(\mathbf{z})d\mathbf{z} = \int_{\mathbb{R}^d} ((\mathbf{z}-\bar{\mathbf{z}}_t)(\mathbf{z}-\bar{\mathbf{z}}_t)^\top - \mathbf{P}_t)(m(n_t p_t, \mathbf{z}) - \bar{m}_t) p_t(\mathbf{z})d\mathbf{z} = \mathrm{Cov}_t(m, (\mathbf{z}-\bar{\mathbf{z}}_t)(\mathbf{z}-\bar{\mathbf{z}}_t)^\top) \tag{15}$$

because $\int_{\mathbb{R}^d} \mathbf{P}_t(m(n_t p_t, \mathbf{z}) - \bar{m}_t) p_t(\mathbf{z})d\mathbf{z} = \mathbf{0}$.

Set $\mathbf{U} = (\mathbf{z}-\bar{\mathbf{z}}_t)(\mathbf{z}-\bar{\mathbf{z}}_t)^\top$, the $d\times d$ matrix with $ij$-th entry $U_{ij} = (z_i - \bar{z}_i(t))(z_j - \bar{z}_j(t))$ and $\bar{z}_k(t)$ the $k$-th entry of $\bar{\mathbf{z}}_t$. Then, in index notation, the $kl$-th entry of the *mutation term* of equation (14) involving $\mathbf{M}$ is $\frac{1}{2}\int_{\mathbb{R}^d} U_{kl} \sum_{ij} M_{ij}\partial_{z_i}\partial_{z_j} p_t(\mathbf{z})d\mathbf{z}$. Given that $p_t(\mathbf{z}) \to 0$ and $|\nabla p_t(\mathbf{z})| \to 0$ *sufficiently fast* as $|\mathbf{z}| \to \infty$, apply integration-by-parts twice:

First in $z_i$: $\int_{\mathbb{R}^d} U_{kl} M_{ij}\partial_{z_i}\partial_{z_j} p_t(\mathbf{z})d\mathbf{z} = -\int_{\mathbb{R}^d} (\partial_{z_i} U_{kl}) M_{ij}\partial_{z_j} p_t(\mathbf{z})d\mathbf{z}$.

Then in $z_j$: $-\int_{\mathbb{R}^d} (\partial_{z_i} U_{kl}) M_{ij}\partial_{z_j} p_t(\mathbf{z})d\mathbf{z} = \int_{\mathbb{R}^d} (\partial_{z_i}\partial_{z_j} U_{kl}) M_{ij} p_t(\mathbf{z})d\mathbf{z}$.

So the $kl$-th entry of the *mutation term* of equation (14) becomes $\frac{1}{2}\int_{\mathbb{R}^d} \sum_{ij}(\partial_{z_i}\partial_{z_j} U_{kl}) M_{ij} p_t(\mathbf{z})d\mathbf{z}$. It is straightforward to obtain $\partial_{z_i}\partial_{z_j} U_{kl} = \delta_{ik}\delta_{jl} + \delta_{il}\delta_{jk}$, which does not depend on $\mathbf{z}$. So the $kl$-th entry of the *mutation term* simplifies to $\frac{1}{2}\sum_{ij}(\delta_{ik}\delta_{jl} + \delta_{il}\delta_{jk})M_{ij} = \frac{1}{2}(M_{kl} + M_{lk})$. Because $\mathbf{M}$ is a covariance matrix it also symmetric such that $M_{kl} = M_{lk}$. Hence, the *mutation term* of equation (14) simplifies to $\mathbf{M}$.

Taken together, we find $\dot{\mathbf{P}}_t = \mathbf{M} + \mathrm{Cov}_t(m, (\mathbf{z}-\bar{\mathbf{z}}_t)(\mathbf{z}-\bar{\mathbf{z}}_t)^\top)$ in agreement with equation (6) of the main text.

---

#### 2.3. Writing Covariances in Terms of Gradients

Here I show how to compute the Deterministic Gradient (DG) version of the framework from the Deterministic Covariance (DC) version. To begin, assume trait values are multivariate normal with non-singular covariance matrix so that

$$p(\mathbf{z}) = p(\mathbf{z}, \bar{\mathbf{z}}, \mathbf{P}) = \frac{\exp\left(-\frac{1}{2}(\mathbf{z}-\bar{\mathbf{z}})^\top \mathbf{P}^{-1}(\mathbf{z}-\bar{\mathbf{z}})\right)}{\sqrt{(2\pi)^k \det(\mathbf{P})}}. \tag{16}$$

Setting $\nabla_{\bar{\mathbf{z}}}$ the gradient with respect to $\bar{\mathbf{z}}$ (i.e., the $d$-dimensional vector with $i$-th entry $\partial/\partial\bar{z}_i$) and $q(\mathbf{z}) = \ln p(\mathbf{z})$, we have

$$\nabla_{\bar{\mathbf{z}}}\, q(\mathbf{z}) = -\tfrac{1}{2}\nabla_{\bar{\mathbf{z}}}[(\mathbf{z}-\bar{\mathbf{z}})^\top \mathbf{P}^{-1}(\mathbf{z}-\bar{\mathbf{z}})]. \tag{17}$$

Applying equation (83) from Petersen and Pedersen, (2012), we obtain $\nabla_{\bar{\mathbf{z}}}[(\mathbf{z}-\bar{\mathbf{z}})^\top \mathbf{P}^{-1}(\mathbf{z}-\bar{\mathbf{z}})] = 2\mathbf{P}^{-1}(\mathbf{z}-\bar{\mathbf{z}})$. Hence, $\nabla_{\bar{\mathbf{z}}}\, q(\mathbf{z}) = \mathbf{P}^{-1}(\mathbf{z}-\bar{\mathbf{z}})$. Combining this result with the standard log-derivative identity $\nabla_{\bar{\mathbf{z}}} \ln p(\mathbf{z}) = [\nabla_{\bar{\mathbf{z}}} p(\mathbf{z})]/p(\mathbf{z})$ brings us to

$$\nabla_{\bar{\mathbf{z}}}\, p(\mathbf{z}) = \mathbf{P}^{-1}(\mathbf{z}-\bar{\mathbf{z}})p(\mathbf{z}). \tag{18}$$

.........................................................

##### 2.3.1. *Covariance with trait expressed in terms of $\nabla_{\bar{\mathbf{z}}}$:*

For a function $x(\mathbf{z}, \bar{\mathbf{z}})$ that is differentiable with respect to $\bar{\mathbf{z}}$ and satisfies $\int_{\mathbb{R}^d} (|x(\mathbf{z}, \bar{\mathbf{z}})| + |\nabla_{\bar{\mathbf{z}}} x(\mathbf{z}, \bar{\mathbf{z}})|)p(\mathbf{z})d\mathbf{z} < \infty$, Lebesgue's Dominated Convergence theorem can be applied to the limit definition of $\nabla_{\bar{\mathbf{z}}}$ to justify

$$\nabla_{\bar{\mathbf{z}}}\, \bar{x} = \nabla_{\bar{\mathbf{z}}} \int_{\mathbb{R}^d} x(\mathbf{z}, \bar{\mathbf{z}})\, p(\mathbf{z})d\mathbf{z} = \int_{\mathbb{R}^d} p(\mathbf{z})\, [\nabla_{\bar{\mathbf{z}}} x(\mathbf{z}, \bar{\mathbf{z}})]d\mathbf{z} + \int_{\mathbb{R}^d} x(\mathbf{z}, \bar{\mathbf{z}})\, [\nabla_{\bar{\mathbf{z}}}\, p(\mathbf{z})]d\mathbf{z}. \tag{19}$$

Setting $\overline{\nabla_{\bar{\mathbf{z}}} x} = \int_{\mathbb{R}^d} p(\mathbf{z})\, [\nabla_{\bar{\mathbf{z}}} x]d\mathbf{z}$ and substituting $\nabla_{\bar{\mathbf{z}}} p(\mathbf{z})$ with the right-hand-side of equation (18), and simplifying using the definition of covariance results in

$$\nabla_{\bar{\mathbf{z}}}\, \bar{x} = \overline{\nabla_{\bar{\mathbf{z}}}\, x} + \mathbf{P}^{-1}\mathrm{Cov}(x, \mathbf{z}). \tag{20}$$

Rearranging yields $\mathrm{Cov}(x, \mathbf{z}) = \mathbf{P}\left(\nabla_{\bar{\mathbf{z}}}\, \bar{x} - \overline{\nabla_{\bar{\mathbf{z}}}\, x}\right)$.

. . . . . . . . . . . . . . . . . . . . . . . . . . . . . . . . . . . . . . . . . . . . . . . . . . . . . . . . . . . . . . . .

##### *2.3.2.* ***Covariance with*** $(\mathbf{z}-\bar{\mathbf{z}})(\mathbf{z}-\bar{\mathbf{z}})^\top$ ***expressed in terms of*** $\nabla_\mathbf{P}$***:***

Setting $\nabla_\mathbf{P}$ the gradient with respect to $\mathbf{P}$ (i.e., the $d\times d$ matrix with $ij$-th entry $\partial/\partial P_{ij}$), we can use formula presented in Petersen and Pedersen, (2012) to find $\nabla_\mathbf{P}\ln\det(\mathbf{P}) = \mathbf{P}^{-1}$ and $\nabla_\mathbf{P}\mathrm{tr}(\mathbf{A}\mathbf{P}^{-1}) = -(\mathbf{P}^{-1}\mathbf{A}\mathbf{P}^{-1})^\top$ for a $d\times d$ matrix $\mathbf{A}$. Applying these results to $\nabla_\mathbf{P}\ln p(\mathbf{z})$ and simplifying yields

$$\nabla_\mathbf{P}\,\ln p(\mathbf{z}) = \tfrac{1}{2}\mathbf{P}^{-1}[(\mathbf{z}-\bar{\mathbf{z}})(\mathbf{z}-\bar{\mathbf{z}})^\top - \mathbf{P}]\mathbf{P}^{-1}. \tag{21}$$

Finally, use the relation $\nabla_\mathbf{P}p(\mathbf{z}) = p(\mathbf{z})\nabla_\mathbf{P}\ln p(\mathbf{z})$ to obtain $\nabla_\mathbf{P}p(\mathbf{z})$. None of these calculations assume constancy of $\det(\mathbf{P})$ with respect to $\mathbf{P}$ or any other special assumptions.

For a function $x(\mathbf{z},\mathbf{P})$ that is differentiable with respect to $\mathbf{P}$ and satisfies $\int_{\mathbb{R}^d}(|x(\mathbf{z},\mathbf{P})| + |\nabla_\mathbf{P}x(\mathbf{z},\mathbf{P})|)p(\mathbf{z})d\mathbf{z} < \infty$, we can combine Lebesgue's Dominated Convergence theorem with the limit definition of $\nabla_\mathbf{P}$ to justify $\nabla_\mathbf{P}\int_{\mathbb{R}^d}x(\mathbf{z},\mathbf{P})\,p(\mathbf{z})d\mathbf{z} = \int_{\mathbb{R}^d}\nabla_\mathbf{P}[x(\mathbf{z},\mathbf{P})\,p(\mathbf{z})]d\mathbf{z}$. Using barred expressions for brevity, we then have $\nabla_\mathbf{P}\,\bar{x} = \overline{\nabla_\mathbf{P}\,x} + \int_{\mathbb{R}^d}x(\mathbf{z})[\nabla_\mathbf{P}\,p(\mathbf{z})]\,d\mathbf{z}$. Equation (21) implies $\int_{\mathbb{R}^d}x(\mathbf{z})[\nabla_\mathbf{P}\,p(\mathbf{z})]\,d\mathbf{z} = \frac{1}{2}\mathbf{P}^{-1}\mathrm{Cov}(x,(\mathbf{z}-\bar{\mathbf{z}})(\mathbf{z}-\bar{\mathbf{z}})^\top)\mathbf{P}^{-1}$. We then obtain

$$\mathrm{Cov}(x,(\mathbf{z}-\bar{\mathbf{z}})(\mathbf{z}-\bar{\mathbf{z}})^\top) = 2\,\mathbf{P}\,(\nabla_\mathbf{P}\,\bar{x} - \overline{\nabla_\mathbf{P}\,x})\,\mathbf{P}. \tag{22}$$

. . . . . . . . . . . . . . . . . . . . . . . . . . . . . . . . . . . . . . . . . . . . . . . . . . . . . . . . . . . . . . . .

##### *2.3.3.* ***Covariance with mean trait expressed in terms of*** $\nabla$***:***

To rewrite $\mathrm{Cov}(m,\mathbf{z})$ in terms of $\nabla := (\partial/\partial z_1,\ldots,\partial/\partial z_d)^\top$ (i.e., the gradient with respect to $\mathbf{z}$), note that for multivariate normal $p(\mathbf{z})$ with mean $\bar{\mathbf{z}}$ and covariance $\mathbf{P}$ we have $\nabla_{\bar{\mathbf{z}}}\ln p(\mathbf{z}) = \mathbf{P}^{-1}(\mathbf{z}-\bar{\mathbf{z}})$, and $\nabla\ln p(\mathbf{z}) = -\mathbf{P}^{-1}(\mathbf{z}-\bar{\mathbf{z}})$, and hence

$$\nabla_{\bar{\mathbf{z}}}p(\mathbf{z}) = p(\mathbf{z})\nabla_{\bar{\mathbf{z}}}\ln p(\mathbf{z}) = -p(\mathbf{z})\nabla\ln p(\mathbf{z}) = -\nabla p(\mathbf{z}). \tag{23}$$

In section 2.2.1 above we already obtained

$$\nabla_{\bar{\mathbf{z}}}\bar{x} = \nabla_{\bar{\mathbf{z}}}\int x(\mathbf{z},\bar{\mathbf{z}})\,p(\mathbf{z})\,d\mathbf{z} = \overline{\nabla_{\bar{\mathbf{z}}}x} + \int x(\mathbf{z},\bar{\mathbf{z}})[\nabla_{\bar{\mathbf{z}}}p(\mathbf{z})]\,d\mathbf{z}. \tag{24}$$

Thus,

$$\nabla_{\bar{\mathbf{z}}}\bar{x} - \overline{\nabla_{\bar{\mathbf{z}}}x} = -\int x(\mathbf{z},\bar{\mathbf{z}})[\nabla p(\mathbf{z})]\,d\mathbf{z}. \tag{25}$$

Because of multivariate normality, $p(\mathbf{z})$ and $|\nabla p(\mathbf{z})|$ decay sufficiently rapidly as $|\mathbf{z}|\to\infty$ to justify integration-by-parts. For coordinate $i$, this gives $-\int_{\mathbb{R}^d}x(\mathbf{z},\bar{\mathbf{z}})\,\partial_{z_i}p(\mathbf{z})d\mathbf{z} = \int_{\mathbb{R}^d}(\partial_{z_i}x(\mathbf{z},\bar{\mathbf{z}}))\,p(\mathbf{z})d\mathbf{z}$. In vector form this is $\nabla_{\bar{\mathbf{z}}}\bar{x} - \overline{\nabla_{\bar{\mathbf{z}}}x} = \overline{\nabla x}$. Hence, we can equally write $\mathrm{Cov}(m,\mathbf{z}) = \mathbf{P}\,\overline{\nabla m}$.

. . . . . . . . . . . . . . . . . . . . . . . . . . . . . . . . . . . . . . . . . . . . . . . . . . . . . . . . . . . . . . . .

##### *2.3.4.* ***Covariance with*** $(\mathbf{z}-\bar{\mathbf{z}})(\mathbf{z}-\bar{\mathbf{z}})^\top$ ***expressed in terms of*** $\nabla^2$***:***

To rewrite the response of $\mathbf{P}$ to selection in terms of the Hessian with respect to $\mathbf{z}$,

$$\nabla^2 := \begin{pmatrix} \frac{\partial^2}{\partial z_1^2} & \frac{\partial^2}{\partial z_1\partial z_2} & \cdots & \frac{\partial^2}{\partial z_1\partial z_d} \\ \frac{\partial^2}{\partial z_2\partial z_1} & \frac{\partial^2}{\partial z_2^2} & \cdots & \frac{\partial^2}{\partial z_2\partial z_d} \\ \vdots & \vdots & \ddots & \vdots \\ \frac{\partial^2}{\partial z_d\partial z_1} & \frac{\partial^2}{\partial z_d\partial z_2} & \cdots & \frac{\partial^2}{\partial z_d^2} \end{pmatrix}, \tag{26}$$

first note that, by using multivariate normality, we can leverage results in section 8.4.2 of Petersen and Pedersen, (2012) to obtain $\nabla_\mathbf{P}p(\mathbf{z}) = \frac{1}{2}\nabla^2 p(\mathbf{z})$. In addition, because of multivariate normality $p(\mathbf{z})$ and $|\nabla p(\mathbf{z})|$ decay sufficiently rapidly as $|\mathbf{z}|\to\infty$ to justify applying integration-by-parts twice to $\left(\int_{\mathbb{R}^d}x(\mathbf{z})[\nabla_\mathbf{P}\,p(\mathbf{z})]\,d\mathbf{z}\right)_{ij} = \int_{\mathbb{R}^d}x(\mathbf{z})\,\partial_{z_i}\partial_{z_j}\,p(\mathbf{z})d\mathbf{z}$:

- First in $z_i$: $\int_{\mathbb{R}^d}x(\mathbf{z})\,\partial_{z_i}\partial_{z_j}\,p(\mathbf{z})d\mathbf{z} = -\int_{\mathbb{R}^d}[\partial_{z_i}x(\mathbf{z})]\partial_{z_j}\,p(\mathbf{z})d\mathbf{z}$.
- Then in $z_j$: $-\int_{\mathbb{R}^d}[\partial_{z_i}x(\mathbf{z})]\partial_{z_j}\,p(\mathbf{z})d\mathbf{z} = \int_{\mathbb{R}^d}[\partial_{z_i}\partial_{z_j}x(\mathbf{z})]\,p(\mathbf{z})d\mathbf{z}$.

In the notation above this gives $\nabla_\mathbf{P}\bar{x} - \overline{\nabla_\mathbf{P}x} = \frac{1}{2}\overline{\nabla^2 x}$. Hence, we can equally write $\mathrm{Cov}(x,(\mathbf{z}-\bar{\mathbf{z}})(\mathbf{z}-\bar{\mathbf{z}})^\top) = \mathbf{P}\,\overline{\nabla^2 x}\,\mathbf{P}$.

#### 2.4. Equations (8)

Here I show how to combine the general dynamics

$$\dot{n} = \bar{m}\,n, \tag{27a}$$

$$\dot{\bar{\mathbf{z}}} = \mathrm{Cov}(m, \mathbf{z}), \tag{27b}$$

$$\dot{\mathbf{P}} = \mathbf{M} + \mathrm{Cov}(m, (\mathbf{z}-\bar{\mathbf{z}})(\mathbf{z}-\bar{\mathbf{z}})^\top), \tag{27c}$$

with a fitness function of the form $m(\nu, \mathbf{z}) = r + \mathbf{b}^\top\mathbf{z} - \frac{1}{2}(\boldsymbol{\theta}-\mathbf{z})^\top\boldsymbol{\Psi}\,(\boldsymbol{\theta}-\mathbf{z}) - cn$ to arrive at equations (8) in the main text. These derivations assume the abundance, mean vector, covariance matrix, and skew and kurtosis tensors are finite, but no other restrictions are required.

. . . . . . . . . . . . . . . . . . . . . . . . . . . . . . . . . . . . . . . . . . . . . . . . . . . . . . . . . . . .

##### 2.4.1. *Equation (8a)*

To begin, mean fitness $\bar{m}$ can be obtained by first expanding the quadratic term quantifying stabilizing selection in $m$ above and rearranging to get

$$m(\nu, \mathbf{z}) = (r - \tfrac{1}{2}\boldsymbol{\theta}^\top\boldsymbol{\Psi}\,\boldsymbol{\theta} - cn) + (\mathbf{b}^\top + \boldsymbol{\theta}^\top\boldsymbol{\Psi})\mathbf{z} - \tfrac{1}{2}\mathbf{z}^\top\boldsymbol{\Psi}\mathbf{z} \tag{28}$$

Note that the trace of a scalar is just that scalar so that $\mathbf{z}^\top\boldsymbol{\Psi}\mathbf{z} = \mathrm{tr}(\mathbf{z}^\top\boldsymbol{\Psi}\mathbf{z})$. The trace operator has a so-called cyclic property such that $\mathrm{tr}(\mathbf{ABC}) = \mathrm{tr}(\mathbf{BCA}) = \mathrm{tr}(\mathbf{CAB})$ (see Petersen and Pedersen, 2012), so we can write $\mathbf{z}^\top\boldsymbol{\Psi}\mathbf{z} = \mathrm{tr}(\boldsymbol{\Psi}\mathbf{z}\mathbf{z}^\top)$. Then because both the operations of taking the trace and averaging across traits are linear, we have $\overline{\mathbf{z}^\top\boldsymbol{\Psi}\mathbf{z}} = \mathrm{tr}(\boldsymbol{\Psi}\overline{\mathbf{z}\mathbf{z}^\top})$. But then we also have $\overline{\mathbf{z}\mathbf{z}^\top} = \mathbf{P} + \bar{\mathbf{z}}\bar{\mathbf{z}}^\top$ so that (again using the cyclic property of trace) $\overline{\mathbf{z}^\top\boldsymbol{\Psi}\mathbf{z}} = \mathrm{tr}(\boldsymbol{\Psi}\mathbf{P}) + \bar{\mathbf{z}}^\top\boldsymbol{\Psi}\bar{\mathbf{z}}$. Hence, mean fitness is obtained as

$$\bar{m} = (r - \tfrac{1}{2}\boldsymbol{\theta}^\top\boldsymbol{\Psi}\,\boldsymbol{\theta} - cn) + (\mathbf{b}^\top + \boldsymbol{\theta}^\top\boldsymbol{\Psi})\bar{\mathbf{z}} - \tfrac{1}{2}(\mathrm{tr}(\boldsymbol{\Psi}\mathbf{P}) + \bar{\mathbf{z}}^\top\boldsymbol{\Psi}\bar{\mathbf{z}}, \tag{29}$$

and rearranging brings us to

$$\bar{m} = r + \mathbf{b}^\top\bar{\mathbf{z}} - \tfrac{1}{2}(\boldsymbol{\theta}-\bar{\mathbf{z}})^\top\boldsymbol{\Psi}\,(\boldsymbol{\theta}-\bar{\mathbf{z}}) - \tfrac{1}{2}\mathrm{tr}(\boldsymbol{\Psi}\,\mathbf{P}) - cn. \tag{30}$$

Plugging this into $\dot{n} = \bar{m}\,n$ provides equation (8a) in the main text.

. . . . . . . . . . . . . . . . . . . . . . . . . . . . . . . . . . . . . . . . . . . . . . . . . . . . . . . . . . . .

##### 2.4.2. *Equation (8b)*

For the mean trait dynamics, set $\mathbf{x} = \mathbf{z} - \bar{\mathbf{z}}$ so that $\mathrm{Cov}(m, \mathbf{z}) = \overline{m\mathbf{x}}$. Rewriting $m$ in terms of $\mathbf{x}$ provides (making use of symmetry of $\boldsymbol{\Psi}$)

$$m(\nu, \mathbf{x}) = r + \mathbf{b}^\top\bar{\mathbf{z}} - \tfrac{1}{2}(\boldsymbol{\theta}-\bar{\mathbf{z}})^\top\boldsymbol{\Psi}\,(\boldsymbol{\theta}-\bar{\mathbf{z}}) - cn + (\mathbf{b} + \boldsymbol{\Psi}(\boldsymbol{\theta}-\bar{\mathbf{z}}))^\top\mathbf{x} - \tfrac{1}{2}\mathbf{x}^\top\boldsymbol{\Psi}\mathbf{x}. \tag{31}$$

Setting $\mathbf{a} = \mathbf{b} + \boldsymbol{\Psi}(\boldsymbol{\theta}-\bar{\mathbf{z}})$, we then have $\mathrm{Cov}(m, \mathbf{z}) = \overline{(\mathbf{a}^\top\mathbf{x})\mathbf{x}} - \frac{1}{2}\overline{(\mathbf{x}^\top\boldsymbol{\Psi}\mathbf{x})\mathbf{x}}$ because $\bar{\mathbf{x}} = 0$. Using the matrix identity $(\mathbf{a}^\top\mathbf{x})\mathbf{x} = \mathbf{x}\mathbf{x}^\top\mathbf{a}$ and linearity of averaging across trait values, we have $\overline{(\mathbf{a}^\top\mathbf{x})\mathbf{x}} = \mathbf{P}\mathbf{a}$ because $\overline{\mathbf{x}\mathbf{x}^\top} = \mathbf{P}$.

Now observe that component $i$ of $\overline{(\mathbf{x}^\top\boldsymbol{\Psi}\mathbf{x})\mathbf{x}}$ is $\sum_{jk}\Psi_{jk}\overline{x_i x_j x_k}$, and $S_{ijk} = \overline{x_i x_j x_k}$ is the $ijk$-th entry of the third order skew tensor $\mathbf{S}$. In particular, this coincides with the $i$-th entry of the product $\mathbf{S} : \boldsymbol{\Psi}$ defined in the main text. Together this provides $\mathrm{Cov}(m, \mathbf{z}) = \mathbf{P}\mathbf{b} + \mathbf{P}\boldsymbol{\Psi}(\boldsymbol{\theta}-\bar{\mathbf{z}}) - \frac{1}{2}\mathbf{S} : \boldsymbol{\Psi}$, which agrees with equation (8b) of the main text.

. . . . . . . . . . . . . . . . . . . . . . . . . . . . . . . . . . . . . . . . . . . . . . . . . . . . . . . . . . . .

##### 2.4.3. *Equation (8c)*

Starting with $\dot{\mathbf{P}} = \mathbf{M} + \mathrm{Cov}(m, (\mathbf{z}-\bar{\mathbf{z}})(\mathbf{z}-\bar{\mathbf{z}})^\top)$, set $\mathbf{x} = \mathbf{z} - \bar{\mathbf{z}}$, so that $\mathbf{P} = \overline{\mathbf{x}\mathbf{x}^\top}$ and $\bar{\mathbf{x}} = 0$. Then

$$\dot{\mathbf{P}} = \mathbf{M} + \mathrm{Cov}(m, \mathbf{x}\mathbf{x}^\top)$$

and

$$m = r + \mathbf{b}^\top\bar{\mathbf{z}} - \tfrac{1}{2}(\boldsymbol{\theta}-\bar{\mathbf{z}})^\top\boldsymbol{\Psi}(\boldsymbol{\theta}-\bar{\mathbf{z}}) - cn + \mathbf{a}^\top\mathbf{x} - \tfrac{1}{2}\mathbf{x}^\top\boldsymbol{\Psi}\mathbf{x},$$

where $\mathbf{a} = \mathbf{b} + \boldsymbol{\Psi}(\boldsymbol{\theta}-\bar{\mathbf{z}})$. Then,

$$\mathrm{Cov}(m, \mathbf{x}\mathbf{x}^\top) = \mathrm{Cov}(\mathbf{a}^\top\mathbf{x}, \mathbf{x}\mathbf{x}^\top) - \tfrac{1}{2}\mathrm{Cov}(\mathbf{x}^\top\boldsymbol{\Psi}\mathbf{x}, \mathbf{x}\mathbf{x}^\top),$$

and

$$\mathrm{Cov}(\mathbf{a}^\top\mathbf{x}, \mathbf{x}\mathbf{x}^\top) = \overline{(\mathbf{a}^\top\mathbf{x})\mathbf{x}\mathbf{x}^\top}.$$

The $ijk$-th entry of the third-order skew tensor $\mathbf{S}$ is given by $S_{ijk} = \overline{x_i x_j x_k}$. Denoting the single contraction by $(\mathbf{S}\cdot\mathbf{a})_{ij} = \sum_k S_{ijk}a_k$, we then have $\mathrm{Cov}(\mathbf{a}^\top\mathbf{x}, \mathbf{x}\mathbf{x}^\top) = \mathbf{S}\cdot\mathbf{a}$. The $ijkl$-th entry of the fourth-order kurtosis tensor $\mathbf{K}$ is given by $K_{ijkl} = \overline{x_i x_j x_k x_l}$. Denote the double contraction by $(\mathbf{K} : \boldsymbol{\Psi})_{ij} = \sum_{kl} K_{ijkl}\Psi_{kl}$. Then $\overline{\mathbf{x}^\top\boldsymbol{\Psi}\mathbf{x}} = \mathrm{tr}(\boldsymbol{\Psi}\mathbf{P})$, and using the identity $\mathrm{Cov}(x, y) = \overline{xy} - \bar{x}\bar{y}$, we obtain

$$\mathrm{Cov}(\mathbf{x}^\top\mathbf{\Psi}\mathbf{x}, \mathbf{x}\mathbf{x}^\top) = (\mathbf{K} : \mathbf{\Psi}) - \mathrm{tr}(\mathbf{\Psi}\mathbf{P})\,\mathbf{P}.$$

Because $\mathbf{S}\cdot\mathbf{a} = \mathbf{S}\cdot(\mathbf{b} + \mathbf{\Psi}(\boldsymbol{\theta} - \bar{\mathbf{z}})) = \mathbf{S}\cdot\mathbf{b} + \mathbf{S}\cdot(\mathbf{\Psi}(\boldsymbol{\theta} - \bar{\mathbf{z}}))$, we arrive at

$$\mathrm{Cov}(m, \mathbf{x}\mathbf{x}^\top) = \mathbf{S}\cdot\mathbf{b} + \mathbf{S}\cdot(\mathbf{\Psi}(\boldsymbol{\theta} - \bar{\mathbf{z}})) - \tfrac{1}{2}(\mathbf{K} : \mathbf{\Psi}) + \tfrac{1}{2}\,\mathrm{tr}(\mathbf{\Psi}\mathbf{P})\mathbf{P},$$

which then justifies equation (8c) of the main text.

---

#### 2.5. Equilibrium of $n, \bar{\mathbf{z}}, \mathbf{P}$ when $m$ takes the form of equation (7) in the main text

##### *2.5.1. Existence of Guassian Equilibrium and justification for equations (9)-(11) in main text*

From equation (7) in supplementary material section 2.1.3 above, we have $\dot{p} = (m - \bar{m})p + \frac{1}{2}\nabla^\top\mathbf{M}\nabla p$. Consider a Gaussian density of the form

$$f(\mathbf{z}) = Z^{-1}\exp\Big(-\tfrac{1}{2}(\mathbf{z} - \boldsymbol{\mu})^\top\mathbf{Q}^{-1}(\mathbf{z} - \boldsymbol{\mu})\Big), \tag{32}$$

with mean $\boldsymbol{\mu}$, covariance matrix $\mathbf{Q}$, and normalizing constant $Z$. Writing $m(\mathbf{z})$ in place of $m(\nu, \mathbf{z})$, follow equation (7) in the main text and assume

$$m(\mathbf{z}) = r + \mathbf{b}^\top\mathbf{z} - \tfrac{1}{2}(\boldsymbol{\theta} - \mathbf{z})^\top\mathbf{\Psi}(\boldsymbol{\theta} - \mathbf{z}) - cn. \tag{33}$$

From (30) above, we already have

$$\bar{m} = \int_{\mathbb{R}^d} m(\mathbf{z})f(\mathbf{z})d\mathbf{z} = r + \mathbf{b}^\top\boldsymbol{\mu} - \tfrac{1}{2}(\boldsymbol{\theta} - \boldsymbol{\mu})^\top\mathbf{\Psi}\,(\boldsymbol{\theta} - \boldsymbol{\mu}) - \tfrac{1}{2}\mathrm{tr}(\mathbf{\Psi}\,\mathbf{Q}) - cn, \tag{34}$$

which holds for any distribution of $\mathbf{z}$ (with finite first and second moments). Set $\mathbf{y} := \mathbf{z} - \boldsymbol{\mu}$. Then $(\boldsymbol{\theta} - \mathbf{z}) = (\boldsymbol{\theta} - \boldsymbol{\mu}) - \mathbf{y}$ and

$$(\boldsymbol{\theta} - \mathbf{z})^\top\mathbf{\Psi}(\boldsymbol{\theta} - \mathbf{z}) = (\boldsymbol{\theta} - \boldsymbol{\mu})^\top\mathbf{\Psi}(\boldsymbol{\theta} - \boldsymbol{\mu}) - 2\mathbf{y}^\top\mathbf{\Psi}(\boldsymbol{\theta} - \boldsymbol{\mu}) + \mathbf{y}^\top\mathbf{\Psi}\mathbf{y}. \tag{35}$$

Plugging this into $m(\mathbf{z})$, and then using this to simplify $m(\mathbf{z}) - \bar{m}$ leads to

$$m(\mathbf{z}) - \bar{m} = \mathbf{a}^\top(\mathbf{z} - \boldsymbol{\mu}) - \tfrac{1}{2}(\mathbf{z} - \boldsymbol{\mu})^\top\mathbf{\Psi}(\mathbf{z} - \boldsymbol{\mu}) + \tfrac{1}{2}\mathrm{tr}(\mathbf{\Psi}\mathbf{Q}), \tag{36}$$

where $\mathbf{a} = \mathbf{b} + \mathbf{\Psi}\boldsymbol{\theta}$.

From the form of $f$ assumed, we have $\ln f(\mathbf{z}) = -\ln Z - \frac{1}{2}(\mathbf{z} - \boldsymbol{\mu})^\top\mathbf{Q}^{-1}(\mathbf{z} - \boldsymbol{\mu})$. Using Petersen and Pedersen, (2012), equation (83), the gradient of a quadratic form is $\nabla\big[(\mathbf{z} - \boldsymbol{\mu})^\top\mathbf{Q}^{-1}(\mathbf{z} - \boldsymbol{\mu})\big] = 2\,\mathbf{Q}^{-1}(\mathbf{z} - \boldsymbol{\mu})$. Therefore, $\nabla\ln f(\mathbf{z}) = -\mathbf{Q}^{-1}(\mathbf{z} - \boldsymbol{\mu})$, and by the log-derivative identity, $\nabla f(\mathbf{z}) = -\mathbf{Q}^{-1}(\mathbf{z} - \boldsymbol{\mu})\,f(\mathbf{z})$. Differentiating again gives $\nabla\nabla^\top\ln f(\mathbf{z}) = -\mathbf{Q}^{-1}$. Using the identity for derivatives of exponentials,

$$\nabla\nabla^\top f(\mathbf{z}) = f(\mathbf{z})\,\nabla\nabla^\top\ln f(\mathbf{z}) + f(\mathbf{z})\,[\nabla\ln f(\mathbf{z})][\nabla\ln f(\mathbf{z})]^\top, \tag{37}$$

and hence

$$\nabla\nabla^\top f(\mathbf{z}) = f(\mathbf{z})\Big[-\mathbf{Q}^{-1} + \mathbf{Q}^{-1}(\mathbf{z} - \boldsymbol{\mu})(\mathbf{z} - \boldsymbol{\mu})^\top\mathbf{Q}^{-1}\Big]. \tag{38}$$

Using $\nabla^\top\mathbf{M}\nabla f = \mathrm{tr}(\mathbf{M}\,\nabla\nabla^\top f)$, we obtain

$$\nabla^\top\mathbf{M}\nabla f(\mathbf{z}) = f(\mathbf{z})\Big[(\mathbf{z} - \boldsymbol{\mu})^\top\mathbf{Q}^{-1}\mathbf{M}\mathbf{Q}^{-1}(\mathbf{z} - \boldsymbol{\mu}) - \mathrm{tr}(\mathbf{M}\mathbf{Q}^{-1})\Big]. \tag{39}$$

Divide the stationary condition $0 = (m(\mathbf{z}) - \bar{m})f(\mathbf{z}) + \frac{1}{2}\nabla^\top\mathbf{M}\nabla f(\mathbf{z})$ by $f(\mathbf{z})$ to get

$$0 = m(\mathbf{z}) - \bar{m} + \tfrac{1}{2}\Big[(\mathbf{z} - \boldsymbol{\mu})^\top\mathbf{Q}^{-1}\mathbf{M}\mathbf{Q}^{-1}(\mathbf{z} - \boldsymbol{\mu}) - \mathrm{tr}(\mathbf{M}\mathbf{Q}^{-1})\Big]. \tag{40}$$

Plugging the result for $m(\mathbf{z}) - \bar{m}$ returns

$$-\mathbf{a}^\top(\mathbf{z} - \boldsymbol{\mu}) + \tfrac{1}{2}(\mathbf{z} - \boldsymbol{\mu})^\top\mathbf{\Psi}(\mathbf{z} - \boldsymbol{\mu}) - \tfrac{1}{2}\mathrm{tr}(\mathbf{\Psi}\mathbf{Q}) = \tfrac{1}{2}(\mathbf{z} - \boldsymbol{\mu})^\top\mathbf{Q}^{-1}\mathbf{M}\mathbf{Q}^{-1}(\mathbf{z} - \boldsymbol{\mu}) - \tfrac{1}{2}\mathrm{tr}(\mathbf{M}\mathbf{Q}^{-1}). \tag{41}$$

Now match coefficients. For the quadratic term, we have $\mathbf{\Psi} = \mathbf{Q}^{-1}\mathbf{M}\mathbf{Q}^{-1}$. Equivalently, $\mathbf{M} = \mathbf{Q}\,\mathbf{\Psi}\,\mathbf{Q}$. Denoting $\mathbf{\Psi}^{1/2}$ the principle square root of $\mathbf{\Psi}$ (defined using the eigenvalue decomposition $b\Psi^{1/2} = \mathbf{U}\mathbf{\Lambda}^{1/2}\mathbf{U}$), and multiplying both sides of $\mathbf{M}$ by $\mathbf{\Psi}^{1/2}$ provides

$$\mathbf{\Psi}^{1/2}\mathbf{M}\mathbf{\Psi}^{1/2} = \mathbf{\Psi}^{1/2}\mathbf{Q}\mathbf{\Psi}\mathbf{Q}\mathbf{\Psi}^{1/2} = \mathbf{\Psi}^{1/2}\mathbf{Q}\mathbf{\Psi}^{1/2}\mathbf{\Psi}^{1/2}\mathbf{Q}\mathbf{\Psi}^{1/2} = (\mathbf{\Psi}^{1/2}\mathbf{Q}\mathbf{\Psi}^{1/2})^2. \tag{42}$$

Hence $\mathbf{Q} = \mathbf{\Psi}^{1/2}(\mathbf{\Psi}^{1/2}\mathbf{M}\mathbf{\Psi}^{1/2})^{1/2}\mathbf{\Psi}^{1/2}$, in agreement with the expression for $\hat{\mathbf{P}}$ given by equation (9) in the main text. For the linear term, we have $\mathbf{a} = 0$, which shows $\boldsymbol{\mu} = \mathbf{\Psi}^{-1}\mathbf{b} + \boldsymbol{\theta}$ in agreement with the expression for $\hat{\bar{\mathbf{z}}}$ given by equation (10) in the main text. For the constant trace terms, we already have $\mathbf{Q}\mathbf{\Psi} = \mathbf{M}\mathbf{Q}^{-1}$. So the cyclic property of traces shows $\mathrm{tr}(\mathbf{\Psi}\mathbf{Q}) = \mathrm{tr}(\mathbf{M}\mathbf{Q}^{-1})$. Plugging in the equilibrium for mean trait vector and trait covariance matrix into the non-trivial equilibrium condition for abundance ($\dot{n} = 0$

implies $\bar{m} = 0$ if $\hat{n} > 0$) returns equation (11) in the main text. Then, since the Gaussian ansatz is consistent, the stationary density is exactly

$$f(\mathbf{z}) = \hat{p}(\mathbf{z}) = \frac{1}{\sqrt{(2\pi)^d \det(\hat{\mathbf{P}})}} \exp\Big(-\tfrac{1}{2}(\mathbf{z}-\hat{\bar{\mathbf{z}}})^\top \hat{\mathbf{P}}^{-1}(\mathbf{z}-\hat{\bar{\mathbf{z}}})\Big). \tag{43}$$

This demonstrates that, for sufficiently large intrinsic growth $r > 0$, multivariate DAGA (equation (1) in the main text) has an equilibrium proportional to a Gaussian density $\hat{p}(\mathbf{z})$ with covariance matrix $\hat{\mathbf{P}}$, mean $\hat{\bar{\mathbf{z}}}$, and constant of proportionality $\hat{n}$ given by equations (9)-(11) of the main text. Sufficiently large intrinsic growth $r$ means that $\bar{m}_0 := \int_{\mathbb{R}^d} m_0(\mathbf{z})\hat{p}(\mathbf{z})\,d\mathbf{z} > 0$ with $m_0(\mathbf{z}) = m(\nu,\mathbf{z})|_{n=0}$. In that case $\dot{n} = 0$ implies $\bar{m} = 0$ and hence $\hat{n} = \bar{m}_0/c$, while if $\bar{m}_0 \le 0$ the equilibrium abundance is $\hat{n} = 0$. I next show that the non-trivial equilibrium is globally stable given the initial condition satisfies $n_0 = \int_{\mathbb{R}^d} \nu_0(\mathbf{z})d\mathbf{z} > 0$.

. . . . . . . . . . . . . . . . . . . . . . . . . . . . . . . . . . . . . . . . . . . . . . . . . . . . . . . . . . . .

##### *2.5.2. Global convergence of population density*

Multivariate DAGA is given by $\dot{\nu} = m\nu + \frac{1}{2}\nabla^\top\mathbf{M}\nabla\nu$. From equation (7) in Supplementary Material, Section 2.1.3 above, we then have $\dot{p} = (m-\bar{m})p + \frac{1}{2}\nabla^\top\mathbf{M}\nabla p$. The specific form of $m$ assumed is $m(\nu,\mathbf{z}) = m(\mathbf{z}) = r + \mathbf{b}^\top\mathbf{z} - \frac{1}{2}(\mathbf{z}-\boldsymbol{\theta})^\top\boldsymbol{\Psi}(\mathbf{z}-\boldsymbol{\theta}) - cn$. Hence $m-\bar{m}$ is independent of $n$, and therefore so are the dynamics of $p$. Equilibrium $\hat{p}$ given above satisfies $0 = (m-\hat{\bar{m}})\hat{p} + \frac{1}{2}\nabla^\top\mathbf{M}\nabla\hat{p}$, and equilibrium $\hat{n}$ satisfies $0 = \hat{\bar{m}}\hat{n}$. Given sufficiently large intrinsic growth rate $r$, we have $\hat{n} > 0$ and thus $\hat{\bar{m}} = 0$. Together this implies equilibrium population density is $\hat{\nu} = \hat{n}\hat{p}$.

To demonstrate $\nu_t \to \hat{\nu}$ as $t \to \infty$ whenever $n_0 = \int_{\mathbb{R}^d}\nu_0(\mathbf{z})d\mathbf{z} > 0$ and $r$ is large enough, I take advantage of the independence of $\dot{p}_t$ from $n_t$ and consider a standard expression for the relative entropy of $p_t$ to $\hat{p}$. I show this quantity is decreasing, and use this result to show $p_t \to \hat{p}$. Building on this, I then show $\bar{m}_t \to 0$ and use this to show $n_t \to \hat{n}$. Finally, I combine these results to show global convergence of $\nu_t \to \hat{\nu}$.

**Convergence of trait distribution**

Define $q_t := p_t/\hat{p}$. Using $p_t = q_t\hat{p}$ in the equation for $\dot{p}_t$ and subtracting the stationary equation for $\hat{p}$ multiplied by $q_t$, a short product–rule computation gives

$$\hat{p}\,\dot{q}_t = \tfrac{1}{2}\nabla^\top\big(\mathbf{M}\,\hat{p}\,\nabla q_t\big), \qquad \dot{q}_t = \frac{1}{2\,\hat{p}}\nabla^\top\big(\mathbf{M}\,\hat{p}\,\nabla q_t\big). \tag{44}$$

Using $\int f$ as shorthand for $\int_{\mathbb{R}^d} f(\mathbf{z})d\mathbf{z}$, a standard definition for relative entropy is

$$H(t) := H(p_t \mid \hat{p}) = \int p_t \ln\frac{p_t}{\hat{p}} = \int q_t \ln q_t\,\hat{p}. \tag{45}$$

Differentiate in time using the chain rule for $f(s) = s\ln s$, so $\frac{d}{ds}f(s) = 1 + \ln s$:

$$\frac{d}{dt}\big(q_t \ln q_t\big) = (1+\ln q_t)\,\dot{q}_t. \tag{46}$$

Hence $\dot{H}(t) = \int(1+\ln q_t)\,\dot{q}_t\,\hat{p}$. Now use the normalization $\int q_t\hat{p} = 1$ for all $t$. Differentiating this in time gives

$$0 = \frac{d}{dt}\int q_t\hat{p} = \int \dot{q}_t\,\hat{p}, \tag{47}$$

because $\hat{p}$ does not depend on $t$. Therefore $\int \dot{q}_t\,\hat{p} = 0$. Hence, $\dot{H}(t) = \int \dot{q}_t\,\hat{p}\,\ln q_t$. Substituting $\dot{q}_t$ and integrating by parts (boundary terms vanish by Gaussian tails of $p_t, \hat{p}$) gives

$$\dot{H}(t) = \frac{1}{2}\int \nabla^\top(\mathbf{M}\,\hat{p}\,\nabla q_t)\,\ln q_t = -\frac{1}{2}\int(\nabla q_t)^\top\mathbf{M}\,\nabla\ln q_t\,\hat{p}. \tag{48}$$

Using $\nabla q_t = q_t\,\nabla\ln q_t$,

$$\dot{H}(t) = -\frac{1}{2}\int q_t\,(\nabla\ln q_t)^\top\mathbf{M}\,\nabla\ln q_t\,\hat{p} \le 0, \tag{49}$$

since $\mathbf{M}$ is symmetric positive definite. Moreover $\dot{H}(t) = 0$ iff $\nabla\ln q_t \equiv 0$, and hence iff $q_t \equiv 1$ and $p_t = \hat{p}$. Thus $H(t)$ is non-increasing and constant only at $p_t = \hat{p}$. Under mild compactness/tightness assumptions (automatically satisfied here because both $p_t$ and $\hat{p}$ have Gaussian tails), any limit point $p_\infty$ of $p_t$ must satisfy $\dot{H} \equiv 0$ and hence $p_\infty = \hat{p}$. Therefore

$$\int |p_t - \hat{p}| \to 0 \text{ as } t \to \infty, \tag{50}$$

which quantifies the notion by which $p_t \to \hat{p}$.

**Convergence of abundance**

Note that

$$\bar{m}_t - \hat{\bar{m}} = \int_{\mathbb{R}^d}\big(m(\nu_t,\mathbf{z}) - m(\hat{\nu},\mathbf{z})\big)\,p_t(\mathbf{z})\,d\mathbf{z} + \int_{\mathbb{R}^d} m(\hat{\nu},\mathbf{z})\,\big(p_t(\mathbf{z}) - \hat{p}(\mathbf{z})\big)\,d\mathbf{z}. \tag{51}$$

Using the explicit quadratic form of $m$ and the convergence $p_t \to \hat{p}$ (which implies convergence of the relevant moments $\bar{\mathbf{z}}_t \to \hat{\bar{\mathbf{z}}}$ and $\mathbf{P}_t \to \hat{\mathbf{P}}$), both terms tend to 0, so $\bar{m}_t \to \hat{\bar{m}}$ as $t \to \infty$. Then for large enough $t$, $\bar{m}_t = \varepsilon_t - c(n_t - \hat{n})$ for some $\varepsilon_t \to 0$ as $t \to \infty$. But since $\hat{\bar{m}} = 0$, this implies $n_t \to \hat{n}$ as $t \to \infty$.

**Convergence of population density**

Taken together, we have

$$\int |\nu_t - \hat{\nu}| = \int \left|n_t p_t - \hat{n}\hat{p}\right| \leq |n_t - \hat{n}| + \hat{n} \int |p_t - \hat{p}| \to 0 \tag{52}$$

as $t \to \infty$ whenever $n_0 > 0$ and intrinsic growth $r$ is sufficiently large. Thus the Gaussian equilibrium $\hat{\nu}$ is globally asymptotically stable.

#### 3. Translating Between Continuous and Discrete Time Models

In this section, I describe how classical discrete time quantitative genetic models relate to the continuous time framework presented in the main text. Discrete time models work with fitness surfaces $W(\mathbf{z})$, which can be thought of as the average lifetime reproductive output of an individual with trait value $\mathbf{z}$ (however, as described in section 5 below, continuous-time individual-based models also utilize $W(\mathbf{z})$ so that its inclusion in a model does not imply generations are non-overlapping). Given a trait distribution $p(\mathbf{z})$, mean fitness is given by $\bar{W} = \int_{\mathbb{R}^d} W(\mathbf{z})p(\mathbf{z})d\mathbf{z}$. Then the trait distribution after a round of selection is given by $p^*(\mathbf{z}) = p(\mathbf{z})W(\mathbf{z})/\bar{W}$, and the post-selection mean trait is $\bar{\mathbf{z}}^* = \int_{\mathbb{R}^d} \mathbf{z}p^*(\mathbf{z})d\mathbf{z}$. A selection differential is then defined as $\mathbf{S} = \bar{\mathbf{z}}^* - \bar{\mathbf{z}}$, and a selection gradient as $\boldsymbol{\beta} = \mathbf{P}^{-1}\mathbf{S}$ (Lande and Arnold, 1983). Interfacing this approach with the imperfect inheritance model in the main text, the response to selection in discrete time is classically obtained as $\Delta\bar{\mathbf{z}} = \mathbf{G}\boldsymbol{\beta}$ (Lande, 1979).

To map results obtained using the discrete time approach to the continuous time setting presented here, I describe the formal connection between fitness surfaces $W$ and the growth rates $m$. Because there is a formal association between the two, in the main text I simply referred to $m$ as a fitness function. The basic idea is to begin with an individual-based description of a population, and then take a diffusion limit to arrive at a population-level description. This limiting process provides a formal way to calculate $m$ from $W$. I provide a brief overview of this process while omitting technical details, and then show how to use this approach to translate from classical fitness surfaces to continuous time growth rates.

Because $W(\mathbf{z})$ is an index associated with individuals that carry trait value $\mathbf{z}$, it inherently assumes an individual-based description of a population. This can be made more precise using the concept of a *point-mass* to represent an individual with trait value $\mathbf{z}$. In particular, the point-mass of an individual maps subsets $U$ of $\mathbb{R}^d$ to either 0 or 1 to indicate whether the individuals trait lies in $U$. That is, denoting $\mathbf{z}_i$ the trait value of the $i$-th individual in the population, the point-mass of this individual is defined by

$$\delta_{\mathbf{z}_i}(U) = \begin{cases} 1, & \mathbf{z}_i \text{ is in } U, \\ 0, & \mathbf{z}_i \text{ is not in } U. \end{cases} \tag{53}$$

For a population of $n$ individuals, the entire population can be summarized by the sum $N = \sum_{i=1}^n \delta_{\mathbf{z}_i}$. Hence $N(U) = \sum_{i=1}^n \delta_{\mathbf{z}_i}(U)$ returns the abundance of the sub-population taking trait values in $U \subset \mathbb{R}^d$, and $N(\mathbb{R}^d) = n$ in particular. This provides a flexible way to describe the distribution of traits in a population. In the main text, trait distributions are assumed to be multivariate normal. However, because the population described by $N$ is comprised of a sum of discrete individuals, the trait distribution cannot formally take the shape of a multivariate normal density. Even so, if the population size is very large, we can make histograms that summarize $N$ which appear like densities of continuous distributions. Mathematically, this connection is made precise by diffusion-limits (Feller, 1951). In this section I describe just one component of taking a diffusion-limit that involves rescaling individual contributions to population size. Rescaling this `weight" orunit`" of an individual is useful because we want to approximate continuous trait distributions while retaining a notion of finite population size. For the sake of clarity, other diffusion-limit details such as rescaling time are left out of this section. However, to my knowledge no sufficiently accessible description of the specific kind of diffusion-limit useful for constructing this framework has been published (in particular, one that prioritize intuition over rigor). I therefore provide an informal description of one in section 7 below.

A discussion of diffusion-limits in the univariate trait case is given by Week et al. (2021), and a classical introduction in the contexts of population ecology and population genetics is given by Karlin and Taylor (1981). A variety of these limiting approaches have been developed (e.g., see Champagnat et al., 2006). To be clear, not all of them involve rescaling individual "weights'', but as implied above I focus on one that does here. For concreteness, consider the state of a population of $n$ individuals at the initial time of the model, and follow the notation introduced above so that the population is summarized by $N = \sum_{i=1}^n \delta_{\mathbf{z}_i}$. The operation of rescaling individuals here involves replacing each individual by several other individuals whose weights sum to the weight of the individual they replace. One way to interpret this is to consider the trait values of individuals as uncertain. For instance, the trait value of individual $i$ may take values $\mathbf{z}_{i,1}, \mathbf{z}_{i,2}$ with probabilities one-half each. This leads to the rescaled population summarized by $N^2 = \frac{1}{2}\left(\sum_{i=1}^n \delta_{\mathbf{z}_{i,1}} + \sum_{i=1}^n \delta_{\mathbf{z}_{i,2}}\right)$. We can simplify indexing by collecting trait values of the rescaled individuals into the finite sequence $Z_2 = \{\mathbf{z}_1, \mathbf{z}_2, \dots, \mathbf{z}_{2n}\}$ with $\mathbf{z}_{2i} = \mathbf{z}_{i,1}$ (so the $\mathbf{z}_{i,1}$ are inserted into even slots of $Z_2$) and $\mathbf{z}_{2i-1} = \mathbf{z}_{i,2}$ (so the $\mathbf{z}_{i,2}$ are inserted into the odd slots of $Z_2$). More generally, write the $k$-th stage of rescaling as

$$N^k = \frac{1}{k}\sum_{i=1}^{nk} \delta_{\mathbf{z}_i}. \tag{54}$$

This approach to rescaling preserves the total abundance of the population because $N^k(\mathbb{R}^d) = n$ for each $k$. This is desired because we want to approximate continuous distributions of traits in the population while preserving the notion of population size.

Because we are free to choose criteria for how traits are added at each stage of rescaling, we can obtain any continuous initial distribution of traits as $k \to \infty$, but subsequent evolution of this distribution depends on how selection, mutation, etc., are modeled. In addition, depending on the biological determinants of individual fitness, the fitness surface associated with the $k$-th stage of rescaling $W_k(\mathbf{z})$ may depend on $k$. Explicit examples are given by Week et al. (2021), Week and Nuismer (2021), and Week and

Bradburd (2024). Additionally, as mentioned above, details involved with rescaling fitness are also presented in section 5 below. Furthermore, to account for frequency and density dependent selection, the fitness surface associated with the $k$-th rescaling of the population will also likely depend on the rescaled population summary $N^k$ so that $W_k(\mathbf{z})$ should be more formally replaced by $W_k(N^k, \mathbf{z})$. Under additional assumptions discussed for example in Champagnat et al. (2006), Week et al. (2021), and more formally by Méléard and Roelly (1993), the rescaled limit $k \to \infty$ allows for the formal calculation of the fitness function $m$ from the fitness surface $W$ as

$$m(\mathcal{N}, \mathbf{z}) = \lim_{k\to\infty} (W_k(N^k, \mathbf{z}) - 1)k, \tag{55}$$

where $N^k \to \mathcal{N}$. In particular, $\mathcal{N}$ summarizes the population in the diffusion-limit as $k \to \infty$. If we choose to add traits at each stage of rescaling $k$ by sampling them from a continuous distribution $p(\mathbf{z})$, then $\mathcal{N}$ will reflect this distribution in the sense that $\mathcal{N}(U)/n = \int_U p(\mathbf{z})d\mathbf{z}$. That is, unlike $N^k$, the limit $\mathcal{N}$ in this case cannot be written as a sum of point-masses. Under conditions discussed in Li (1998) this continuity of the trait distribution in the diffusion-limit is retained as the population responds to selection, mutation, and drift. This continuity implies the existence of a function $\nu(\mathbf{z})$ called the population density that satisfies $\mathcal{N}(U) = \int_U \nu(\mathbf{z})d\mathbf{z}$. In this case the diffusion-limit of the population is equally described by $\nu(\mathbf{z})$, and here it is convenient to replace $m(\mathcal{N}, \mathbf{z})$ with $m(\nu, \mathbf{z})$ to avoid the additional technical details required for working with the mapping $\mathcal{N}$ (formally, these details involve measure theory as presented by Axler, 2020).

If the rescaled fitness surface does not depend on the state of the population, and takes the special form of being the $k$-th root of $W$ at stage $k$ so that $W_k(N, \mathbf{z}) = W^{1/k}(\mathbf{z})$, then $m(\nu, \mathbf{z}) = m(\mathbf{z}) = \ln W(\mathbf{z})$, where $m(\mathbf{z})$ is introduced here because the resulting fitness function does not depend on the population density in the diffusion-limit $\nu$. This, for example, is the case for classical fitness surfaces (e.g., Lande, 1979; Jones et al., 2007; Arnold et al., 2008; Jones et al., 2012), such as the directional selection fitness surface

$$W_{\text{dir}}(\mathbf{z}) = W_0 e^{\mathbf{b}^\top \mathbf{z}}, \tag{56}$$

and the stabilizing selection fitness surface

$$W_{\text{stab}}(\mathbf{z}) = W_0 e^{-\frac{1}{2}(\boldsymbol{\theta}-\mathbf{z})^\top \boldsymbol{\Psi}(\boldsymbol{\theta}-\mathbf{z})}, \tag{57}$$

which are associated with the respective growth rate functions

$$m_{\text{dir}}(\mathbf{z}) = r + \mathbf{b}^\top \mathbf{z}, \tag{58}$$

$$m_{\text{stab}}(\mathbf{z}) = r - \tfrac{1}{2}(\boldsymbol{\theta} - \mathbf{z})^\top \boldsymbol{\Psi}\, (\boldsymbol{\theta} - \mathbf{z}), \tag{59}$$

where $r = \ln W_0$. Plugging these growth rates into equations (11b) and (11c) of the main text, ignoring both mutation and drift, and assuming multivariate normal trait distributions, we arrive at

$$\dot{\bar{\mathbf{z}}}_{\text{dir}} = \mathbf{G}\,\mathbf{b}, \quad \dot{\mathbf{G}}_{\text{dir}} = \mathbf{0}, \tag{60}$$

$$\dot{\bar{\mathbf{z}}}_{\text{stab}} = \mathbf{G}\,\boldsymbol{\Psi}\,(\boldsymbol{\theta} - \bar{\mathbf{z}}), \quad \dot{\mathbf{G}}_{\text{stab}} = -\mathbf{G}\,\boldsymbol{\Psi}\,\mathbf{G}. \tag{61}$$

Note, calculation of these expressions by applying a deterministic version of the framework (such as equations (16) and (17) in the main text) does not rely on weak selection approximations. In contrast, calculations involved with their discrete time counterparts often do (Jones et al., 2007, 2012).

#### 4. The Martingale Problem Approach

Here I outline an approach to formalize the framework and obtain the heuristics presented in the main text. Intuition for this approach comes from viewing populations as function spaces. This is formalized by viewing population processes as solutions to martingale problems, which is discussed at the end of the following subsection.

To communicate these ideas to as wide of audience as possible, most technical details are omitted and some are communicated with less accuracy than standard treatments. The goal here is to keep the discussion accessible to anyone familiar with the concepts of *norm*'s and *inner-products*'s on function spaces. It is assumed that readers familiar with the theory of measure-valued processes will be able to fill in *all* of the missing details, and notice where technically inaccurate statements are obviously stand-ins for their more accurate but technically involved counterparts. For readers unfamiliar with this theory, recommended entry points are suggested at the end of the following subsection.

##### 4.1. Formulation

For now, assume perfect inheritance and work directly with trait values $\mathbf{z} \in \mathbb{R}^d$. Under this approach, populations are summarized by finite *measures* on the trait space $\mathbb{R}^d$ (referred to as *mappings* in section 3 above). Measures are functions that map sets to numbers. In this context, a population measure is a function that maps subsets $U \subset \mathbb{R}^d$ to the abundance of the population in $U$. Here I focus on finite populations. So, if $\mathcal{N}_t$ is a measure representing a finite population on $\mathbb{R}^d$ at time $t$, then $\mathcal{N}_t(\mathbb{R}^d) < \infty$. On the other hand, if $\mathcal{N}_\tau(\mathbb{R}^d)$ is not finite for some $\tau > 0$, then consider only the time interval $[0, \tau)$. This is to handle cases where abundance can reach an infinite value in finite time. Biologically, this situation has been referred to as unstable persistence in models of mutualism (Wolin and Lawlor, 1984; Vandermeer and Boucher, 1978). Mathematically, reaching infinite values in finite time is called explosion (see sections 6.6 and 9.7 of Klebaner, 1998). However, once expressions for the framework are in place, one can examine the case where population size $\mathcal{N}_t(\mathbb{R}^d)$ is very large to obtain the Deterministic Covariance version and Deterministic Gradient version of the framework as deterministic approximations of the full stochastic framework.

To continue describing the foundations of the framework, we need to work with integrals of functions with respect to population measures. Because this is often not taught in courses that introduce norms and inner-products on function spaces, I briefly provide intuition for understanding these integrals. In particular, the integral of a function $x(\mathbf{z})$ over a set $U \subset \mathbb{R}^d$ with respect to the population measure $\mathcal{N}_t$ can be written as $\int_U x(\mathbf{z})\mathcal{N}_t(d\mathbf{z})$. Intuitively, one can think of this integral as a generalization of the classical Riemann integral $\int_U x(\mathbf{z})d\mathbf{z}$, but with $\mathcal{N}_t$ providing different weights to different parts of trait space. Taking this informal intuition further, we can partition $U$ into a disjoint set of subsets $\mathcal{U}_q = \{U_1, \dots, U_q\}$ such that $\cup_{i=1}^q U_i = U$, then pick a $\mathbf{z}_i \in U_i$ for each $i = 1, \dots, q$, and compute the sum $S_t(\mathcal{U}_q) = \sum_{i=1}^q x(\mathbf{z}_i)\mathcal{N}_t(U_i)$. Then assuming that, as $q \to \infty$, the partition $\mathcal{U}_q$ becomes more and more refined (similar to how sequences of partitions are used to define the multivariate Riemann integral), we can intuitively consider $\int_U x(\mathbf{z})\mathcal{N}_t(d\mathbf{z}) = \lim_{q\to\infty} S_t(\mathcal{U}_q)$. However, it should be carefully noted that the actual definition of the integral with respect to $\mathcal{N}_t$ depends on a concept known as Lebesgue integration (an excellent introduction is given by Axler, 2020), and there are important reasons why the intuitive approach outlined here fails in general.

For each time $t$, the population measure $\mathcal{N}_t$ defines a space of functions $x : \mathbb{R}^d \to \mathbb{R}$ having finite norm

$$(\!| x |\!)_t = \sqrt{\int_{\mathbb{R}^d} |x(\mathbf{z})|^2 \mathcal{N}_t(d\mathbf{z})} < \infty\,, \tag{62}$$

with inner-product $\langle\!\langle x, y \rangle\!\rangle_t = \int_{\mathbb{R}^d} x(\mathbf{z})y(\mathbf{z})\mathcal{N}_t(d\mathbf{z})$. Note the less conventional symbols $(\!| \cdot |\!)_t$ and $\langle\!\langle \cdot, \cdot \rangle\!\rangle_t$ are used to emphasize that their definitions differ from the symbols $\|\cdot\|$ and $\langle \cdot, \cdot \rangle$ used in the main text. These additional symbols are necessary because $\|\cdot\|$ and $\langle \cdot, \cdot \rangle$ depend on the dynamics of $\mathcal{N}_t$ (the reproductive variance $v$ in particular), but here these dynamics have not yet been defined. Computing $(\!| x |\!)_t$ for different functions returns information about the population. For example, computing $(\!| x |\!)_t^2$ with $x(\mathbf{z}) = 1$ returns the total abundance of the population at time $t$. For this reason, the functions satisfying equation (62) are often referred to as *test functions*. The relationship between different summary statistics of the population, obtained from using different test functions, can then be computed using the inner-product.

In the special case that $\mathcal{N}_t$ admits a density $\nu_t$, so that $\int_{\mathbb{R}^d} x(\mathbf{z})\mathcal{N}_t(d\mathbf{z}) = \int_{\mathbb{R}^d} x(\mathbf{z})\nu_t(\mathbf{z})d\mathbf{z}$ for *any* test function $x$, then the dynamics of the population process can be characterized by a stochastic partial differential equation (spde), such as

$$\dot{\nu}_t(\mathbf{z}) = m(\nu_t, \mathbf{z})\,\nu_t(\mathbf{z}) + \tfrac{1}{2}\,\nabla^\top \mathbf{M}\,\nabla\,\nu_t(\mathbf{z}) + \sqrt{v\,\nu_t(\mathbf{z})}\,\dot{W}_t(\mathbf{z})\,, \tag{63}$$

where $\dot{W}_t(\mathbf{z})$ is a space-time white noise process on $\mathbb{R}^d$. A nice introduction to spde's and space-time white noise is given by Dalang and Sanz-Solé (2024). Equation (63) is simply meant to provide an example of what a spde looks like. In particular, we might at first expect a population process similar to those studied in the main text to be characterized as a solution to equation (63).

In general, densities rarely exist for population processes, and this is especially true for traits with dimensionality greater than one (Dawson, 1993; Li, 1998; Etheridge, 2000; Perkins, 2002). As a consequence, the above spde is in general meaningless for multivariate traits. Hence, there is a need for a more general approach to characterize the population processes of interest here. Indeed, under very general conditions on the growth rate $m$, Méléard and Roelly (1993, p 109) have shown that the population process $\mathcal{N}_t$ can instead be characterized by

$$\mathcal{M}_t(x) := \int_{\mathbb{R}^d} x(\mathbf{z})\,\mathcal{N}_t(d\mathbf{z}) - \int_{\mathbb{R}^d} x(\mathbf{z})\,\mathcal{N}_0(d\mathbf{z}) - \int_0^t \int_{\mathbb{R}^d} \left(m(\mathcal{N}_s, \mathbf{z}) + \tfrac{1}{2}\nabla^\top \mathbf{M}\nabla\right) x(\mathbf{z})\,\mathcal{N}_s(d\mathbf{z})\,ds, \tag{64}$$

where $\mathcal{M}_t$ satisfies the martingale condition $\mathbb{E}[\,\mathcal{M}_{t+s}(x)\,|\,\mathcal{M}_t(x)\,] = \mathcal{M}_t(x)$ for *all* $x$ and $s > 0$. In particular, since $\mathcal{M}_0(x) = 0$, we have $\mathbb{E}[\,\mathcal{M}_t(x)\,] = 0$ (by the law of total expectation). Comparing with equation (64) above, the condition $\mathbb{E}[\,\mathcal{M}_t(x)\,] = 0$ is essentially a stochastic generalization of the fundamental theorem of calculus. Note that while the spde approach shows the magnitude of stochasticity explicitly as $\sqrt{v\,\nu_t(\mathbf{z})}$, the martingale characterization expresses this indirectly in terms of a *quadratic variation* discussed below. Details demonstrating the application of Méléard and Roelly (1993) to arrive at equation (64) above are given in Supplementary Material, Section 7 below.

Equation (64) can be rewritten using a differential notation shorthand, which is common shorthand for writing stochastic differential equations (for a discussion of this notational convention, see section 4.4 of Klebaner, 1998). To do so, first simplify notation by writing $\mathcal{N}_t(x) = \int_{\mathbb{R}^d} x(\mathbf{z})\mathcal{N}_t(d\mathbf{z})$ and $\mathcal{L}(\mathcal{N}_t, \mathbf{z}) = m(\mathcal{N}_t, \mathbf{z}) + \frac{1}{2}\nabla^\top \mathbf{M}\nabla$ (or simply $\mathcal{L}$). Note that the population measure $\mathcal{N}_t$ is first defined as a mapping from subsets of $\mathbb{R}^d$ to numbers. Here, with a common notational abuse, the same symbol is used to denote the induced mapping from functions to real numbers via integration against this measure. Both notations refer to the same underlying population object, and the intended interpretation is clear from context. Then, for each test function $x$, (64) can be rewritten as an ordinary stochastic differential equation. This expression is not derived through algebraic manipulation, but is a compact notational representation of the integral martingale formulation above.

$$d\mathcal{N}_t(x) = \mathcal{N}_t(\mathcal{L}x)\,dt + d\mathcal{M}_t(x)\,. \tag{65}$$

Equation (65) is particularly useful for placing the framework on a mathematically rigorous basis. For instance, setting $x(\mathbf{z}) = 1$ returns the stochastic differential equation (17) for the abundance of the population presented in the main text. Furthermore, the above expression already allows us to establish a simplified verion of the *additive property* of the heuristics presented in Table 1 of the main text. Replacing $x$ with $ax + by$ in equation (64) and then rewriting in the form of (65) shows that $d\mathcal{M}_t(ax + by) =$

$a\, d\mathcal{M}_t(x) + b\, d\mathcal{M}_t(y)$. To see why, observe that each integral on the right-hand-side of equation (64) is linear with respect to $x$, and hence $\mathcal{M}_t(ax+by) = a\,\mathcal{M}_t(x) + b\,\mathcal{M}_t(y)$ (note the lack of $d$'s in this expression). Rewriting in differential notation shorthand then establishes the result. The more general version of the additive property allows $a$ and $b$ to be stochastic processes (typically univariate phenotypic moments), which is justified in section 4.2 below. I return to an expression similar to (65) in section 5, and use it to derive the dynamical equations presented in the main text.

Importantly, for each test function $x(\mathbf{z})$, the martingale $\mathcal{M}_t(x)$ is associated with a so-called *quadratic variation*, which provides insights into its variance. Again following Méléard and Roelly (1993, p 109), the quadratic variation of $\mathcal{M}_t(x)$ is given by

$$[\,\mathcal{M}(x)\,]_t := v \int_0^t \int_{\mathbb{R}^d} |x(\mathbf{z})|^2 \mathcal{N}_s(d\mathbf{z}) ds = v \int_0^t (\!| x |\!)_s^2 ds\,. \tag{66}$$

This quadratic variation will serve as the basis for establishing the heuristics presented in the main text, which allow for manipulating stochastic terms and deriving analytical models. In particular, the above quadratic variation illustrates how to compute products of stochastic terms that share a common test function $x$ (i.e., $d\mathcal{M}_t(x)\, d\mathcal{M}_t(x)$). Then, to establish heuristics for computing products of stochastic terms based on different test functions $x$ and $y$ (i.e., $d\mathcal{M}_t(x)\, d\mathcal{M}_t(y)$), I introduce the following bilinear form, referred to as the *quadratic covariation* (Rogers and Williams, 2000a) and is a special case of the result obtained by Roelly-Coppoletta (1986, p 177):

$$[\,\mathcal{M}(x),\, \mathcal{M}(y)\,]_t = v \int_0^t \int_{\mathbb{R}^d} x(\mathbf{z}) y(\mathbf{z}) \mathcal{N}_s(d\mathbf{z}) ds = v \int_0^t \langle\!\langle x, y \rangle\!\rangle_s ds\,. \tag{67}$$

**What's the problem?** This is called a *martingale problem* approach because the population process $\mathcal{N}_t$ in equation (64) can be viewed as an unknown. Very roughly, the problem is then to find a process $\mathcal{N}_t$ such that $\mathcal{M}_t(x)$ is a martingale for each $x$. As a consequence, this solution features the quadratic variation and quadratic covariation described by equations (66)–(67) (Roelly-Coppoletta, 1986; Méléard and Roelly, 1993). However, the technical prerequisites for specifying the martingale problem more precisely are outside the scope of this supplementary material. Further information can be found in Stroock and Varadhan (1997), where the martingale problem approach is fully developed in the context of finite-dimensional systems of stochastic differential equations. For the present context, an infinite-dimensional extension is required, and this has been treated by Dawson (1993). An introduction to measure-valued population processes is provided by Etheridge (2000). A comprehensive introduction to stochastic calculus is given by Rogers & Williams (2000b; 2000a). More concise, but still advanced introductions are given by Le Gall (2016), Bass (2011), and Durret (1996). Introductions focused on stochastic differential equations are given by Øksendal (2013) and Evans (2012). More elementary (and therefore extremely useful!) introductions are given by Klebaner (1998) and Calin (2015). An introduction in the context of population genetics is given by Etheridge (2011).

With expressions for the quadratic variation $[\,\mathcal{M}(x)\,]_t$ and the quadratic covariation $[\,\mathcal{M}(x),\, \mathcal{M}(y)\,]_t$ in hand, we are now ready to develop the heuristics introduced in the main text, which is the subject of the following section.

---

#### 4.2. Heuristics

Below I present justification for the heuristics presented in Table 1 of the main text. To facilitate comparisons with the main text, I adopt the norm $\|\cdot\|$ and inner-product $\langle\cdot,\cdot\rangle$ used there. These are related to the norm $(\!|\cdot|\!)_t$ and inner-product $\langle\!\langle\cdot,\cdot\rangle\!\rangle_t$ introduced above via $\|\, x\,\|_t = \sqrt{v}\,(\!| x |\!)_t$ and $\langle\, x,\, y\,\rangle_t = v\,\langle\!\langle\, x,\, y\,\rangle\!\rangle_t$ (which hold for test functions $x, y$). Note that in the main text subscripts indexing time are ommitted to minimize notational complexity, but are needed here due to the appearance of integrals across intervals of time.

##### *4.2.1. The Scaling Property*

A simplified version of the *scaling property* states that $d\mathcal{M}_t(\hat{x}) := d\mathcal{M}_t(x)/\|\,x\,\|_t$ is the stochastic differential of a standard Brownian motion (where "standard" just means it starts at zero and has variance $t$ at time $t$). Notation-wise, it is worth noting that $\mathcal{M}_t(x)$ is technically defined for fixed test functions $x$, but $\hat{x}_t(\mathbf{z}) = x(\mathbf{z})/\|\,x\,\|_t$ is not a fixed function. This is the reason why the symbol $d\mathcal{M}_t(\hat{x})$ has to be defined as above, and the benefit of this convention will be clear when combining the additive and multiplicative properties while deriving the stochastic framework below. The more general version of the scaling property (which states that, for $q_t(\mathbf{z})$ a stochastic process that is also a random test function for each fixed $t$ that may also depend on $\mathcal{N}_t$, $d\mathcal{M}_t(\hat{q}) := d\mathcal{M}_t(q)/\|\,q\,\|_t$ is the stochastic differential of a standard Brownian motion) will be made sense of at the end of this justification.

Formally, the scaling property may be justified by the representation of continuous martingales as integrals with respect to Brownian motion (integration with respect to Brownian motion is introduced in Chapter 4 of Evans, 2012). More precisely, Corollary 14.5.4 on page 359 of Cohen and Elliot (2015) justifies the integral representation $\mathcal{M}_t(x) = \int_0^t \|\,x\,\|_s dB_s$, where integration is with respect to a standard Brownian motion $B_t$ (i.e., $B_0 = 0$ and the variance of $B_t$ is $t$). In differential notation shorthand, we have $d\mathcal{M}_t(x) = \|\,x\,\|_t dB_t$. Then, informally dividing each side by $\|\,x\,\|_t$ gives the result. To finish the formal justification, use integration with respect to the martingale $\mathcal{M}_t(x)$ (integration with respect to martingales is introduced in Chapter 10 of Bass, 2011) to define the process $B'_t := \int_0^t \|\,x\,\|_s^{-1} d\mathcal{M}_s(x)$. Here, the meaning of $d\mathcal{M}_t(x)$ extends beyond its interpretation as merely shorthand for writing integral expressions. Theorem 30.7 on page 55 of Rogers and Williams (2000a) implies that $B'_t$ is a continuous martingale with associated quadratic variation given by

$$[B']_t = \int_0^t \frac{1}{\|\,x\,\|_s^2} d[\mathcal{M}(x)]_s\,. \tag{68}$$

Writing equation (66) in differential form provides $d[\mathcal{M}(x)]_s = \|\,x\,\|_s^2\, ds$. Hence, $[B']_t = \int_0^t ds = t$. In addition, because $B'_0 = 0$, we can apply the Lévy characterization of Brownian motion (e.g., Theorem 8.6.1 on page 162 of Øksendal, 2013) to conclude that $B'_t$ is a standard Brownian motion.

The notation used for the scaling property can be generalized by setting $d\mathcal{M}_t(a_t x) := a_t d\mathcal{M}_t(x)$ for a test function $x$ and a univariate process $a_t$ (i.e., one that takes values in $\mathbb{R}$). This notational convention will be needed to justify the multiplicative and additive properties below. More precisely, the process $a_t$ is assumed to take the form $a_t = f(\mathcal{N}_t)$. For instance, $f(\mathcal{N}_t) = \mathcal{N}_t(z_i)/\mathcal{N}_t(1)$ is the $i$-th component of the mean trait $\bar{z}_i$ at time $t$. By Theorem 30.7 on page 55 of Rogers and Williams (2000a), if $f$ satisfies $\int_0^t f^2(\mathcal{N}_s)\| x \|_s^2 ds < \infty$ for each $t > 0$, then $\mathcal{Q}_t(f,x) := \int_0^t f(\mathcal{N}_s)d\mathcal{M}_s(x) = \int_0^t a_s d\mathcal{M}_s(x)$ is a continuous martingale with quadratic variation

$$[\mathcal{Q}(f,x)]_t = \int_0^t f^2(\mathcal{N}_s)d[\mathcal{M}(x)]_s. \tag{69}$$

Furthermore, Equation 30.8 on page 56 of Rogers and Williams (2000a) states that the martingales $\mathcal{Q}_t(f_1,x)$ and $\mathcal{Q}_t(f_2,y)$ will have quadratic covariation

$$[\mathcal{Q}(f_1,x),\mathcal{Q}(f_2,y)]_t = \int_0^t f_1(\mathcal{N}_s)f_2(N_s)d[\mathcal{M}(x),\mathcal{M}(y)]_s, \tag{70}$$

where $d[\mathcal{M}(x),\mathcal{M}(y)]_t = \langle x, y \rangle_t \, dt$ is differential notation shorthand for the same statement expressed by equation (67) above.

With this convention the scaling property is obtained for a given test function $x$ by setting $f(\mathcal{N}_t) = 1/\sqrt{v\mathcal{N}_t(|x|^2)}$ so that $a_t = \| x \|_t^{-1}$. Furthermore, it should be carefully noted that, for many choices of $f$, $d\mathcal{M}_t(a_t x)$ will not be the stochastic differential of a Brownian motion.

Finally, the more general scaling property that states $d\mathcal{M}_t(\hat{q}) := d\mathcal{M}_t(q)/\| q \|_t$ is the stochastic differential of a standard Brownian motion (where $q_t(\mathbf{z})$ is a stochastic process such that, for each fixed $t$, $q_t$ is a test function that may also depend on $\mathcal{N}_t$) is justified by first defining the notation $\int_{\mathbb{R}^d} x(\mathbf{z})\mathcal{M}(dt,d\mathbf{z}) := d\mathcal{M}_t(x)$. Here $\mathcal{M}$ is still used for $\mathcal{M}(dt,d\mathbf{z})$, even though it is now technically a measure on $\mathbb{R}_{\geq 0} \times \mathbb{R}^d$, because it still represents the same fundamental underlying object as $\mathcal{M}_t$. Then we can write $\mathcal{Q}_t(f,x)$ as

$$\mathcal{Q}_t(f,x) = \int_0^t \int_{\mathbb{R}^d} f(\mathcal{N}_s)x(\mathbf{z})\mathcal{M}(ds,d\mathbf{z}). \tag{71}$$

Integrals of this form are natural generalizations of integration with respect to martingales, and are formally treated in Chapter 2 of Walsh (1986). What is important here is that we can now define

$$\mathcal{Q}_t(q) := \int_0^t \int_{\mathbb{R}^d} q_s(\mathbf{z})\mathcal{M}(ds,d\mathbf{z}), \tag{72}$$

which, setting $\|q\|_t := \sqrt{v\int_{\mathbb{R}^d} q_t^2(\mathbf{z})\mathcal{N}_t(d\mathbf{z})}$, requires $\mathbb{E}\left[\int_0^t \|q\|_s^2 ds\right] < \infty$ for each $t > 0$. Given this condition, $\mathcal{Q}_t(q)$ is again a continuous martingale. Following Chapter 2 of Walsh (1986), we then have the quadratic covariation between $\mathcal{Q}_t(q^{(1)})$ and $\mathcal{Q}_t(q^{(2)})$, for two processes $q_t^{(1)}(\mathbf{z}), q_t^{(2)}(\mathbf{z})$, is given by

$$[\mathcal{Q}(q^{(1)}),\mathcal{Q}(q^{(2)})]_t = \int_0^t \langle q^{(1)}, q^{(2)} \rangle_s ds, \tag{73}$$

where $\langle q^{(1)}, q^{(2)} \rangle_t = v\int_{\mathbb{R}^d} q_t^{(1)}(\mathbf{z})q_t^{(2)}(\mathbf{z})\mathcal{N}_t(d\mathbf{z})$. From here, define $d\mathcal{M}_t(q) := d\mathcal{Q}_t(q)$. Then, following the same arguments for $B_t'$ above, we have $\mathcal{Q}_t' := \int_0^t d\mathcal{M}_s(\hat{q}) = \int_0^t \|q\|_s^{-1} d\mathcal{M}_s(q)$ is a continuous martingale with quadratic variation

$$[\mathcal{Q}']_t = \int_0^t \frac{1}{\|q\|_s^2} d[\mathcal{Q}_t(q)] = \int_0^t ds = t. \tag{74}$$

So Lévy's characterization of Brownian motion again justifies the general scaling property.

Lastly, for a scalar-valued process $a_t$, define $a_t d\mathcal{M}_t(q) := d\mathcal{Q}_t(aq)$ so that $a_t d\mathcal{M}_t(q) = d\mathcal{M}_t(a_t q)$.

. . . . . . . . . . . . . . . . . . . . . . . . . . . . . . . . . . . . . . . . . . . . . . . . . . . . . . . . .

###### *4.2.2.* ***The Multiplicative Property***

The multiplicative property states $d\mathcal{M}_t(q^{(1)})\, d\mathcal{M}_t(q^{(2)}) = \langle q^{(1)}, q^{(2)} \rangle_t \, dt$, where $q_t^{(i)}(\mathbf{z})$ is a process that, for each fixed $t$, is also a (random) test function, and $\langle q^{(1)}, q^{(2)} \rangle_t = v\int_{\mathbb{R}^d} q_t^{(1)}(\mathbf{z})q_t^{(2)}(\mathbf{z})\mathcal{N}_t(d\mathbf{z})$. Formal justification for this property is based on the general definition of the quadratic covariation process (for example, given by Definition 31.3 on page 58 of Rogers and Williams, 2000a). Here I provide justification that leverages known results for the quadratic covariation process. To begin, consider the simplified situation where $q_t^{(1)}(\mathbf{z}) = x(\mathbf{z})$ and $q_t^{(2)}(\mathbf{z}) = y(\mathbf{z})$ are fixed test functions. The differential notation used by the multiplicative property then simplifies to a convenient shorthand defined by $d\mathcal{M}_t(x)\, d\mathcal{M}_t(y) := d[\,\mathcal{M}(x),\, \mathcal{M}(y)\,]_t$, where $d[\,\mathcal{M}(x),\, \mathcal{M}(y)\,]_t$ is already informal shorthand based on $[\,\mathcal{M}(x),\, \mathcal{M}(y)\,]_t$. For the special case studied here (determined by equations (64) and (67)), we have that $d[\,\mathcal{M}(x),\, \mathcal{M}(y)\,]_t = \langle x, y \rangle_t \, dt$ is shorthand to express the same statement as equation (67) above. The reasoning behind this differential shorthand is clarified by seeing where it occurs in Itô's formula, which serves as a generalization of the chain rule for stochastic calculus. In particular, the differential notation shorthand establishes general heuristics to facilitate calculations involved with applying Itô's formula. I therefore paraphrase the statement of Itô's formula in the following paragraph, which applies more generally to a finite collection of martingales $\{U_t^1, \ldots, U_t^k\}$ (i.e., $U_t^i$ is a stochastic process taking values in $\mathbb{R}$ such that $\mathbb{E}[U_t^i \,|\, U_s^i] = U_s^i$ for $s < t$). The multiplicative property presented in Table 1 of the main text is then based on the special case where these martingales are of the form $U_t^i = \mathcal{M}_t(x_i)$, with $\mathcal{M}_t$ defined by equations (64) and (67) and test functions $x_1, \ldots, x_k$.

Following Theorem 32.8 on page 60 of Rogers and Williams (2000a), given a $k$-dimensional process $U_t = (U_t^1, \ldots, U_t^k)$, where each component $U_t^i$ is a martingale, and a function $F(u)$ with continuous second derivatives $\partial_i \partial_j F$ (where $\partial_i = \partial/\partial u^i$), Itô's formula states (in differential notation shorthand)

$$dF(U_t) = \sum_{i=1}^{k} \partial_i F(U_t)\, dU_t^i + \frac{1}{2} \sum_{i,j=1}^{k} \partial_i \partial_j F(U_t)\, d[\, U^i,\, U^j\,]_t. \tag{75}$$

With this result in place, heuristical calculations involved with its application are facilitated by defining $dU_t^i\, dU_t^j := d[\, U^i,\, U^j\,]_t$. Setting $U_t^i = \mathcal{M}_t(x_i)$ and $U_t^j = \mathcal{M}_t(x_j)$ establishes the simplified multiplicative property where $q_t^{(1)}(\mathbf{z}) = x(\mathbf{z})$ and $q_t^{(2)}(\mathbf{z}) = y(\mathbf{z})$ are fixed test functions. The general case is established by setting $U_t^i = \mathcal{Q}_t(q^{(i)})$ and $U_t^j = \mathcal{Q}_t(q^{(j)})$, where the definition of $\mathcal{Q}_t(q) := \int_0^t \int_{\mathbb{R}^d} q_s(\mathbf{z}) \mathcal{M}(ds, d\mathbf{z})$ is given by equation (72). This definition formalizes the notation $d\mathcal{M}_t(q^{(i)}) = d\mathcal{Q}_t(q^{(i)})$. Then, writing equation (73) above in differential notation shorthand justifies the multiplicative property in the most general form needed to derive the stochastic framework from the martingale problem approach.

. . . . . . . . . . . . . . . . . . . . . . . . . . . . . . . . . . . . . . . . . . . . . . . . . . . . . . . . . . . . .

###### *4.2.3.* ***The Additive Property***

As mentioned in section 4.1, a simplified version of the additive property $d\mathcal{M}_t(ax + by) = a\, d\mathcal{M}_t(x) + b\, d\mathcal{M}_t(y)$ (which holds for constant scalars $a, b$ and test functions $x(\mathbf{z}), y(\mathbf{z})$) is justified by combining equation (64) with the differential notation shorthand introduced in equation (65). In particular, the linearity of the integrals appearing in equation (64) in addition to the linearity of the operator $\mathcal{L}(\mathcal{N}_t, \mathbf{z}) = m(\mathcal{N}_t, \mathbf{z}) + \frac{1}{2}\nabla^\top \mathbf{M} \nabla$ implies $\mathcal{M}_t(ax + by) = a\, \mathcal{M}_t(x) + b\, \mathcal{M}_t(y)$. Then rewriting using the differential notation shorthand establishes this simplified additive property in the case where $a, b$ are scalars.

For the more general case of processes $q_t(\mathbf{z})$ that, for each $t$, are test functions, most of the work to justify the additive property comes down to relating the notational conventions introduced here. For instance, recall the definition $d\mathcal{M}_t(q) := d\mathcal{Q}_t(q)$ (with $\mathcal{Q}_t$ defined by equation (72) above). Following Chapter 2 of Walsh (1986), we have that $\mathcal{Q}_t$ is linear such that $\mathcal{Q}_t(aq^{(1)} + bq^{(2)}) = a\mathcal{Q}_t(q^{(1)}) + b\mathcal{Q}_t(q^{(2)})$. In addition, for a scalar-valued process $a_t$, the convention $a_t d\mathcal{M}_t(q) := d\mathcal{M}_t(aq) := d\mathcal{Q}_t(aq)$ is justified because $a_t q_t(\mathbf{z})$ is again a process that is a (random) test function for each fixed $t$. Hence, if $b_t$ is another scalar-valued process, a generalized version of the additive property states $d\mathcal{M}_t(aq^{(1)} + bq^{(2)}) = a_t d\mathcal{M}_t(q^{(1)}) + b_t d\mathcal{M}_t(q^{(2)})$.

#### 5. Derivation of The Stochastic Framework from The Martingale Problem Approach

To obtain the Brownian Motion Gradient version ($BG$) and the Martingale Gradient version ($MG$), I begin in section 5.1 by interfacing the martingale problem approach with the model of imperfect inheritance presented in section 2.1.3 of the main text. After this, I present the derivation for abundance dynamics in section 5.2. In section 5.3 I derive mean trait dynamics and show the noise processes driving them do not covary with abundance dynamics. Lastly, in section 5.4 I derive additive genetic variance dynamics and show its noise process does not covary with that of abundance dynamics, and also that it does not covary with the noise process driving mean trait dynamics when traits are multivariate normal.

##### 5.1. Imperfect Inheritance

The population measure of additive genetic values is denoted by $\mathcal{G}_t$. From this the population measure of trait values $\mathcal{N}_t$ is the measure satisfying

$$\int_{\mathbb{R}^d} x(\mathbf{z})\, \mathcal{N}_t(d\mathbf{z}) = \int_{\mathbb{R}^d} \int_{\mathbb{R}^d} x(\mathbf{g} + \mathbf{e})\, \mathcal{G}_t(d\mathbf{g}) \varepsilon(\mathbf{e}) d\mathbf{e}, \tag{76}$$

for each function $x$, with $\varepsilon(\mathbf{e})$ the density of phenotypic residuals. Following Lande (1979), I assume residuals are multivariate normally distributed with mean $\mathbf{0}$ and covariance matrix $\mathbf{E}$. As a short-hand, write $\mathcal{N}_t = \mathcal{G}_t * \varepsilon$.

To model imperfect inheritance, I define $\mathcal{M}_t$ in terms of $\mathcal{G}_t$ instead of $\mathcal{N}_t$. This is slightly different than simply replacing $\mathcal{N}_t$ with $\mathcal{G}_t$ in equation (64) of Supplementary Material, Section 2.1 because fitness is still determined by expressed trait values. So I define fitness of additive genetic values as

$$\tilde{m}(\mathcal{G}_t, \mathbf{g}) = \int_{\mathbb{R}^d} m(\mathcal{N}_t, \mathbf{g} + \mathbf{e}) \varepsilon(\mathbf{e}) d\mathbf{e} = \int_{\mathbb{R}^d} m(\mathcal{G}_t * \varepsilon, \mathbf{g} + \mathbf{e}) \varepsilon(\mathbf{e}) d\mathbf{e}, \tag{77}$$

and define $\tilde{\mathcal{L}}(\mathcal{G}_t, \mathbf{g})\, x(\mathbf{g}) = \tilde{m}(\mathcal{G}_t, \mathbf{g})\, x(\mathbf{g}) + \frac{1}{2}\nabla^\top \mathbf{M} \nabla\, x(\mathbf{g})$, with gradients taken with respect to $\mathbf{g}$. Then, using the notation $\mathcal{G}_t(x) = \int_{\mathbb{R}^d} x(\mathbf{g})\, \mathcal{G}_t(d\mathbf{g})$, the replacement for equation (64) that accommodates imperfect inheritance is given by

$$\mathcal{M}_t(x) = \mathcal{G}_t(x) - \mathcal{G}_0(x) - \int_0^t \mathcal{G}_s(\tilde{\mathcal{L}} x) ds \tag{78}$$

This defines the martingale process referred to in the main text. For each $x$, this process can be expressed as a differential equation:

$$d\mathcal{G}_t(x) = \mathcal{G}_t(\tilde{\mathcal{L}} x)\, dt \,+\, d\mathcal{M}_t(x). \tag{79}$$

To keep the following sections concise, I omit subscripts denoting time dependence and use the heuristic notation presented in Table 1 of the main text. Note that the heuristics presented in the main text assume that $\mathcal{G}$ admits the population density $\gamma(\mathbf{g})$ such that the distribution of additive genetic values has a density $\tilde{p}(\mathbf{g})$ (and thus $\gamma(\mathbf{g}) = n\, \tilde{p}(\mathbf{g})$). In the calculations below, this assumption is only necessary when multivariate normality is assumed. However, for the purposes here, nothing is gained by treating the more general case where $\mathcal{G}$ does not necessarily admit a density. Instead, the more general case requires the use of notation that differs from the heuristics presented in the main text. Hence, for the sake of clarity, the following calculations are presented under the assumption that $\mathcal{G}(x) = n\int_{\mathbb{R}^d} x(\mathbf{g})\tilde{p}(\mathbf{g})d\mathbf{g}$. In addition, the symbol $\tilde{m}(\gamma,\mathbf{g})$ will be written in place of $\tilde{m}(\mathcal{G},\mathbf{g})$, and similarly $\tilde{\mathcal{L}}(\gamma,\mathbf{g})$ will be written in place of $\tilde{\mathcal{L}}(\mathcal{G},\mathbf{g})$. For the sake of brevity, these will often be written simply as $\tilde{m}$ and $\tilde{\mathcal{L}}$.

---

#### 5.2. Total Abundance $n$

Because $\mathcal{G}(1) = n\int_{\mathbb{R}^d} 1\tilde{p}(\mathbf{g})d\mathbf{g} = n$ and $\tilde{\mathcal{L}}1 = \tilde{m}$ (recalling $\tilde{\mathcal{L}}(\gamma,\mathbf{g}) = \tilde{m}(\gamma,\mathbf{g}) + \frac{1}{2}\nabla^{\top}\mathbf{M}\nabla$) which gives $\mathcal{G}(\tilde{\mathcal{L}}1) = n\int_{\mathbb{R}^d}(\tilde{\mathcal{L}}1)\tilde{p}(\mathbf{g})d\mathbf{g} = n\int_{\mathbb{R}^d}\tilde{m}(\gamma,\mathbf{g})\tilde{p}(\mathbf{g})d\mathbf{g} = \bar{m}\,n$ (recall from equation (14) in the main text that $\bar{m} = \bar{\tilde{m}}$), equation (79) can be used to write

$$dn = \bar{m}\,n\,dt + d\mathcal{M}(1). \tag{80}$$

This provides the expression of abundance dynamics in the $MG$ version of the framework (equation (26c) in the main text). Using the scaling property obtained in Supplementary Material, Section 4.2, we immediately have $d\mathcal{M}(1) = \|1\|\,dB_n$ where $B_n := \mathcal{M}(\hat{1})$ is a standard normal Brownian motion, and by definition $\|1\| = \sqrt{v\,n}\sqrt{\int_{\mathbb{R}^d} 1^2\tilde{p}(\mathbf{g})d\mathbf{g}} = \sqrt{v\,n}$. Thus,

$$dn = \bar{m}\,n\,dt + \sqrt{v\,n}\,dB_n. \tag{81}$$

This provides the expression for abundance dynamics presented in the $BG$ version of the framework (equation (18) in the main text). Neither of these expressions for abundance dynamics require the multivariate normal trait assumption.

---

#### 5.3. Mean Trait Vector $\bar{\mathbf{z}}$

##### *5.3.1. Martingale Gradient Version*

Setting $x(\mathbf{g}) = g_i$, equation (79) immediately shows $d\mathcal{G}(g_i) = \mathcal{G}(\tilde{\mathcal{L}}g_i)dt + d\mathcal{M}_t(g_i)$. Because $\tilde{\mathcal{L}}g_i = \tilde{m}\,g_i$ ($x(\mathbf{g}) = g_i$ has no curvature), we have $\mathcal{G}(\tilde{\mathcal{L}}g_i) = n\int_{\mathbb{R}^d}(\tilde{\mathcal{L}}g_i)\tilde{p}(\mathbf{g})d\mathbf{g} = n\int_{\mathbb{R}^d}\tilde{m}\,g_i\tilde{p}(\mathbf{g})d\mathbf{g} = n\,\overline{\tilde{m}\,g_i} = n\,(\mathrm{Cov}(\tilde{m},g_i) + \bar{m}\,\bar{g}_i)$. Combining this with $\mathring{g}_i := \mathcal{G}(g_i) = n\int_{\mathbb{R}^d} g_i\tilde{p}(\mathbf{g})d\mathbf{g}$, we arrive at

$$d\mathring{g}_i = d\mathcal{G}(g_i) = n\,(\mathrm{Cov}(\tilde{m},g_i) + \bar{m}\bar{g}_i)\,dt + d\mathcal{M}(g_i). \tag{82}$$

Because $\mathring{g}_i = \mathcal{G}(g_i) = n\int_{\mathbb{R}^d} g_i\tilde{p}(\mathbf{g})d\mathbf{g} = n\,\bar{g}_i$, we then have $\bar{g}_i = \mathring{g}_i/n$. Itô's quotient rule (provided, for example, at the bottom of page 140 in Calin, 2015) immediately states

$$d\left(\frac{\mathring{g}_i}{n}\right) = \frac{1}{n}\left(d\mathring{g}_i - \frac{\mathring{g}_i}{n}dn - \frac{1}{n}d\mathring{g}_i\,dn + \frac{\mathring{g}_i}{n^2}(dn)^2\right). \tag{83}$$

Using our results for $d\mathring{g}_i$ and $dn$, we have

$$d\mathring{g}_i\,dn = (n\,(\mathrm{Cov}(\tilde{m},g_i) + \bar{m}\bar{g}_i)\,dt + d\mathcal{M}(g_i))\,(\bar{m}\,n\,dt + d\mathcal{M}(1))\,. \tag{84}$$

Expanding the product and then applying the general heuristics $dt^2 = 0$ and $dt\,d\mathcal{M}(x) = 0$ shows $d\mathring{g}_i\,dn = d\mathcal{M}(g_i)\,d\mathcal{M}(1)$. The multiplicative property immediately gives $d\mathcal{M}(1)\,d\mathcal{M}(g_i) = v\,n\int_{\mathbb{R}^d} g_i\tilde{p}(\mathbf{g})d\mathbf{g}\,dt = v\,n\,\bar{g}_i\,dt$. Hence, $d\mathring{g}_i\,dn = v\,n\,\bar{g}_i\,dt$.

We also have $(dn)^2 = (\bar{m}\,n\,dt + d\mathcal{M}(1))^2$. Expanding the square and then applying the general heuristics $dt^2 = 0$ and $dt\,d\mathcal{M}(x) = 0$ shows $(dn)^2 = (d\mathcal{M}(1))^2$. The multiplicative property immediately gives $(d\mathcal{M}(1))^2 = v\,n\int_{\mathbb{R}^d} 1\tilde{p}(\mathbf{g})d\mathbf{g}\,dt = v\,n\,dt$. Hence, $(dn)^2 = v\,n\,dt$.

. . . . . . . . . . . . . . . . . . . . . . . . . . . . . . . . . . . . . . . . . . . . . . . . . . . . . . . . .

Because $\bar{g}_i = \mathring{g}_i/n$, we immediately have (with no intermediate steps to show) $d\bar{g}_i = d(\mathring{g}_i/n)$. Combining this with our results for $d\mathring{g}_i\,dn$ and $(dn)^2$ shows

$$d\bar{g}_i = \frac{1}{n}\left(d\mathring{g}_i - \frac{\mathring{g}_i}{n}dn - \frac{1}{n}(v\,n\,\bar{g}_i\,dt) + \frac{\mathring{g}_i}{n^2}(v\,n\,dt)\right). \tag{85}$$

Substituting $\mathring{g}_i = n\,\bar{g}_i$ and simplifying shows

$$d\bar{g}_i = \frac{1}{n}\left(d\mathring{g}_i - \bar{g}_i\,dn - (v\,\bar{g}_i\,dt) + (v\,\bar{g}_i\,dt)\right). \tag{86}$$

Because $-(v\,\bar{g}_i\,dt) + (v\,\bar{g}_i\,dt) = 0$, we then have

$$d\bar{g}_i = \frac{1}{n}\left(d\mathring{g}_i - \bar{g}_i\, dn\right). \tag{87}$$

Substituting in our results for $d\mathring{g}_i$ and $dn$ shows

$$d\mathring{g}_i - \bar{g}_i\, dn = n\left(\mathrm{Cov}(\tilde{m}, g_i) + \bar{m}\bar{g}_i\right) dt + d\mathcal{M}(g_i) - \bar{g}_i\left(\bar{m}\, n\, dt + d\mathcal{M}(1)\right) = n\,\mathrm{Cov}(\tilde{m}, g_i)\, dt + d\mathcal{M}(g_i) - \bar{g}_i\, d\mathcal{M}(1). \tag{88}$$

By the additive property we have $d\mathcal{M}(g_i) - \bar{g}_i\, d\mathcal{M}(1) = d\mathcal{M}(g_i) - d\mathcal{M}(\bar{g}_i) = d\mathcal{M}(g_i - \bar{g}_i)$. Substituting these results into the expression for $d\bar{g}_i$ leads to

$$d\bar{g}_i = \mathrm{Cov}(\tilde{m}, g_i)dt + \tfrac{1}{n} d\mathcal{M}(g_i - \bar{g}_i). \tag{89}$$

Assuming multivariate normality, equation (20) in index notation immediately gives $\mathrm{Cov}(\tilde{m}, g_i) = \sum_j G_{ij}(\partial_j\, \bar{m} - \overline{\partial_j\, \tilde{m}})$, where $\partial_i = \partial/\partial\bar{z}_i = \partial/\partial\bar{g}_i$ because $\bar{e}_i = 0$ (and thus, $\bar{z}_i = \bar{g}_i$). Combining these last three results gives

$$d\bar{z}_i = \sum_j G_{ij}(\partial_j\, \bar{m} - \overline{\partial_j\, \tilde{m}})dt + \tfrac{1}{n} d\mathcal{M}(g_i - \bar{g}_i), \tag{90}$$

By definition

$$\overline{\partial_j\, \tilde{m}} = \int_{\mathbb{R}^d} \partial_j \tilde{m}(\gamma, \mathbf{g})\tilde{p}(\mathbf{g})d\mathbf{g} = \int_{\mathbb{R}^d} \partial_j \int_{\mathbb{R}^d} m(\gamma, \mathbf{g} + \mathbf{e})\varepsilon(\mathbf{e})d\mathbf{e}\tilde{p}(\mathbf{g})d\mathbf{g}. \tag{91}$$

Using the change of variables $\mathbf{g} = \mathbf{z} - \mathbf{e}$, we obtain

$$\overline{\partial_j\, \tilde{m}} = \int_{\mathbb{R}^d} \partial_j \int_{\mathbb{R}^d} m(\nu, \mathbf{z})\varepsilon(\mathbf{e})\tilde{p}(\mathbf{z} - \mathbf{e})d\mathbf{e}d\mathbf{z} = \int_{\mathbb{R}^d} \partial_j m(\nu, \mathbf{z}) \int_{\mathbb{R}^d} \varepsilon(\mathbf{e})\tilde{p}(\mathbf{z} - \mathbf{e})d\mathbf{e}d\mathbf{z}. \tag{92}$$

But $p(\mathbf{z}) = \int_{\mathbb{R}^d} \varepsilon(\mathbf{e})\tilde{p}(\mathbf{z} - \mathbf{e})d\mathbf{e}$. We thus have $\overline{\partial_j\, \tilde{m}} = \int_{\mathbb{R}^d} \partial_j m(\nu, \mathbf{z}) \int_{\mathbb{R}^d} p(\mathbf{z})d\mathbf{z} = \overline{\partial_j\, m}$. Finally, plugging this into equation (90) gives

$$d\bar{z}_i = \sum_j G_{ij}(\partial_j\, \bar{m} - \overline{\partial_j\, m})dt + \tfrac{1}{n} d\mathcal{M}(g_i - \bar{g}_i), \tag{93}$$

which is the mean trait dynamics under the Martingale Gradient version ($MG$) of the framework expressed using index notation (i.e., equation (26b) of the main text).

. . . . . . . . . . . . . . . . . . . . . . . . . . . . . . . . . . . . . . . . . . . . . . . . . . . . . . . . . . .

##### 5.3.2. *Brownian Motion Gradient Version*

The Brownian Motion Gradient version ($BG$) is obtained by first using the scaling property which states that we can write

$$d\mathcal{M}(g_i - \bar{g}_i) = \|g_i - \bar{g}_i\| d\mathcal{M}(\widehat{g_i - \bar{g}_i}), \tag{94}$$

with $B_{\bar{z}_i} := \mathcal{M}(\widehat{g_i - \bar{g}_i})$ a standard Brownian motion and $\|g_i - \bar{g}_i\| = \sqrt{v\, n \int_{\mathbb{R}^d} (g_i - \bar{g}_i)^2 \tilde{p}(\mathbf{g})d\mathbf{g}} = \sqrt{v\, n\, G_{ii}}$. Then, mean trait dynamics under the Brownian Motion Gradient version in index notation are given by

$$d\bar{z}_i = \sum_j G_{ij}(\partial_j\, \bar{m} - \overline{\partial_j\, m})dt + \sqrt{\tfrac{v}{n} G_{ii}}dB_{\bar{z}_i}, \tag{95}$$

where carefully note that the $\sqrt{n}$ in the numerator of $\sqrt{v\, n\, G_{ii}}$ interacts with the $n$ in the denominator appearing in equation (93) to leave a $\sqrt{n}$ in the denominator of equation (95).

. . . . . . . . . . . . . . . . . . . . . . . . . . . . . . . . . . . . . . . . . . . . . . . . . . . . . . . . . . .

**Sphering Vector-Valued Martingales**

The stochastic dynamics of the mean trait components are not independent, and their correlated dynamics are quantified using the multiplicative property. In particular, while $B_{\bar{z}_i}$ is a standard Brownian motion, it is not in general independent of $B_{\bar{z}_j}$. Collect these non-independent standard Brownian motions into a vector $\mathring{\mathbf{B}}_{\bar{\mathbf{z}}} = (B_{\bar{z}_1}, \ldots, B_{\bar{z}_d})^\top$. Then, by definition of integration with respect to Brownian motion, we have

$$\mathring{\mathbf{B}}_{\bar{\mathbf{z}}}(t) = \int_0^t d\mathring{\mathbf{B}}_{\bar{\mathbf{z}}}(s) \tag{96}$$

with $d\mathring{\mathbf{B}}_{\bar{\mathbf{z}}} = (dB_{\bar{z}_1}, \ldots, dB_{\bar{z}_d})^\top$. Intuitively, we can think about $dB_{\bar{z}_i}$ as being normally distributed with mean zero and variance $dB_{\bar{z}_i}^2 = dt$ (where this last equality is just a standard heuristic from stochastic calculus). More generally, we can think about $d\mathring{\mathbf{B}}_{\bar{\mathbf{z}}}$ as multivariate normal with mean zero and covariance matrix given by the outer-product

$$\mathbb{E}\left(d\mathring{\mathbf{B}}_{\bar{\mathbf{z}}} d\mathring{\mathbf{B}}_{\bar{\mathbf{z}}}^\top\right) = \mathbb{E}\begin{pmatrix} dB_{\bar{z}_1}^2 & dB_{\bar{z}_1}dB_{\bar{z}_2} & \cdots & dB_{\bar{z}_1}dB_{\bar{z}_d} \\ dB_{\bar{z}_2}dB_{\bar{z}_1} & dB_{\bar{z}_2}^2 & \cdots & dB_{\bar{z}_2}dB_{\bar{z}_d} \\ \vdots & \vdots & \ddots & \vdots \\ dB_{\bar{z}_d}dB_{\bar{z}_1} & dB_{\bar{z}_d}dB_{\bar{z}_2} & \cdots & dB_{\bar{z}_d}^2 \end{pmatrix}. \tag{97}$$

The standard heuristic $dB_{\bar{z}_i}^2 = dt$ shows that the diagonal entries of this matrix are equal to $dt$. The off-diagonals, which quantify correlations between the dynamics of the entries of $\mathring{\mathbf{B}}_{\bar{\mathbf{z}}}$, can be computed using the multiplicative property because $dB_{\bar{z}_i} :=$

$d\mathcal{M}(\widehat{g_i - \bar{g}_i})$. The multiplicative property (in combination with the scaling property) then states

$$dB_{\bar{z}_i} dB_{\bar{z}_j} = d\mathcal{M}(\widehat{g_i - \bar{g}_i}) d\mathcal{M}(\widehat{g_j - \bar{g}_j}) = \frac{d\mathcal{M}(g_i - \bar{g}_i)}{\|g_i - \bar{g}_i\|} \frac{d\mathcal{M}(g_j - \bar{g}_j)}{\|g_j - \bar{g}_j\|} \tag{98}$$

$$= \frac{\langle g_i - \bar{g}_i, g_j - \bar{g}_j \rangle}{v\, n \sqrt{G_{ii} G_{jj}}} dt = \frac{v\, n \int_{\mathbb{R}^d} (g_i - \bar{g}_i)(g_j - \bar{g}_j) \tilde{p}(\mathbf{g}) d\mathbf{g}}{v\, n \sqrt{G_{ii} G_{jj}}} dt = \frac{G_{ij}}{\sqrt{G_{ii} G_{jj}}} dt. \tag{99}$$

Hence, the dynamics of the Brownian motions $B_{\bar{z}_i}$ and $B_{\bar{z}_j}$ covary according to the genetic correlation between traits $z_i$ and $z_j$: $\rho_{ij} = G_{ij}/\sqrt{G_{ii} G_{jj}}$. Note that, although covariance matrices are formally defined using an expectation (e.g., $\mathbb{E}(d\mathring{\mathbf{B}}_{\bar{\mathbf{z}}} d\mathring{\mathbf{B}}_{\bar{\mathbf{z}}}^\top)$), the informal argument developed here leverages heuristics where products of stochastic differentials are deterministic quantities (e.g., $dB_{\bar{z}_i}^2 = dt$). So the expectation is not needed to express heuristical covariance matrices such as $d\mathring{\mathbf{B}}_{\bar{\mathbf{z}}} d\mathring{\mathbf{B}}_{\bar{\mathbf{z}}}^\top$, and thus will be dropped to simplify notation.

Building on this intuitive perspective, we can leverage a so-called *sphering transform* common in data analysis (reviewed by Kessy et al., 2018) to use the matrix $d\mathring{\mathbf{B}}_{\bar{\mathbf{z}}} d\mathring{\mathbf{B}}_{\bar{\mathbf{z}}}^\top$ to rewrite $d\mathring{\mathbf{B}}_{\bar{\mathbf{z}}}$ it terms of a vector of uncorrelated standard Brownian motions $\mathbf{B}_{\bar{\mathbf{z}}}$. The main idea here is that, for a $d$-dimensional multivariate normal vector $\mathbf{v}$ with covariance matrix $\mathbf{\Sigma}$, the new vector $\mathbf{v}' = \sqrt{\mathbf{\Sigma}}^{-1} \mathbf{v}$ (where $\sqrt{\mathbf{\Sigma}}$ is any $d \times d$ invertible matrix that satisfies $\sqrt{\mathbf{\Sigma}} \sqrt{\mathbf{\Sigma}}^\top = \mathbf{\Sigma}$, which exists if $\mathbf{\Sigma}$ is positive definite) is also multivariate normal but now with identity covariance matrix $\mathbf{I}$. Conversely, we can start with a multivariate normal vector $\mathbf{v}'$ with covariance matrix $\mathbf{I}$, then the new vector $\mathbf{v} = \sqrt{\mathbf{\Sigma}} \mathbf{v}'$ will be multivariate normal with covariance matrix $\mathbf{\Sigma}$ (where here $\sqrt{\mathbf{\Sigma}}$ need not be invertible and $\mathbf{\Sigma} = \sqrt{\mathbf{\Sigma}} \sqrt{\mathbf{\Sigma}}^\top$ can be singular). This application of the matrix $\sqrt{\mathbf{\Sigma}}$ has been called a covariance-shaping transform (the inverse of the ZCA-Mahalanobis sphering transform in particular, see Kessy et al., 2018).

Setting $\mathbf{R} = d\mathring{\mathbf{B}}_{\bar{\mathbf{z}}} d\mathring{\mathbf{B}}_{\bar{\mathbf{z}}}^\top / dt$ (dividing by $dt$ because $dB_{\bar{z}_i} dB_{\bar{z}_j} = \rho_{ij} dt$) and defining $\mathbf{B}_{\bar{\mathbf{z}}}$ as a $d$-dimensional vector of independent standard Brownian motions, we have

$$\mathring{\mathbf{B}}_{\bar{\mathbf{z}}}(t) = \int_0^t \sqrt{\mathbf{R}(\mathbf{s})}\, d\mathbf{B}_{\bar{\mathbf{z}}}(s), \tag{100}$$

and hence $d\mathring{\mathbf{B}}_{\bar{\mathbf{z}}} = \sqrt{\mathbf{R}}\, d\mathbf{B}_{\bar{\mathbf{z}}}$. As an immediate consequence of equation (95), the mean trait dynamics can be written in matrix form as

$$d\bar{\mathbf{z}} = \mathbf{G}(\nabla_{\bar{\mathbf{z}}} \bar{m} - \overline{\nabla_{\bar{\mathbf{z}}} m}) dt + \sqrt{\tfrac{v}{n}} \sqrt{\mathbf{D_G}}\, d\mathring{\mathbf{B}}_{\bar{\mathbf{z}}}, \tag{101}$$

where $\sqrt{\mathbf{D_G}}$ is the $d \times d$ diagonal matrix with $ii$-th entry equal to $\sqrt{G_{ii}}$ and $ij$-th entry equal to zero when $i \neq j$. Plugging in our sphering transform result $d\mathring{\mathbf{B}}_{\bar{\mathbf{z}}} = \sqrt{\mathbf{R}}\, d\mathbf{B}_{\bar{\mathbf{z}}}$, the dynamics of $\bar{\mathbf{z}}$ is given in terms of the vector of independent standard Brownian motions $\mathbf{B}_{\bar{\mathbf{z}}}$ as

$$d\bar{\mathbf{z}} = \mathbf{G}(\nabla_{\bar{\mathbf{z}}} \bar{m} - \overline{\nabla_{\bar{\mathbf{z}}} m}) dt + \sqrt{\tfrac{v}{n}} \sqrt{\mathbf{D_G}} \sqrt{\mathbf{R}}\, d\mathbf{B}_{\bar{\mathbf{z}}}. \tag{102}$$

Setting $\mathbf{A} = \sqrt{\mathbf{D_G}} \sqrt{\mathbf{R}}$ we have

$$\mathbf{A}\mathbf{A}^\top = (\sqrt{\mathbf{D_G}} \sqrt{\mathbf{R}})(\sqrt{\mathbf{D_G}} \sqrt{\mathbf{R}})^\top = \sqrt{\mathbf{D_G}}\, \mathbf{R} \sqrt{\mathbf{D_G}}. \tag{103}$$

Using index notation, the $ij$-th entry of this product is given by $(\mathbf{A}\mathbf{A}^\top)_{ij} = \sum_{kl} (\sqrt{\mathbf{D_G}})_{ik} \mathbf{R}_{kl} (\sqrt{\mathbf{D_G}})_{ik}$. Because $\sqrt{\mathbf{D_G}}$ is diagonal such that $(\sqrt{\mathbf{D_G}})_{ij} = \sqrt{G_{ij}} \delta_{ij}$ with $\delta_{ii} := 1$ and $\delta_{ij} := 0$ when $i \neq j$, the sum simplifies to $(\mathbf{A}\mathbf{A}^\top)_{ij} = \sqrt{G_{ii}} \mathbf{R}_{ij} \sqrt{G_{jj}}$. Recalling that $\mathbf{R}_{ij} = \rho_{ij} = G_{ij}/\sqrt{G_{ii} G_{jj}}$, we see $\mathbf{A}\mathbf{A}^\top = \mathbf{G}$. Then, setting $\sqrt{\mathbf{G}} := \mathbf{A}$ we arrive at equation (19) in the main text:

$$d\bar{\mathbf{z}} = \mathbf{G}(\nabla_{\bar{\mathbf{z}}} \bar{m} - \overline{\nabla_{\bar{\mathbf{z}}} m}) dt + \sqrt{\tfrac{v}{n}} \sqrt{\mathbf{G}}\, d\mathbf{B}_{\bar{\mathbf{z}}}. \tag{104}$$

It turns out we can set $\sqrt{\mathbf{G}} := \mathbf{A}$ for any $d \times d$ matrix $\mathbf{A}$ that satisfies $\mathbf{A}\mathbf{A}^\top = \mathbf{G}$ (for a discussion, see section 5.1 of Durrett, 1996). As a final note regarding the intuitive sphering approach outlined above, formal justification requires handling $\mathbf{G}$ as a stochastic process (which can be done using notions of quadratic covariation for martingales to represent martingales as integrals with respect to Brownian motions). However, under this formalization, nothing fundamentally changes here other than how mathematical objects are interpreted. In particular, the result for expressing $d\bar{\mathbf{z}}$ in terms of $\mathbf{B}_{\bar{\mathbf{z}}}$ holds after formalizing all the details.

. . . . . . . . . . . . . . . . . . . . . . . . . . . . . . . . . . . . . . . . . . . . . . . . . . . . . . . . . . . . .

##### 5.3.3. *Uncorrelated noise processes driving* $n, \bar{\mathbf{z}}$

Lastly, to show the noise process driving mean trait dynamics is uncorrelated from the noise process driving abundance dynamics, expand the product $dn\, d\bar{z}_i$ and apply the general heuristics $dt^2 = dt\, d\mathcal{M}(x) = 0$ to arrive at $dn\, d\bar{z}_i = \frac{1}{n} d\mathcal{M}(1)\, d\mathcal{M}(g_i - \bar{g}_i)$. Using the multiplicative property shows $d\mathcal{M}(1)\, d\mathcal{M}(g_i - \bar{g}_i) = \langle 1, g_i - \bar{g}_i \rangle\, dt = v\, n \int_{\mathbb{R}^d} (g_i - \bar{g}_i) \tilde{p}(\mathbf{g}) d\mathbf{g}\, dt = 0\, dt = 0$, which holds for any distribution of additive genetic values $\mathbf{g}$. This calculation immediately implies $dB_n\, dB_{\bar{z}_i} = 0$ (based on the definitions of $dB_n$ and $dB_{\bar{z}_i}$). Then, leveraging the same intuitive perspective used to justify the sphering argument above, we see that the noise processes driving $n$ and $\bar{z}_i$ are uncorrelated for any distribution of additive genetic values.

##### 5.4. Additive Genetic Covariance Matrix G

###### 5.4.1. Martingale Gradient Version

Set $\mathring{G}_{ij} := \mathcal{G}(g_i g_j)$. Recalling that $\mathcal{G}(x) = n\int_{\mathbb{R}^d} x(\mathbf{g})\tilde{p}(\mathbf{g})d\mathbf{g}$, we then have $\mathring{G}_{ij} = n\int_{\mathbb{R}^d} g_i g_j \tilde{p}(\mathbf{g})d\mathbf{g} = n\,\overline{g_i g_j} = n\,(G_{ij} + \bar{g}_i\bar{g}_j)$ (because $\mathrm{Cov}(g_i, g_j) = G_{ij}$). Equation (79) states $d\mathring{G}_{ij} = \mathcal{G}(\tilde{\mathcal{L}}(g_i g_j))dt + d\mathcal{M}(g_i g_j)$. Because $\tilde{\mathcal{L}} = \tilde{m} + \frac{1}{2}\nabla^\top\mathbf{M}\nabla$, we have

$$\tilde{\mathcal{L}}(g_i g_j) = \tilde{m} g_i g_j + \tfrac{1}{2}\nabla^\top\mathbf{M}\nabla(g_i g_j) = \tilde{m} g_i g_j + \tfrac{1}{2}\sum_{kl} M_{kl}\partial_{g_k}\partial_{g_l}(g_i g_j), \tag{105}$$

where the summation on the right is just the definition of $\nabla^\top\mathbf{M}\nabla$ (see equation (2) in the main text). Using the product rule from elementary calculus, we find $\partial_{g_l}(g_i g_j) = \delta_{il} g_j + \delta_{jl} g_i$, with $\delta_{ii} := 1$ and $\delta_{ij} := 0$ when $i \neq j$. In addition, $\partial_{g_k}\partial_{g_l}(g_i g_j) = \delta_{il}\delta_{jk} + \delta_{jl}\delta_{ik}$. Substituting into the summation we find $\frac{1}{2}\nabla^\top\mathbf{M}\nabla(g_i g_j) = \frac{1}{2}(M_{ij} + M_{ji})$. Symmetry of $\mathbf{M}$ then demonstrates $\tilde{\mathcal{L}}(g_i g_j) = \tilde{m} g_i g_j + M_{ij}$. So $\mathcal{G}(\tilde{\mathcal{L}}(g_i g_j)) = n\int_{\mathbb{R}^d} \tilde{m}(g_i g_j + M_{ij})\tilde{p}(\mathbf{g})d\mathbf{g} = n\,(\overline{\tilde{m} g_i g_j} + M_{ij}) = n\,(\mathrm{Cov}(\tilde{m}, g_i g_j) + \bar{m}\,\overline{g_i g_j} + M_{ij})$. Hence,

$$d\mathring{G}_{ij} = n\,(M_{ij} + \mathrm{Cov}(\tilde{m}, g_i g_j) + \bar{m}\,\overline{g_i g_j})\,dt + d\mathcal{M}(g_i g_j). \tag{106}$$

Itô's quotient rule immediately states (see page 140 of Calin, 2015)

$$d\left(\frac{\mathring{G}_{ij}}{n}\right) = \frac{d\mathring{G}_{ij}}{n} - \frac{\mathring{G}_{ij}}{n}\frac{dn}{n} - \frac{d\mathring{G}_{ij}}{n}\frac{dn}{n} + \frac{\mathring{G}_{ij}}{n}\frac{dn^2}{n^2}. \tag{107}$$

Substituting the above result for $d\mathring{G}_{ij}$ and the expression for $dn$ given by equation (56) into the product $d\mathring{G}_{ij}\,dn$, then expanding the product and applying the general heuristics $dt^2 = dt\,d\mathcal{M}(x) = 0$, we obtain $d\mathring{G}_{ij}\,dn = d\mathcal{M}(g_i g_j)\,d\mathcal{M}(1)$. The multiplicative property states $d\mathcal{M}(g_i g_j)\,d\mathcal{M}(1) = \langle g_i g_j, 1\rangle\,dt = vn\int_{\mathbb{R}^d} g_i g_j \tilde{p}(\mathbf{g})d\mathbf{g}\,dt = v\,n\,\overline{g_i g_j}\,dt$. So $d\mathring{G}_{ij}\,dn = v\,n\,\overline{g_i g_j}\,dt$. We also have (obtained just below equation (60) above) $dn^2 = v\,n\,dt$. Then substituting these results in addition to $\overline{g_i g_j} = \mathring{G}_{ij}/n$ (which was obtained at the beginning of this subsection) and the result for $d\mathring{G}_{ij}$ into equation (107), and cancelling the $n$'s that appear in both numerator and denominator, we arrive at (with no additional steps other than the stated substitutions and cancelling of $n$'s)

$$d\overline{g_i g_j} = \left((M_{ij} + \mathrm{Cov}(\tilde{m}, g_i g_j) + \bar{m}\,\overline{g_i g_j})\,dt + \frac{1}{n}d\mathcal{M}(g_i g_j)\right) - \overline{g_i g_j}\left(\bar{m}\,dt + \frac{1}{n}d\mathcal{M}(1)\right) - \frac{v}{n}\overline{g_i g_j}\,dt + \overline{g_i g_j}\frac{v}{n}dt. \tag{108}$$

Simplifying by canceling out the terms that sum to zero, and using the additive property which immediately shows $d\mathcal{M}(g_i g_j) - \overline{g_i g_j}d\mathcal{M}(1) = d\mathcal{M}(g_i g_j - \overline{g_i g_j})$, we arrive at

$$d\overline{g_i g_j} = (M_{ij} + \mathrm{Cov}(\tilde{m}, g_i g_j))dt + \frac{1}{n}d\mathcal{M}(g_i g_j - \overline{g_i g_j}). \tag{109}$$

. . . . . . . . . . . . . . . . . . . . . . . . . . . . . . . . . . . . . . . . . . . . . . . . . . . . . . . . . . .

Consider the function $f(x, y, z) = x - yz$ (where $x, y, z$ are just scalar variables here, not to be confused with test functions or trait values). Itô's formula (e.g., equation (4.73) on page 114 of Klebaner, 1998) applied to $f(X_t, Y_t, Z_t)$ states

$$\begin{aligned} df(X_t, Y_t, Z_t) = {} & [\partial_x f(X_t, Y_t, Z_t)]dX_t + [\partial_y f(X_t, Y_t, Z_t)]dY_t + [\partial_z f(X_t, Y_t, Z_t)]dZ_t \\ & + \frac{1}{2}\left(\partial_x^2 f(X_t, Y_t, Z_t)]dX_t^2 + [\partial_y^2 f(X_t, Y_t, Z_t)]dY_t^2 + [\partial_z^2 f(X_t, Y_t, Z_t)]dZ_t^2\right) \\ & + (\partial_x\partial_y f(X_t, Y_t, Z_t)]dX_t dY_t + [\partial_x\partial_z f(X_t, Y_t, Z_t)]dX_t dY_t + [\partial_y\partial_z f(X_t, Y_t, Z_t)]dY_t dZ_t). \end{aligned} \tag{110}$$

For the specific functional form $f(x, y, z) = x - yz$, we then have

$$df(X_t, Y_t, Z_t) = dX_t - Z_t dY_t - Y_t dZ_t - dY_t dZ_t. \tag{111}$$

Note that $G_{ij} = f(\overline{g_i g_j}, \bar{g}_i, \bar{g}_j) = \overline{g_i g_j} - \bar{g}_i\bar{g}_j$ (this is an immediate consequence of the definition of covariance). Applying Itô's formula for $f(x, y, z)$ to $dG_{ij} = df(\overline{g_i g_j}, \bar{g}_i, \bar{g}_j)$ provides

$$dG_{ij} = d\overline{g_i g_j} - \bar{g}_j d\bar{g}_i - \bar{g}_i d\bar{g}_j - d\bar{g}_i\,d\bar{g}_j. \tag{112}$$

Substituting our result for $d\overline{g_i g_j}$ and the result for $d\bar{g}_i$ given in equation (89) into the expression for $dG_{ij}$ provides

$$\begin{aligned} dG_{ij} = {} & (M_{ij} + \mathrm{Cov}(\tilde{m}, g_i g_j))dt + \tfrac{1}{n}d\mathcal{M}(g_i g_j - \overline{g_i g_j}) - \bar{g}_j\left(\mathrm{Cov}(\tilde{m}, g_i)dt + \tfrac{1}{n}d\mathcal{M}(g_i - \bar{g}_i)\right) \\ & - \bar{g}_i\left(\mathrm{Cov}(\tilde{m}, g_j)dt + \tfrac{1}{n}d\mathcal{M}(g_j - \bar{g}_j)\right) - d\bar{g}_i\,d\bar{g}_j. \end{aligned} \tag{113}$$

Using $dt^2 = dt\,d\mathcal{M}(x) = 0$, expand the product $d\bar{g}_i\,d\bar{g}_j$ and simplify to find $d\bar{g}_i\,d\bar{g}_j = d\mathcal{M}(g_i - \bar{g}_i)d\mathcal{M}(g_j - \bar{g}_j)/n^2$. The multiplicative property shows $d\mathcal{M}(g_i - \bar{g}_i)d\mathcal{M}(g_j - \bar{g}_j) = v\,n\int_{\mathbb{R}^d}(g_i - \bar{g}_i)(g_j - \bar{g}_j)\tilde{p}(\mathbf{g})d\mathbf{g}\,dt = v\,n\,G_{ij}\,dt$. In addition, bilinearity of covariance justifies $\mathrm{Cov}(\tilde{m}, g_i g_j) - \bar{g}_j\mathrm{Cov}(\tilde{m}, g_i) - \bar{g}_i\mathrm{Cov}(\tilde{m}, g_j) = \mathrm{Cov}(\tilde{m}, g_i g_j - \bar{g}_j g_i - \bar{g}_i g_j)$. Because $\mathrm{Cov}(\tilde{m}, \bar{g}_i\bar{g}_j) = 0$ (covariance with a constant is zero), we have $\mathrm{Cov}(\tilde{m}, g_i g_j - \bar{g}_j g_i - \bar{g}_i g_j) = \mathrm{Cov}(\tilde{m}, g_i g_j - \bar{g}_j g_i - \bar{g}_i g_j + \bar{g}_i \bar{g}_j) = \mathrm{Cov}(\tilde{m}, (g_i - \bar{g}_i)(g_j - \bar{g}_j))$. Using the additive property, we also have

$$d\mathcal{M}(g_i g_j - \overline{g_i g_j}) - \bar{g}_j d\mathcal{M}(g_i - \bar{g}_i) - \bar{g}_i d\mathcal{M}(g_j - \bar{g}_j) = d\mathcal{M}(g_i g_j - \overline{g_i g_j} - \bar{g}_j g_i + \bar{g}_i \bar{g}_j - \bar{g}_i g_j + \bar{g}_i \bar{g}_j) = d\mathcal{M}((g_i - \bar{g}_i)(g_j - \bar{g}_j) - G_{ij}), \tag{114}$$

where the right equality leveraged $\overline{g_i g_j} = G_{ij} + \bar{g}_i \bar{g}_j$. Plugging these results in to our expression for $dG_{ij}$ provides

$$dG_{ij} = (M_{ij} + \mathrm{Cov}(\tilde{m}, (g_i - \bar{g}_i)(g_j - \bar{g}_j)) - \tfrac{v}{n} G_{ij})dt + \frac{1}{n} d\mathcal{M}((g_i - \bar{g}_i)(g_j - \bar{g}_j) - G_{ij}). \tag{115}$$

Assuming multivariate normality, equation (22) in section 2.2 directly shows

$$dG_{ij} = (M_{ij} + 2\sum_{kl} G_{ik}(\partial_{kl}\tilde{m} - \overline{\partial_{kl}\tilde{m}})G_{jl} - \tfrac{v}{n} G_{ij})dt + \frac{1}{n} d\mathcal{M}((g_i - \bar{g}_i)(g_j - \bar{g}_j) - G_{ij}). \tag{116}$$

Finally, the justification given above equation (93) can be extended to justify $\overline{\partial_{kl}\tilde{m}} = \overline{\partial_{kl} m}$, which then justifies equation (26c) in the main text (the Martingale Gradient version).

. . . . . . . . . . . . . . . . . . . . . . . . . . . . . . . . . . . . . . . . . . . . . . . . . . . . . . . .

##### *5.4.2. **Brownian Motion Gradient Version***

For the Brownian Motion Gradient version, the scaling property shows $d\mathcal{M}\big((g_i - \bar{g}_i)(g_j - \bar{g}_j) - G_{ij}\big) = \|(g_i - \bar{g}_i)(g_j - \bar{g}_j) - G_{ij}\| dB_{G_{ij}}$ with $dB_{G_{ij}}$ a standard Brownian motion. By definition, we also have

$$\|(g_i - \bar{g}_i)(g_j - \bar{g}_j) - G_{ij}\| = \sqrt{v\, n \int_{\mathbb{R}^d} ((g_i - \bar{g}_i)(g_j - \bar{g}_j) - G_{ij})^2 \tilde{p}(\mathbf{g}) d\mathbf{g}}. \tag{117}$$

Expanding the square leads to

$$\sqrt{v\, n \int_{\mathbb{R}^d} ((g_i - \bar{g}_i)(g_j - \bar{g}_j) - G_{ij})^2 \tilde{p}(\mathbf{g}) d\mathbf{g}} = \sqrt{v\, n \left(\overline{(g_i - \bar{g}_i)^2 (g_j - \bar{g}_j)^2} - G_{ij}^2\right)}. \tag{118}$$

Assuming multivariate normality, Isserlis' theorem (Isserlis, 1918) immediately implies

$$\overline{(g_i - \bar{g}_i)(g_j - \bar{g}_j)(g_k - \bar{g}_k)(g_l - \bar{g}_l)} = G_{ij}G_{kl} + G_{ik}G_{jl} + G_{il}G_{jk}. \tag{119}$$

Applying Isserlis' theorem then provides the simplification

$$\|(g_i - \bar{g}_i)(g_j - \bar{g}_j) - G_{ij}\| = \sqrt{v\, n}\sqrt{G_{ii}G_{jj} + G_{ij}^2}. \tag{120}$$

Together this justifies equation (23) in the main text (from the Brownian Motion Gradient version of the framework). Carefully note, however, that the standard Brownian motions $B_{G_{ij}}$ and $B_{G_{kl}}$ will in general covary even under the assumption of multivariate normality. More generally, the multiplicative property shows

$$\begin{aligned} &d\mathcal{M}((g_i - \bar{g}_i)(g_j - \bar{g}_j) - G_{ij})\, d\mathcal{M}((g_k - \bar{g}_k)(g_l - \bar{g}_l) - G_{kl}) = \langle (g_i - \bar{g}_i)(g_j - \bar{g}_j) - G_{ij}, (g_k - \bar{g}_k)(g_l - \bar{g}_l) - G_{kl} \rangle\, dt \\ &= v\, n \int_{\mathbb{R}^d} ((g_i - \bar{g}_i)(g_j - \bar{g}_j) - G_{ij})((g_k - \bar{g}_k)(g_l - \bar{g}_l) - G_{kl}) \tilde{p}(\mathbf{g}) d\mathbf{g}\, dt = v\, n \left(\overline{(g_i - \bar{g}_i)(g_j - \bar{g}_j)(g_k - \bar{g}_k)(g_l - \bar{g}_l)} - G_{ij}G_{kl}\right) dt. \end{aligned} \tag{121}$$

Applying Isserlis' theorem again gives the simplification

$$d\mathcal{M}((g_i - \bar{g}_i)(g_j - \bar{g}_j) - G_{ij})\, d\mathcal{M}((g_k - \bar{g}_k)(g_l - \bar{g}_l) - G_{kl}) = v\, n\, (G_{ik}G_{jl} + G_{il}G_{jk}) dt. \tag{122}$$

Then, because $dB_{G_{ij}} = d\mathcal{M}((g_i - \bar{g}_i)(g_j - \bar{g}_j) - G_{ij}) / \|(g_i - \bar{g}_i)(g_j - \bar{g}_j) - G_{ij}\|$, we can combine the above results to see

$$dB_{G_{ij}}\, dB_{G_{kl}} = \frac{G_{ik}G_{jl} + G_{il}G_{jk}}{\sqrt{G_{ii}G_{jj} + G_{ij}^2}\sqrt{G_{kk}G_{ll} + G_{kl}^2}} dt, \tag{123}$$

which is equation (24) of the main text.

. . . . . . . . . . . . . . . . . . . . . . . . . . . . . . . . . . . . . . . . . . . . . . . . . . . . . . . .

**Vectorizing and Sphering Matrix-Valued Martingales**

Now consider the $d \times d$ symmetric matrix of continuous martingales $\mathbf{\Omega_G}$ defined so that its $ij$-th entry is given by $(\mathbf{\Omega_G})_{ij} = \mathcal{M}((g_i - \bar{g}_i)(g_j - \bar{g}_j) - G_{ij})$ (that is $\mathbf{\Omega_G}$ is a matrix-valued martingale). Using this definition, we can intuitively think of $(d\mathbf{\Omega_G})_{ij} = d\mathcal{M}((g_i - \bar{g}_i)(g_j - \bar{g}_j) - G_{ij})$ as a normally distributed random variable with mean zero and variance given by $(d\mathbf{\Omega_G})^2_{ij} = d\mathcal{M}((g_i - \bar{g}_i)(g_j - \bar{g}_j) - G_{ij})^2 = v\, n\,(G_{ii}G_{jj} + G^2_{ij})dt$. We can extend this perspective to consider $d\mathbf{\Omega_G}$ as a matrix-valued normal random variable, in which case the covariance structure is summarized by a fourth-order tensor. In order to leverage a sphering argument to express $d\mathbf{\Omega_G}$ in terms of a matrix-valued Brownian motion differential $d\mathbf{B_G}$, I *vectorize* the matrix $d\mathbf{\Omega_G}$ by putting into one-to-one correspondence with the pair of indices $(i,j)$ for $1 \le i \le j \le d$ with a vectorized index $\alpha$ (the exact mapping between $\alpha$ and $(i,j)$, denoted $\alpha \leftrightarrow (i,j)$, doesn't matter so long as it's one-to-one). In particular, the vectorized index $\alpha$ spans $u = d(d+1)/2$ different values. With the vectorized index $\alpha$, write $\mathrm{vec}[d\mathbf{\Omega_G}]$ for the $u$-dimensional vector with $\alpha$-th entry given by $\mathrm{vec}[d\mathbf{\Omega_G}]_\alpha = (d\mathbf{\Omega_G})_{ij}$. We can then intuitively think of $\mathrm{vec}[d\mathbf{\Omega_G}]$ as a multivariate normal vector with mean zero and covariance matrix given by the outer-product $\mathrm{mat}[\mathbf{C}] := \mathrm{vec}[d\mathbf{\Omega_G}]\mathrm{vec}[d\mathbf{\Omega_G}]^\top$. More precisely, given another vectorized index $\beta \leftrightarrow (k,l)$, the matrix $\mathrm{mat}[\mathbf{C}]$ has as its $\alpha\beta$-th entry

$$\mathrm{mat}[\mathbf{C}]_{\alpha\beta} = (d\mathbf{\Omega_G})_{ij}(d\mathbf{\Omega_G})_{kl} = v\, n\,(G_{ik}G_{jl} + G_{il}G_{jk})dt. \tag{124}$$

Because the informal argument made here leverages heuristics where products of stochastic differentials are deterministic quantities (e.g., $dB^2_{G_{ij}} = dt$), expectations are not needed to express covariance matrices such as $\mathrm{mat}[\mathbf{C}]$ (which otherwise would be needed in more formal settings). To simplify notation, these expectations are omitted for the rest of this argument.

Define the fourth-order tensor $\mathbf{\Gamma}$ such that $\Gamma_{ijkl} = G_{ik}G_{jl} + G_{il}G_{jk}$. Then $\mathbf{\Gamma} = \mathbf{G}\overline{\otimes}\mathbf{G} + \mathbf{G}\underline{\otimes}\mathbf{G}$, where $(\mathbf{G}\overline{\otimes}\mathbf{G})_{ijkl} = G_{ik}\, G_{jl}$ is called the *upper open product*, and $(\mathbf{G}\underline{\otimes}\mathbf{G})_{ijkl} = G_{il}\, G_{jk}$ is called the *lower open product*. Write $\mathrm{mat}[\mathbf{\Gamma}]$ as the $u \times u$ matrix ($u = d(d+1)/2$) with $\alpha\beta$-th entry ($\alpha \leftrightarrow (i,j)$ and $\beta \leftrightarrow (k,l)$) given by $\mathrm{mat}[\mathbf{\Gamma}]_{\alpha\beta} = \Gamma_{ijkl}$. We can then write the covariance matrix for $\mathrm{vec}[d\mathbf{\Omega_G}]$ as

$$\mathrm{mat}[\mathbf{C}] = v\, n\, \mathrm{mat}[\mathbf{\Gamma}]\, dt \tag{125}$$

Taking a similar sphering approach used to express $d\bar{\mathbf{z}}$ in terms of a vector of independent standard Brownian motions $\mathbf{B}_{\bar{\mathbf{z}}}$, we can write

$$\mathrm{vec}[d\mathbf{\Omega_G}] = \sqrt{v\, n}\sqrt{\mathrm{mat}[\mathbf{\Gamma}]}\mathrm{vec}[d\mathbf{B_G}], \tag{126}$$

where $\sqrt{\mathrm{mat}[\mathbf{\Gamma}]}$ is any $u \times u$ matrix that satisfies $\sqrt{\mathrm{mat}[\mathbf{\Gamma}]}\sqrt{\mathrm{mat}[\mathbf{\Gamma}]}^\top = \mathrm{mat}[\mathbf{\Gamma}]$, and $\mathrm{vec}[d\mathbf{B_G}]$ is multivariate normal with mean zero and covariance matrix equal to the $u \times u$ identity matrix times $dt$ (i.e., $\mathrm{vec}[d\mathbf{B_G}]$ has uncorrelated entries each with variance $dt$):

$$\mathrm{vec}[d\mathbf{B_G}]\mathrm{vec}[d\mathbf{B_G}]^\top = \mathbf{I}_u\, dt. \tag{127}$$

The vectorized indices $\alpha \leftrightarrow (i,j)$ and $\beta \leftrightarrow (k,l)$ tell us how to reshape the matrix $\mathrm{vec}[d\mathbf{B_G}]\mathrm{vec}[d\mathbf{B_G}]^\top$ into a fourth-order tensor:

$$(\mathrm{vec}[d\mathbf{B_G}]\mathrm{vec}[d\mathbf{B_G}]^\top)_{\alpha\beta} = (d\mathbf{B_G})_{ij}(d\mathbf{B_G})_{kl}. \tag{128}$$

The constraint asserted by equation (127) creates a problem that has a canonical solution given by $(d\mathbf{B_G})_{ij}(d\mathbf{B_G})_{kl} = \frac{1}{2}(\delta_{ik}\delta_{jl} + \delta_{il}\delta_{jk})\, dt$. In turn, this solution defines the fourth-order tensor $d\mathbf{B_G} \otimes d\mathbf{B_G}$ (in particular, $(d\mathbf{B_G} \otimes d\mathbf{B_G})_{ijkl} = (d\mathbf{B_G})_{ij}(d\mathbf{B_G})_{kl}$) that determines the covariance structure of the symmetric normal matrix $d\mathbf{B_G}$.

Again using the vectorized indices $\alpha \leftrightarrow (i,j)$ and $\beta \leftrightarrow (k,l)$, define $\sqrt{\mathbf{\Gamma}}$ to be the fourth-order tensor with $ijkl$-th entry given by $\sqrt{\mathbf{\Gamma}}_{ijkl} = \sqrt{\mathrm{mat}[\mathbf{\Gamma}]}_{\alpha\beta}$. Then, because

$$\mathrm{vec}[d\mathbf{\Omega_G}]_\alpha = \sqrt{v\, n}\sum_\beta \sqrt{\mathrm{mat}[\mathbf{\Gamma}]}_{\alpha\beta}\mathrm{vec}[d\mathbf{B_G}]_\beta, \tag{129}$$

we also have

$$(d\mathbf{\Omega_G})_{ij} = \sqrt{v\, n}\sum_{kl}\sqrt{\mathbf{\Gamma}}_{ijkl}(d\mathbf{B_G})_{kl}. \tag{130}$$

Define the double contraction $\sqrt{\mathbf{\Gamma}} : d\mathbf{B_G}$ as the $d \times d$ matrix with $ij$-th entry $(\sqrt{\mathbf{\Gamma}} : d\mathbf{B_G})_{ij} = \sum_{kl}\sqrt{\mathbf{\Gamma}}_{ijkl}(d\mathbf{B_G})_{kl}$. Then

$$d\mathbf{\Omega_G} = \sqrt{v\, n}\sqrt{\mathbf{\Gamma}} : d\mathbf{B_G}. \tag{131}$$

By definition of $\sqrt{\mathrm{mat}[\mathbf{\Gamma}]}$, we have $\mathrm{mat}[\mathbf{\Gamma}]_\alpha = \sum_\gamma \sqrt{\mathrm{mat}[\mathbf{\Gamma}]}_{\alpha\gamma}\sqrt{\mathrm{mat}[\mathbf{\Gamma}]}_{\beta\gamma}$, with the sum taken across another vectorized index $\gamma \leftrightarrow (m,n)$. This is equivalent to the statement $\Gamma_{ijkl} = \sum_{mn}\sqrt{\mathbf{\Gamma}}_{ijmn}\sqrt{\mathbf{\Gamma}}_{klmn}$, with the sum on the right used to define the double contraction $\sqrt{\mathbf{\Gamma}} : \sqrt{\mathbf{\Gamma}}$. In particular, we have $\mathbf{\Gamma} = \sqrt{\mathbf{\Gamma}} : \sqrt{\mathbf{\Gamma}}$. A choice of $\sqrt{\mathbf{\Gamma}}$ that satisfies $\mathbf{\Gamma} = \mathbf{G}\overline{\otimes}\mathbf{G} + \mathbf{G}\underline{\otimes}\mathbf{G}$ is $\sqrt{\mathbf{\Gamma}}_{ijkl} = (\sqrt{\mathbf{G}}_{ik}\sqrt{\mathbf{G}}_{jl} + \sqrt{\mathbf{G}}_{il}\sqrt{\mathbf{G}}_{jk})/\sqrt{2}$. This notation then justifies equation (21) in the main text (the matrix-valued stochastic differential equation for $\mathbf{G}$ under the Brownian Motion Gradient version of the framework).

. . . . . . . . . . . . . . . . . . . . . . . . . . . . . . . . . . . . . . . . . . . . . . . . . . . . . . . . . . . .

###### 5.4.3. *Uncorrelated noise processes driving* $n, \bar{\mathbf{z}}, \mathbf{G}$

Finally, to show that the noise processes driving the stochastic dynamics of additive genetic variances and abundance do not covary, use the multiplicative property to compute $dn\, dG_{ij} = d\mathcal{M}(1)d\mathcal{M}((g_i - \bar{g}_i)(g_j - \bar{g}_j) - G_{ij}) = \langle\, 1,\, (g_i - \bar{g}_i)(g_j - \bar{g}_j) - G_{ij}\,\rangle\, dt = v\, n \int_{\mathbb{R}^d} ((g_i - \bar{g}_i)(g_j - \bar{g}_j) - G_{ij})\tilde{p}(\mathbf{g})d\mathbf{g}\, dt = 0\, dt = 0$ for any distribution of additive genetic values $\mathbf{g}$. Following the same intuitive perspective used to justify the sphering argument above, we see that the noise processes driving $G_{ij}$ and $n$ are uncorrelated. For additive genetic variances and mean traits, compute

$$dG_{ij}\, d\bar{z}_k = \langle\, (g_i - \bar{g}_i)(g_j - \bar{g}_j) - G_{ij},\, g_k - \bar{g}_k\,\rangle\, dt = v\, n \int_{\mathbb{R}^d} (g_i - \bar{g}_i)(g_j - \bar{g}_j)(g_k - \bar{g}_k)\tilde{p}(\mathbf{g})d\mathbf{g}\, dt = n\, v\, S_{ijk}, \tag{132}$$

where $S_{ijk}$ is the $ijk$-th entry of the third-order skew tensor for the distribution of additive genetic values $\tilde{p}(\mathbf{g})$. For multivariate normal distributions $S_{ijk} = 0$. Hence, the noise processes driving the stochastic dynamics of additive genetic variances and mean traits do not covary when additive genetic values are multivariate normal.

### 6. Derivations of the dynamics of genetic correlation and of the inverse of its hyperbolic tangent in response to drift

#### 6.1. Derivation of the Stochastic Differential Equation $d\rho = -\frac{1}{2}\frac{v}{n}\rho(1-\rho^2)dt + \sqrt{\frac{v}{n}}(1-\rho^2)dB_\rho$

The genetic correlation between traits $i$ and $j$ is given by $\rho_{ij} = G_{ij}/\sqrt{G_{ii}G_{jj}}$. The goal here is to derive a stochastic differential equation that characterizes the response of $\rho_{ij}$ to drift for asexually reproducing populations (without mutation, selection, or any other process affecting $\rho_{ij}$). Itô's formula applied to $\rho_{ij} = \rho(G_{ij}, G_{ii}, G_{jj}) := G_{ij}/\sqrt{G_{ii}G_{jj}}$ provides

$$\begin{aligned} d\rho = (\partial_{ij}\rho)dG_{ij} + (\partial_{ii}\rho)dG_{ii} + (\partial_{jj}\rho)dG_{jj} \\ + \frac{1}{2}\left[(\partial^2_{ij}\rho)(dG_{ij})^2 + (\partial^2_{ii}\rho)(dG_{ii})^2 + (\partial^2_{jj}\rho)(dG_{jj})^2\right] \\ + (\partial_{ij}\partial_{ii}\rho)dG_{ij}dG_{ii} + (\partial_{ij}\partial_{jj}\rho)dG_{ij}dG_{jj} + (\partial_{ii}\partial_{jj}\rho)dG_{ii}dG_{jj}. \end{aligned} \tag{133}$$

Throughout this section $\rho$ is shorthand for $\rho(G_{ij}, G_{ii}, G_{jj})$. The partial derivatives of $\rho$ are

$$\partial_{ij}\rho = \frac{1}{\sqrt{G_{ii}G_{jj}}},\ \partial_{ii}\rho = -\frac{1}{2}\frac{G_{ij}}{G_{ii}^{3/2}G_{jj}^{1/2}} = -\frac{\rho}{2G_{ii}},\ \partial_{jj}\rho = -\frac{1}{2}\frac{G_{ij}}{G_{ii}^{1/2}G_{jj}^{3/2}} = -\frac{\rho}{2G_{jj}}, \tag{134}$$

and the second derivatives are

$$\begin{aligned} \partial^2_{ij}\rho = 0,\ \partial_{ij}\partial_{ii}\rho = -\frac{1}{2}\frac{1}{G_{ii}^{3/2}G_{jj}^{1/2}} = -\frac{1}{2},\frac{\partial_{ij}\rho}{G_{ii}},\ \partial_{ij}\partial_{jj}\rho = -\frac{1}{2}\frac{1}{G_{ii}^{1/2}G_{jj}^{3/2}} = -\frac{1}{2},\frac{\partial_{ij}\rho}{G_{jj}}, \\ \partial^2_{ii}\rho = \frac{3}{4}\frac{G_{ij}}{G_{ii}^{5/2}G_{jj}^{1/2}} = \frac{3}{4}\frac{\rho}{G_{ii}^2},\ \partial^2_{jj}\rho = \frac{3}{4}\frac{G_{ij}}{G_{ii}^{1/2}G_{jj}^{5/2}} = \frac{3}{4}\frac{\rho}{G_{jj}^2},\ \partial_{ii}\partial_{jj}\rho = \frac{1}{4}\frac{G_{ij}}{G_{ii}^{3/2}G_{jj}^{3/2}} = \frac{1}{4}\frac{\rho}{G_{ii}G_{jj}}. \end{aligned} \tag{135}$$

Write $d\rho = K + L$, with $K$ containing the first derivatives and $L$ containing the second derivatives. In particular

$$K = (\partial_{ij}\rho)dG_{ij} + (\partial_{ii}\rho)dG_{ii} + (\partial_{jj}\rho)dG_{jj}. \tag{136}$$

We can split $K$ into deterministic and stochastic components $K = K_{\text{det}} + K_{\text{stoch}}$ such that

$$K_{\text{det}} = (\partial_{ij}\rho)dG_{ij}^{(\text{det})} + (\partial_{ii}\rho)dG_{ii}^{(\text{det})} + (\partial_{jj}\rho)dG_{jj}^{(\text{det})}, \tag{137}$$

with $dG_{ij}^{(\text{det})} = -\frac{v}{n}G_{ij}\, dt$ and

$$K_{\text{stoch}} = (\partial_{ij}\rho)dG_{ij}^{(\text{stoch})} + (\partial_{ii}\rho)dG_{ii}^{(\text{stoch})} + (\partial_{jj}\rho)dG_{jj}^{(\text{stoch})}, \tag{138}$$

with $dG_{ij}^{(\text{stoch})} = \frac{1}{n}d\mathcal{M}((g_i - \bar{g}_i)(g_j - \bar{g}_j) - G_{ij})$. Now plugging in expressions for the partial derivatives of $\rho$ into $K_{\text{det}}$ and canceling terms that appear in both numerator and denominator leads to

$$K_{\text{det}} = -\frac{v}{n}\left(\frac{G_{ij}}{\sqrt{G_{ii}G_{jj}}} - \frac{1}{2}\frac{G_{ij}}{\sqrt{G_{ii}G_{jj}}} - \frac{1}{2}\frac{G_{ij}}{\sqrt{G_{ii}G_{jj}}}\right) = 0. \tag{139}$$

. . . . . . . . . . . . . . . . . . . . . . . . . . . . . . . . . . . . . . . . . . . . . . . . . . . . . . . . . . . .

##### 6.1.1. $K_{\text{stoch}}$

For $K_{\text{stoch}}$, substitute $dG_{ij}^{(\text{stoch})} = \frac{1}{n} d\mathcal{M}((g_i - \bar{g}_i)(g_j - \bar{g}_j) - G_{ij})$ to get

$$K_{\text{stoch}} = \frac{1}{n}(\partial_{ij}\rho) d\mathcal{M}[(g_i - \bar{g}_i)(g_j - \bar{g}_j) - G_{ij}] + \frac{1}{n}(\partial_{ii}\rho) d\mathcal{M}[(g_i - \bar{g}_i)^2 - G_{ii}] + \frac{1}{n}(\partial_{jj}\rho) d\mathcal{M}[(g_j - \bar{g}_j)^2 - G_{jj}]. \tag{140}$$

The additive property then justifies

$$K_{\text{stoch}} = \frac{1}{n} d\mathcal{M}[(\partial_{ij}\rho)((g_i - \bar{g}_i)(g_j - \bar{g}_j) - G_{ij}) + (\partial_{ii}\rho)((g_i - \bar{g}_i)^2 - G_{ii}) + (\partial_{jj}\rho)((g_j - \bar{g}_j)^2 - G_{jj})]. \tag{141}$$

Define

$$H_{ij} := (\partial_{ij}\rho)\Big((g_i - \bar{g}_i)(g_j - \bar{g}_j) - G_{ij}\Big) + (\partial_{ii}\rho)\Big((g_i - \bar{g}_i)^2 - G_{ii}\Big) + (\partial_{jj}\rho)\Big((g_j - \bar{g}_j)^2 - G_{jj}\Big), \tag{142}$$

so that $K_{\text{stoch}} = \frac{1}{n} d\mathcal{M}(H_{ij})$. Distributing and then plugging the expressions for the partial derivatives of $\rho$, and then substituting $\rho = G_{ij}/\sqrt{G_{ii}G_{jj}}$ and canceling terms that appear in both numerator and denominator leads to

$$H_{ij} = \left(\frac{(g_i - \bar{g}_i)(g_j - \bar{g}_j)}{\sqrt{G_{ii}G_{jj}}} - \rho\right) + \left(-\frac{\rho}{2}\frac{(g_i - \bar{g}_i)^2}{G_{ii}} + \frac{\rho}{2}\right) + \left(-\frac{\rho}{2}\frac{(g_j - \bar{g}_j)^2}{G_{jj}} + \frac{\rho}{2}\right) = \frac{(g_i - \bar{g}_i)(g_j - \bar{g}_j)}{\sqrt{G_{ii}G_{jj}}} - \frac{\rho}{2}\frac{(g_i - \bar{g}_i)^2}{G_{ii}} - \frac{\rho}{2}\frac{(g_j - \bar{g}_j)^2}{G_{jj}} \tag{143}$$

The scaling property states $d\mathcal{M}(H_{ij}) = \|H_{ij}\| dB_{\rho_{ij}}$ for some standard Brownian motion $B_{\rho_{ij}}$. Additionally, we have $\|H_{ij}\| = \sqrt{v\,n \int_{\mathbb{R}^d} (H_{ij})^2 \tilde{p}(\mathbf{g}) d\mathbf{g}} = \sqrt{v\,n}\,\overline{H_{ij}^2}$. Expanding the square leads to

$$\overline{H_{ij}^2} = \frac{\overline{(g_i - \bar{g}_i)^2(g_j - \bar{g}_j)^2}}{G_{ii}G_{jj}} + \frac{\rho^2}{4}\frac{\overline{(g_i - \bar{g}_i)^4}}{G_{ii}^2} + \frac{\rho^2}{4}\frac{\overline{(g_j - \bar{g}_j)^4}}{G_{jj}^2} - \rho\frac{\overline{(g_i - \bar{g}_i)^3(g_j - \bar{g}_j)}}{G_{ii}^{3/2}G_{jj}^{1/2}} - \rho\frac{\overline{(g_i - \bar{g}_i)(g_j - \bar{g}_j)^3}}{G_{ii}^{1/2}G_{jj}^{3/2}} + \frac{\rho^2}{2}\frac{\overline{(g_i - \bar{g}_i)^2(g_j - \bar{g}_j)^2}}{G_{ii}G_{jj}}. \tag{144}$$

Isserlis' theorem (Isserlis, 1918) immediately implies

$$\overline{(g_i - \bar{g}_i)(g_j - \bar{g}_j)(g_k - \bar{g}_k)(g_l - \bar{g}_l)} = G_{ij}G_{kl} + G_{ik}G_{jl} + G_{il}G_{jk}. \tag{145}$$

Hence

$$\overline{(g_i - \bar{g}_i)^2(g_j - \bar{g}_j)^2} = G_{ii}G_{jj} + 2G_{ij}^2, \qquad \overline{(g_i - \bar{g}_i)^3(g_j - \bar{g}_j)} = 3G_{ii}G_{ij}, \qquad \overline{(g_i - \bar{g}_i)(g_j - \bar{g}_j)^3} = 3G_{jj}G_{ij}. \tag{146}$$

Substituting into $\overline{H_{ij}^2}$ gives

$$\overline{H_{ij}^2} = \frac{G_{ii}G_{jj} + 2G_{ij}^2}{G_{ii}G_{jj}} + \frac{\rho^2}{4}\frac{3G_{ii}^2}{G_{ii}^2} + \frac{\rho^2}{4}\frac{3G_{jj}^2}{G_{jj}^2} - \rho\frac{3G_{ii}G_{ij}}{G_{ii}^{3/2}G_{jj}^{1/2}} - \rho\frac{3G_{jj}G_{ij}}{G_{ii}^{1/2}G_{jj}^{3/2}} + \frac{\rho^2}{2}\frac{G_{ii}G_{jj} + 2G_{ij}^2}{G_{ii}G_{jj}}. \tag{147}$$

Cancelling common terms appearing in numerators and denominators, and substituting $G_{ij}/\sqrt{G_{ii}G_{jj}} = \rho$ gives

$$\overline{H_{ij}^2} = 1 + 2\rho^2 + \frac{3}{4}\rho^2 + \frac{3}{4}\rho^2 - 3\rho^2 - 3\rho^2 + \frac{\rho^2}{2}(1 + 2\rho^2) = 1 - 2\rho^2 + \rho^4 = (1 - \rho^2)^2. \tag{148}$$

Hence, $\|H_{ij}\| = \sqrt{v\,n}\,(1 - \rho^2)$, and, equating $B_\rho = B_{\rho_{ij}}$, we then have

$$K = K_{\text{det}} + K_{\text{stoch}} = 0 + \frac{1}{n}\|H_{ij}\|\, dB_{\rho_{ij}} = \sqrt{\frac{v}{n}}\,(1 - \rho^2)\, dB_\rho. \tag{149}$$

- - - - - - - - - - - - - - - - - - - - - - - - - - - - - - - - - - - - - - - - - - - - - - - - - - - - - - - - - - - -

### 6.1.2. $L$

For $L$, use the multiplication property to find

$$(dG_{ij})(dG_{kl}) = d\mathcal{M}((g_i - \bar{g}_i)(g_j - \bar{g}_j) - G_{ij}), d\mathcal{M}((g_k - \bar{g}_k)(g_l - \bar{g}_l) - G_{kl}) = \frac{v}{n}(G_{ik}G_{jl} + G_{il}G_{jk})\, dt. \tag{150}$$

In particular,

$$(dG_{ij})^2 = \frac{v}{n}(G_{ii}G_{jj} + G_{ij}^2)\, dt, \qquad (dG_{ii})(dG_{jj}) = 2\frac{v}{n}G_{ij}^2\, dt. \tag{151}$$

From the expression for $d\rho$ above, we see $L$ is given by

$$L = \tfrac{1}{2}(\partial_{ii}^2\rho)(dG_{ii})^2 + \tfrac{1}{2}(\partial_{jj}^2\rho)(dG_{jj})^2 + (\partial_{ij}\partial_{ii}\rho)(dG_{ij})(dG_{ii}) + (\partial_{ij}\partial_{jj}\rho)(dG_{ij})(dG_{jj}) + (\partial_{ii}\partial_{jj}\rho)(dG_{ii})(dG_{jj}). \tag{152}$$

Plugging in results for second partial derivatives of $\rho$ then gives

$$\tfrac{1}{2}(\partial^2_{ii}\rho)(dG_{ii})^2 = \tfrac{1}{2}\left(\frac{3}{4}\frac{G_{ij}}{G_{ii}^{5/2}G_{jj}^{1/2}}\right)\left(\frac{2v}{n}G_{ii}^2\,dt\right) = \frac{v}{n}\frac{3}{4}\frac{G_{ij}}{G_{ii}^{1/2}G_{jj}^{1/2}}\,dt. \tag{153}$$

$$\tfrac{1}{2}(\partial^2_{jj}\rho)(dG_{jj})^2 = \tfrac{1}{2}\left(\frac{3}{4}\frac{G_{ij}}{G_{ii}^{1/2}G_{jj}^{5/2}}\right)\left(\frac{2v}{n}G_{jj}^2\,dt\right) = \frac{v}{n}\frac{3}{4}\frac{G_{ij}}{G_{ii}^{1/2}G_{jj}^{1/2}}\,dt. \tag{154}$$

$$(\partial_{ij}\partial_{ii}\rho)(dG_{ij})(dG_{ii}) = \left(-\frac{1}{2}\frac{1}{G_{ii}^{3/2}G_{jj}^{1/2}}\right)\left(\frac{2v}{n}G_{ii}G_{ij}\,dt\right) = -\frac{v}{n}\frac{G_{ij}}{G_{ii}^{1/2}G_{jj}^{1/2}}\,dt. \tag{155}$$

$$(\partial_{ij}\partial_{jj}\rho)(dG_{ij})(dG_{jj}) = \left(-\frac{1}{2}\frac{1}{G_{ii}^{1/2}G_{jj}^{3/2}}\right)\left(\frac{2v}{n}G_{jj}G_{ij}\,dt\right) = -\frac{v}{n}\frac{G_{ij}}{G_{ii}^{1/2}G_{jj}^{1/2}}\,dt. \tag{156}$$

$$(\partial_{ii}\partial_{jj}\rho)(dG_{ii})(dG_{jj}) = \left(\frac{1}{4}\frac{G_{ij}}{G_{ii}^{3/2}G_{jj}^{3/2}}\right)\left(\frac{2v}{n}G_{ij}^2\,dt\right) = \frac{v}{n}\frac{1}{2}\frac{G_{ij}^3}{G_{ii}^{3/2}G_{jj}^{3/2}}\,dt. \tag{157}$$

Plugging these results into $L$ and simplifying leads to

$$L = \frac{v}{n}\frac{3}{2}\frac{G_{ij}}{G_{ii}^{1/2}G_{jj}^{1/2}}\,dt - \frac{v}{n}2\frac{G_{ij}}{G_{ii}^{1/2}G_{jj}^{1/2}}\,dt + \frac{v}{n}\frac{1}{2}\frac{G_{ij}^3}{G_{ii}^{3/2}G_{jj}^{3/2}}\,dt = \frac{v}{n}\left(\frac{3}{2}\rho - 2\rho + \frac{1}{2}\rho^3\right)dt = -\frac{1}{2}\frac{v}{n}\rho(1-\rho^2)\,dt. \tag{158}$$

---

#### 6.2. Stationary distribution and boundary behavior

In the last subsection we obtained

$$d\rho = b(\rho)\,dt + \sigma(\rho)\,dB_\rho, \tag{159}$$

with $b(\rho) = -\frac{1}{2}\frac{v}{n}\rho(1-\rho^2)$ and $\sigma(\rho) = \sqrt{\frac{v}{n}}(1-\rho^2)$. Because this is a regular one-dimensional diffusion, a formal stationary distribution (if it exists) is given by $\pi(\rho) \propto \exp\left(\int^\rho 2b(u)/\sigma^2(u)\,du\right)/\sigma^2(\rho)$ (Etheridge, 2010; Karlin and Taylor, 1981). Here $\sigma^2(\rho) = \frac{v}{n}(1-\rho^2)^2$ and $2b(\rho)/\sigma^2(\rho) = -\rho/(1-\rho^2)$, so $\int^\rho -u/(1-u^2)\,du = \frac{1}{2}\log(1-\rho^2)+C$, implying $\exp(\int^\rho 2b/\sigma^2) \propto (1-\rho^2)^{1/2}$. Substituting back gives the stationary *shape* $\hat{q}(\rho) \propto (1-\rho^2)^{-3/2}$, which is non-integrable on $(-1,1)$. Thus no normalizable stationary distribution exists.

To classify the boundary behavior, compute the scale and speed densities. The scale density is $s'(\rho) = \exp(-\int^\rho 2b(u)/\sigma(u)^2\,du) \propto (1-\rho^2)^{-1/2}$, yielding the scale function $s(\rho) = \int_0^\rho (1-u^2)^{-1/2}du = \arcsin(\rho)$, which is finite at $\rho = \pm 1$. The speed density is $m(\rho) = 2/(\sigma(\rho)^2 s'(\rho)) = \frac{2n}{v}(1-\rho^2)^{-3/2}$, which diverges non-integrably at both boundaries. Referring to Table 6.2 for boundary classification in Chapter 15 of Karlin and Taylor (1981), this combination (finite scale function at the boundary together with divergent speed measure) implies that $\rho = \pm 1$ are attracting in the sense that probability mass is pushed toward them, but they are unattainable from the interior (i.e., the probability of reaching them in finite time zero) for all $\rho \in (-1,1)$.

---

#### 6.3. Derivation of the stochastic differential equation $du = \frac{1}{2}\frac{v}{n}\tanh(u)dt + \sqrt{\frac{v}{n}}dB_u$

In Supplementary Material, Section 6.1 we obtained $d\rho = -\frac{1}{2}\frac{v}{n}\rho(1-\rho^2)\,dt + \sqrt{\frac{v}{n}}(1-\rho^2)\,dB_\rho$. Define $u := \tanh^{-1}(\rho)$ (the inverse hyperbolic tangent of $\rho$). Itô's formula states

$$du = u'(\rho)\,d\rho + \frac{1}{2}u''(\rho)\,(d\rho)^2. \tag{160}$$

Use the identity

$$u = \frac{1}{2}\ln\left(\frac{1+\rho}{1-\rho}\right) \tag{161}$$

in combination with elementary calculus to obtain

$$u'(\rho) = \frac{d}{d\rho}\tanh^{-1}(\rho) = \frac{1}{1-\rho^2} \tag{162}$$

and

$$u''(\rho) = \frac{d}{d\rho}\left(\frac{1}{1-\rho^2}\right) = \frac{2\rho}{(1-\rho^2)^2}. \tag{163}$$

Set $a(\rho) = -\frac{1}{2}\frac{v}{n}\rho(1-\rho^2)$, and $b(\rho) = \sqrt{\frac{v}{n}}(1-\rho^2)$. Then $d\rho = a(\rho)\,dt + b(\rho)\,dB_\rho$. Because $B_\rho$ is a standard Brownian motion, we have the general heuristics $dt^2 = dt\,dB_\rho = 0$ and $(dB_\rho)^2 = dt$. Expanding the square $(d\rho)^2$ and applying these heuristics leads to

$$(d\rho)^2 = b^2(\rho)\,dt = \frac{v}{n}(1-\rho^2)^2\,dt. \tag{164}$$

Substituting into Itô's formula for $du$ leads to

$$du = u'(\rho)\,d\rho + \frac{1}{2}u''(\rho)\,(d\rho)^2 = u'(\rho)\big[a(\rho)\,dt + b(\rho)\,dB_\rho\big] + \frac{1}{2}u''(\rho)b^2(\rho)\,dt. \tag{165}$$

Collect $dt$ and $dB_\rho$ to get

$$du = \left[u'(\rho)a(\rho) + \frac{1}{2}u''(\rho)b^2(\rho)\right]dt + u'(\rho)b(\rho)\,dB_\rho. \tag{166}$$

Note

$$u'(\rho)b(\rho) = \frac{1}{1-\rho^2}\sqrt{\frac{v}{n}}(1-\rho^2) = \sqrt{\frac{v}{n}}, \tag{167}$$

and

$$u'(\rho)a(\rho) = \frac{1}{1-\rho^2}\left(-\frac{1}{2}\frac{v}{n}\rho(1-\rho^2)\right) = -\frac{1}{2}\frac{v}{n}\rho, \tag{168}$$

and also

$$\frac{1}{2}u''(\rho)b^2(\rho) = \frac{1}{2}\left(\frac{2\rho}{(1-\rho^2)^2}\right)\left(\frac{v}{n}(1-\rho^2)^2\right) = \frac{v}{n}\rho. \tag{169}$$

Then

$$\left[u'(\rho)a(\rho) + \frac{1}{2}u''(\rho)b^2(\rho)\right]dt = \frac{1}{2}\frac{v}{n}\rho\,dt. \tag{170}$$

Substituting $\rho = \tanh(u)$ and setting $B_u := B_\rho$, we then arrive at

$$du = \frac{1}{2}\frac{v}{n}\tanh(u)\,dt + \sqrt{\frac{v}{n}}\,dB_u. \tag{171}$$

. . . . . . . . . . . . . . . . . . . . . . . . . . . . . . . . . . . . . . . . . . . . . . . . . . . . . . . . . . .

##### 6.3.1. *Approximating* $du$ *when* $|u| \gg 1$

When $u \gg 1$, we can justify $du \approx v\,dt/2n + \sqrt{v/n}dB_u$ by leveraging the identity $\tanh(u) = (e^u - e^{-u})/(e^u + e^{-u})$. For $u \gg 1$, $e^u$ dominates $e^{-u}$. So divide the numerator and denominator by $e^u$ to get

$$\tanh(u) = \frac{1-e^{-2u}}{1+e^{-2u}}. \tag{172}$$

Set $\varepsilon := e^{-2u}$ so $u \gg 1$ implies $0 < \varepsilon \ll 1$, and $\tanh(u) = (1-\varepsilon)/(1+\varepsilon)$. To first order in $\varepsilon$ we have $1/(1+\varepsilon) \approx 1-\varepsilon$. So, again to first order in $\varepsilon$, we have

$$\tanh(u) \approx (1-\varepsilon)(1-\varepsilon) \approx 1 - 2\varepsilon. \tag{173}$$

Back substituting $\varepsilon = e^u$ provides $\tanh(u) \approx 1 - 2e^{-2u}$. Hence $\tanh(u) \approx 1$ when $u \gg 1$. An analogous argument holds for $u \ll -1$ which leads to the approximation $\tanh(u) \approx -1$. Taken together, when $u \gg 1$ we have

$$du \approx \frac{1}{2}\frac{v}{n}dt + \sqrt{\frac{v}{n}}\,dB_u, \tag{174}$$

and when $u \ll -1$ we have

$$du \approx -\frac{1}{2}\frac{v}{n}dt + \sqrt{\frac{v}{n}}\,dB_u. \tag{175}$$

. . . . . . . . . . . . . . . . . . . . . . . . . . . . . . . . . . . . . . . . . . . . . . . . . . . . . . . . . . .

##### 6.3.2. *Approximating* $\mathbb{E}[u_t]$ *when* $t \gg 1$

To understand the behavior of $\mathbb{E}[u_t]$ when $t \gg 1$, recall that $u_t = \tanh^{-1}(\rho_t)$ satisfies

$$du_t = \frac{1}{2}\frac{v}{n}\tanh(u_t)\,dt + \sqrt{\frac{v}{n}}\,dB_u, \tag{176}$$

so in integral form

$$u_t = u_0 + \frac{1}{2}\frac{v}{n}\int_0^t \tanh(u_s)\,ds + \sqrt{\frac{v}{n}}\,B_u. \tag{177}$$

Taking expectations and using $\mathbb{E}[B_u] = 0$ gives the exact identity

$$\mathbb{E}[u_t] = u_0 + \frac{1}{2}\frac{v}{n}\int_0^t \mathbb{E}[\tanh(u_s)]\,ds. \tag{178}$$

The ode $\dot{q} = a\tanh(q)$ has the solution $q_t = \sinh^{-1}\left(\sinh(q_0)\,e^{a\,t}\right)$. To check, set $S_t = \sinh(q_0)\,e^{a\,t} = \sinh(q_t)$ so that $q_t = \sinh^{-1}(S_t)$ and hence $\dot{q} = \dot{S}/\sqrt{1+S^2}$, $\dot{S}_t = a\,S_t = a\sinh(q_t)$, and $\sqrt{1+S_t^2} = \cosh(q_t)$ (because $\sinh^2(q) - \cosh^2(q) = 1$). Thus, $\dot{q} = a\sinh(q)/\cosh(q) = a\tanh(q)$. For positive $u$, $\tanh(u)$ is convex and Jensen's inequality (Jensen, 1906) then justifies $\tanh(\mathbb{E}[u]) \le \mathbb{E}[\tanh(u)]$. This implies that the solution to $\frac{d}{dt}\mathbb{E}[u_t] = a\,\mathbb{E}[\tanh(u_s)] \ge a\tanh(\mathbb{E}[u_s])$ will satisfy $\mathbb{E}[u_t] \ge q_t$ for all $t \ge 0$ given that $q_0 = u_0 > 0$ (here I'm assuming $u_0 = \mathbb{E}[u_0]$ is not random). In addition, because $|\tanh(u)| \le 1$ for all $u$, we also have $\frac{d}{dt}\mathbb{E}[u_t] = a\,\mathbb{E}[\tanh(u_t)] \le a$ which implies $q_t \le \mathbb{E}[u_t] \le u_0 + a\,t$ for all $t \ge 0$. Asymptotically, $q_t \approx q_0 + a\,t$ for large $t$ which then implies $\mathbb{E}[u_t] \approx u_0 + a\,t$ for large $t$. Because tanh is an odd function (i.e., $\tanh(-u) = -\tanh(u)$), the analogous result holds for $u_0 < 0$. Letting $a = v/2n$ gives the approximation of $\mathbb{E}[u_t]$ discussed below equation (40) in the main text.

### 7. Justification of Martingale Process $\mathcal{M}_t$ and Multivariate DAGA by a Diffusion-Limit

The martingale process $\mathcal{M}_t$ characterized by equation (64) above in Supplementary Material, Section 4.1 (which establishes the mathematical foundation for the framework presented in the main text) can be formally constructed using diffusion-limits of individual-based branching processes. To illustrate one way to do this, I describe in the first subsection below an individual-based process and an approach to rescale it that are mathematically convenient. This choice is meant to keep the illustration as accessible as possible. It should be noted, however, that many alternative approaches that converge to the same diffusion-limit exist (including ones based on continuous-time or discrete-time individual-based processes). Hence, more biologically appealing but mathematically involved approaches are possible.

General conditions under which these diffusion-limits converge are described by a theorem that was proved by Méléard and Roelly (1993). Hence, after describing the individual-based process and the approach to rescaling, I simply demonstrate that the conditions required by the theorem of Méléard and Roelly (1993) are satisfied. Finally, an important limitation of the proof given by Méléard and Roelly (1993) is the requirement for the fitness function $m(\nu, \mathbf{z})$ to be bounded in both $\nu$ and $\mathbf{z}$. Classical models of competition, cooperation, and directional and stabilizing selection rely on fitness functions that are unbounded either in $\nu$ (for competition and cooperation) or in $\mathbf{z}$ (for directional and stabilizing selection). Therefore, in the third subsection below I describe an approach to adapt the result of Méléard and Roelly (1993) to allow for such unbounded fitness functions.

Before describing a possible approach to construct the framework with unbounded $m$, in the second subsection I show how to obtain multivariate DAGA (i.e., equation (1) of the main text). This is done by observing that, in the absence of demographic stochasticity, equation (64) above characterizes weak solutions to multivariate DAGA.

#### 7.1. Informal Description of the Diffusion-Limit

Following the notation introduced in section 3 above, the state of the individual-based process at time $t \geq 0$ is summarized by the discrete population measure $N_t = \sum_{i=1}^{n(t)} \delta_{\mathbf{z}_i(t)}$, where $\delta_{\mathbf{z}_i(t)}$ is the point-mass representing the $i$-th individual in the population. Traits of individuals are determined at birth and are constant throughout their lifetime. Because individuals can be added or removed from the population, their indexing will depend on time. If individual $j$ is removed from the population at time $t$, the index of individual $j+k$ is replaced by $j+k-1$ for each $k = 1, \ldots, n(t) - j$ (i.e., the remaining individuals with indices greater than the individual being removed "move down by 1"). If a number $j$ of individuals is added to the population at time $t$, they are given the indices $n(t^-)+1, \ldots, n(t^-)+j$ in no particular order (where $n(t^-)$ is the size of the population an instant before time $t$ such that $n(t) = n(t^-)+j$ is the updated population size after the $j$ individuals are added). Even though the trait value of each individual is constant during its lifetime, the indexed values $\mathbf{z}_i(t)$ appearing in the subscripts of the point masses depend on time because the indexing is dynamic and different individuals will typically have different trait values.

Individual lifetimes are exponentially distributed with constant rate $\rho > 0$, and are drawn independently at birth. At the end of an individuals lifetime, it is removed and replaced by a Poisson distributed number of offspring. When this occurs, the individual being replaced is said to "branch". Biologically, these assumptions imply that reproduction is asexual and occurs simultaneously with death. Suppose individual $i$ branches at time $t$. The mean of the Poisson distributed number of offspring that replace it is written as $W(N_{t^-}, \mathbf{z}_i(t^-))$ (biologically, this is called the *fitness* of the branching individual because it quantifies the expected lifetime reproductive output of that individual). The shape of the fitness surface $W$ as a function of population state $N$ and individual trait value $\mathbf{z}$ determines how selection operates and how trends in population growth change over time. The individuals replacing the branching individual $i$ are assigned trait values that are independently drawn from a $d$-dimensional multivariate normal distribution with mean $\mathbf{z}_i$ and covariance matrix $\mathring{\mathbf{M}} = \mathbf{M}/\rho$. The variation of offspring trait values around their parental value establishes a model of mutation.

Rescaling this process begins with the same approach for rescaling individuals described in section 3. In particular, focusing on the initial state of the population $N_0 = \sum_{i=1}^{n(0)} \delta_{\mathbf{z}_i(0)}$, the $k$-the stage of rescaling is $N_0^k = \frac{1}{k}\sum_{i=1}^{kn(0)} \delta_{\mathbf{z}_i(0)}$ where at each stage traits are sampled from a $d$-dimensional multivariate normal distribution with mean vector $\bar{\mathbf{z}}_0$ and covariance matrix $\mathbf{P}_0$. The initial trait distribution can be much more general than this, but nothing is lost here by keeping it simple. This approach to rescaling individuals implies their "weight" is replaced by $\alpha_k = 1/k$ at stage $k$. Simultaneously, we replace the branching rate $\rho$ with $\rho_k = k\rho$. At branching, the mutation covariance matrix is rescaled as $\mathring{\mathbf{M}}_k = \mathbf{M}/\rho_k = \mathbf{M}/k$. Using this approach, we will find that $v = \rho$ in the diffusion-limit (which intuitively makes sense because faster branching rates imply greater demographic stochasticity). Finally, we replace the fitness surface $W$ with its $k$-th root $W^{1/k}$. I call the rescaled process $N_t^k$ a $k$-system. The theorem of Méléard and Roelly (1993) states that if $\alpha_k \to 0$ and $\rho_k \to \infty$ as $k \to \infty$ (which holds in this case) and, given population measures of the form $N^k = \alpha_k \sum_i \delta_i$ such that $N^k \to \mathcal{N}$ as $k \to \infty$, if the limit

$$m_k(N^k, \mathbf{z}) = (W^{1/k}(N^k, \mathbf{z}) - 1)\rho_k \tag{179}$$

converges to a function $m(\mathcal{N}, \mathbf{z})$ that is continuous and bounded in both $\mathcal{N}$ and $\mathbf{z}$, and finally if the product

$$v_k(N^k, \mathbf{z}) = (W^{2/k}(N^k, \mathbf{z}) - W^{1/k}(N^k, \mathbf{z}) + 1)\alpha_k \rho_k \tag{180}$$

converges a constant $v > 0$, then the rescaled sequence of individual-based processes described here converges to a solution of the martingale problem described in section 4 above. Note the last condition is satisfied in our case because as $k$ increases the roots of the fitness surface $W^{2/k}$ and $W^{1/k}$ become more and more similar so that for large $k$ we have the approximation $v_k \approx \alpha_k \rho_k$. But this just simplifies to the constant $v_k \approx \rho$. Because this approximation becomes exact as $k \to \infty$, we find $v = \rho$.

I provide an outline on how to extend this result to allow for an $m$ that is unbounded in both $\mathcal{N}$ and $\mathbf{z}$ below. However, because this involves adapting proofs on the convergence of rescaled measure valued processes, there are unavoidable technical details. Understanding how to justify the multivariate DAGA partial differential equation (equation (1) in the main text) is more interesting to a broader audience, and so I address this next.

---

#### 7.2. Justification of multivariate DAGA by Deterministic Approximation

The most direct and simple way to justify multivariate DAGA is to set $v = 0$. Note this has to be done after taking the diffusion-limit. Otherwise this would require setting $\rho = 0$ so that branching would never occur in the rescaled processes, and thus no dynamics would be possible. With $v = 0$ equation (64) reduces to the deterministic equation

$$\int_{\mathbb{R}^d} f(\mathbf{z})\mathcal{N}_t(d\mathbf{z}) - \int_{\mathbb{R}^d} f(\mathbf{z})\mathcal{N}_0(d\mathbf{z}) = \int_0^t \int_{\mathbb{R}^d} (m(\mathcal{N}_s,\mathbf{z}) + \tfrac{1}{2}\nabla^\top \mathbf{M}\nabla) f(\mathbf{z})\mathcal{N}_s(d\mathbf{z})\, ds. \tag{181}$$

Solutions to this equation can be interpreted as so-called *generalized* or *weak solutions* to multivariate DAGA (an introduction to the topic is given by Evans, 2022), and this therefore provides a formal approach to derive equation (1) of the main text.

Instead of setting $v = 0$, it is also possible to obtain multivariate DAGA by considering very large population size. The main idea here is that the magnitude of demographic stochasticity is determined by the square-root of the quadratic variation described in equation (66), which itself is linear in the population measure. That is, the magnitude of demographic stochasticity can be roughly quantified by the square root of the population measure. In contrast, the magnitude of the deterministic dynamics are linear with respect to the population measure. Hence, when the population size is very large the effects of stochasticity will be negligible relative to the effects of the deterministic dynamics. Using this reasoning, one arrives again at the deterministic equation (181) above. Formalizing this approach then provides an alternative derivation of equation (1) in the main text.

---

#### 7.3. An Approach to Formally Construct the Framework with Unbounded $m(\nu, \mathbf{z})$

A common approach to formalize the diffusion-limit described above is to start by considering the set of finite measures $\mathfrak{M}_F := \{\mu : \mu(1) < \infty\}$, where $\mu(x) = \int_{\mathbb{R}^d} x(\mathbf{z})\mu(d\mathbf{z})$ so that $\mu(1)$ is the total mass (or abundance) of a population characterized by $\mu$ (Ethier and Kurtz, 1986). With $\mathfrak{M}_F$ in hand, now consider the set of random functions $\mathcal{N}$ mapping time $[0,\infty)$ into $\mathfrak{M}_F$ (so $\mathcal{N}$ is a $\mathfrak{M}_F$-valued stochastic process) that are 1) *right-continuous* and 2) are *left-limited.* Writing $\kappa(\mathcal{N}, \mathring{\mathcal{N}})$ as a (specific) notion of distance between $\mathcal{N}, \mathring{\mathcal{N}} \in \mathfrak{M}_F$ (see Ethier and Kurtz, 1986), right-continuous means that $\lim_{\varepsilon\to 0} \kappa(\mathcal{N}_{t+\varepsilon}, \mathcal{N}_t) = 0$, and left-limited means that there exists $\mathring{\mathcal{N}} \in \mathfrak{M}_F$ such that $\lim_{\varepsilon\to 0} \kappa(\mathcal{N}_{t-\varepsilon}, \mathring{\mathcal{N}}) = 0$. Denote this set of left-limited and right-continuous (llrc) functions by $D_{\mathfrak{M}_F}[0,\infty)$. Note that, by construction, *every* sample path of the individual-based process $N_t$ defined above is a member of $D_{\mathfrak{M}_F}[0,\infty)$. I denote this membership by $N_t \in D_{\mathfrak{M}_F}[0,\infty)$. In addition, we also have inclusion of the $k$-system $N_t^k \in D_{\mathfrak{M}_F}[0,\infty)$. Then construction of the diffusion-limit is accomplished by demonstrating the limit $\mathcal{N} = \lim_k N^k$ converges in $D_{\mathfrak{M}_F}[0,\infty)$ in a very specific way (see section 7.3.6 below). In addition, it must be shown that the limit point $\mathcal{N}$ solves a martingale problem (described in section 7.3.3 below).

Already this shows a restriction that the population measure must remain finite for all time, which prevents accommodating explosion. This issue may be overcome by developing a localized (in time) alternative where $\lim_k N_t^k$ converges in $D_{\mathfrak{M}_F}[0,\tau)$ for some (random) time $\tau$, where at times $t \geq \tau$ the limiting population process would be considered to have infinite abundance. Convergence in this case is formalized using localized martingale problems based on $\tau$. My short description in this paragraph prioritizes intuition, and is slightly misleading. A more accurate description of localization is given in the next section, and is discussed in section 4.6 of Ethier and Kurtz, (1986). An introduction to accommodating explosion with localization in the context of finite dimensional markov processes is given in Chapter 10 of Stroock and Varadhan (1997).

Another issue is that $\mathfrak{M}_F$ only requires measures to be finite (i.e., $\mu(\mathbb{R}^d) < \infty$). However, it is possible for phenotypic moments associated with finite measures to be infinite such that $\int_{\mathbb{R}^d} |\mathbf{z}|^p \mu(d\mathbf{z}) = \infty$ for some $p = 1, 2, \ldots$ (where $|\mathbf{z}|^p$ is the Euclidean norm of $\mathbf{z}$ raised to the $p$-th power). The framework presented here tracks first and second moments (so for $p = 1, 2$). Hence, another kind of explosion may occur here where phenotypic moments become infinite in finite time. Temporal localization may be used again to account for moment explosion. In particular, consider the restricted set of measures $\mathfrak{M}^p := \{\mu : \mu(w_p) < \infty\}$ where $w_p(\mathbf{z}) = 1 + |\mathbf{z}|^p$ and $\mu(w_p) = \int_{\mathbb{R}^d} w_p(\mathbf{z})\mu(d\mathbf{z})$. This is the set of finite measures with finite moments up to order $p$ (with strict inclusion $\mathfrak{M}^p \subset \mathfrak{M}_F$). This set of finite $p$-moment measures $\mathfrak{M}^p$ is, in a sense, dual to the set of so-called $p$-tempered measures $\mathfrak{M}_p$ introduced by Iscoe (1986). More precisely, for $\mu \in \mathfrak{M}^p$ we have $\mu(x) < \infty$ for functions $x(\mathbf{z})$ that grow at most like $|\mathbf{z}|^p$ for large $|\mathbf{z}|$, and for $\mathring{\mu} \in \mathfrak{M}_p$ we have $\mathring{\mu}(\mathring{x}) < \infty$ for functions $\mathring{x}(\mathbf{z})$ that decay at least like $1/|\mathbf{z}|^p$ for large $|\mathbf{z}|$. Here we are interested in $\mathfrak{M}^4$ (i.e., $\mathfrak{M}^p$ with $p = 4$) because, as shown by equation (8c) in the main text and in Supplementary Material, Section 5.4.2, fourth-order moments appear when characterizing the dynamics of the trait covariance matrix. Formal construction of the framework presented here may be done using localized martingales to characterize the convergence of $\lim_k N_t^k$ in $D_{\mathfrak{M}^4}[0,\tau)$ for some (random) time $\tau$ (this is only slightly misleading, and more accurate details are given in the next section).

Constructing the framework using a localized martingale problem with solutions taking values in $\mathfrak{M}^4$ involves a lot of technical details. Here I point out just a few of these details to outline how this approach might be formalized.

. . . . . . . . . . . . . . . . . . . . . . . . . . . . . . . . . . . . . . . . . . . . . . . . . . . . . . . . . . . . .

##### *7.3.1. The Killed Process*

To handle explosion (of either abundance or phenotypic moments), denote by $\partial$ a so-called *cemetery point* for $\mathfrak{M}^4$. The notation is sensible because $\partial A$ is often used to denote the boundary of a set $A$, and the cemetery point $\partial$ will act essentially as the boundary of $\mathfrak{M}^4$ in $\mathfrak{M}_F$ in the sense that when a process *leaves* $\mathfrak{M}^4$ it *hits* $\{\partial\}$. However, denoting the actual boundary of $\mathfrak{M}^4$ in $\mathfrak{M}_F$ by $\partial\mathfrak{M}^4$, for any $\mu \in \partial\mathfrak{M}^4$ we have that any neighborhood of $\mu$ intersects with $\mathfrak{M}^4$. In this sense, the boundary $\partial\mathfrak{M}^4$ cannot be separated from $\mathfrak{M}^4$. This is a general fact for the boundary of any set (Munkres, 2005). In contrast, the cemetery point $\partial$ is defined to have the feature of being isolated from $\mathfrak{M}^4$ (i.e., there exists a neighborhood of $\partial$ that does not intersect with $\mathfrak{M}^4$). This therefore distinguishes $\partial$ from $\partial\mathfrak{M}^4$. Although it may at first seem artificial, the cemetery point $\partial$ is helpful because it turns out that $\partial\mathfrak{M}^4 = \mathfrak{M}_F$ which prevents the definition of explosion in terms of a process *hitting* $\partial\mathfrak{M}^4$. When a process *hits* the set $\{\partial\}$ (which will be defined shortly), we *kill* the process at that time (also defined shortly). This justifies naming $\partial$ a cemetery point.

With $\partial$ in hand, we now work with the extended set $\bar{\mathfrak{M}}^4 := \mathfrak{M}^4 \cup \{\partial\}$. For a llrc $\bar{\mathfrak{M}}^4$-valued process $\mathcal{N}$ (in the notation above, $\mathcal{N} \in D_{\bar{\mathfrak{M}}^4}[0,\infty)$) and for $u = 1,2,\ldots$, define stopping times

$$\tau_u := \inf\{t \geq 0 : \mathcal{N}_t(w_4) \geq u\}, \quad \tau := \lim_{u\to\infty} \tau_u. \tag{182}$$

Then the process $\mathcal{N}$ hits $\{\partial\}$ at time $\tau$, at which point we kill it by setting $\mathcal{N}_t = \partial$ for $t \geq \tau$. It is possible that $\tau = \infty$, and in this case we let $\mathcal{N}$ live.

- - - - - - - - - - - - - - - - - - - - - - - - - - - - - - - - - - - - - - - - - - - - - - - - - - - - - - - - - - - -

##### 7.3.2. Brownian Motion Mutation Model

For the sake of convenience, replace the model of mutation in the individual-based model in section 7.1 (where offspring traits are drawn at branching and then kept constant throughout their lifetime) with a model where offspring traits are initially equal to their parental trait at branching and subsequently follow $d$-dimensional Brownian motion with covariance $\mathbf{M}$. The Brownian motion model of mutation is assumed to be independent from lifetime lengths so that the $i$-th offspring, when it branches, will have expectation equal to the trait of its parent (conditioned on knowing that parents' trait value) and covariance matrix $\mathbf{M}/\rho$ (computed using law of total covariance). This demonstrates agreement with the mutation-at-branching model used in section 7.1 above. The $d$-dimensional Brownian motion mutation model has an associated Kolmogorov forward equation (called the Fokker-Planck equation by physicists) that describes the evolution of the probability density $\zeta_t(\mathbf{z})$ of $\mathbf{z}_i(t)$ (conditioned on knowing the parental trait value), and is given by $\dot{\zeta} = \mathcal{A}\zeta$ where $\mathcal{A}x(\mathbf{z}) = \frac{1}{2}\nabla^\top\mathbf{M}\nabla x(\mathbf{z})$. The operator $\mathcal{A}$ is called the *infinitesimal generator* for the Brownian motion mutation model.

- - - - - - - - - - - - - - - - - - - - - - - - - - - - - - - - - - - - - - - - - - - - - - - - - - - - - - - - - - - -

##### 7.3.3. Localized Martingale Problem

Denoting $C^2(\mathbb{R}^d)$ the set of twice continuously differentiable functions on $\mathbb{R}^d$, define a test function class

$$\mathcal{D} := \{x \in C^2(\mathbb{R}^d) : |x(\mathbf{z})| + |\nabla x(\mathbf{z})| + |\nabla^2 x(\mathbf{z})| \leq C_x\, w_2(\mathbf{z}) \text{ for some } C_x > 0\}. \tag{183}$$

Given the mutation matrix $\mathbf{M}$, growth rate $m(\mathcal{N}_t, \mathbf{z})$, and reproductive variance $v \geq 0$, a process $\mathcal{N} \in D_{\bar{\mathfrak{M}}^4}[0,\infty)$ is said to solve the localized $(\mathbf{M}, m, v)$-martingale problem if for every $x \in \mathcal{D}$ and $u = 1,2,\ldots$,

$$\mathcal{M}_t^u(x) := \mathcal{N}_{t\wedge\tau_u}(x) - \mathcal{N}_0(x) - \int_0^{t\wedge\tau_u}\int_{\mathbb{R}^d}(\mathcal{A}\,x(\mathbf{z}) + m(\mathcal{N}_s,\mathbf{z})\,x(\mathbf{z}))\,\mathcal{N}_s(d\mathbf{z})\,ds \tag{184}$$

is a martingale with quadratic covariation

$$[\mathcal{M}^u(x), \mathcal{M}^u(y)]_t = v\int_0^{t\wedge\tau_u}\int_{\mathbb{R}^d} x(\mathbf{z})y(\mathbf{z})\mathcal{N}_s(d\mathbf{z})\,ds, \tag{185}$$

and $\mathcal{N}_t = \partial$ for $t \geq \tau$. The notation $s \wedge t = \min(s,t)$ is common in probability theory. Additionally, it should be noted that *localization* here refers to localization in time (not space).

- - - - - - - - - - - - - - - - - - - - - - - - - - - - - - - - - - - - - - - - - - - - - - - - - - - - - - - - - - - -

##### 7.3.4. Approximating $k$-System

For each $k = 1,2,\ldots$, the $k$-th rescaling of the individual-based model with Brownian motion mutation is summarized by:

- Individual weight: $\alpha_k = 1/k$
- Population measure: $N_t^k = \alpha_k \sum_{i=1}^{n_t^k} \delta_{\mathbf{z}_{i,k}(0)}$
- Initial abundance: $n_0^k = kn(0)$
- Mutation generator: $\mathcal{A} = \frac{1}{2}\nabla^\top\mathbf{M}\nabla$
- Branching rate: $\rho_k = k\rho$
- Fitness: $W_k(N,\mathbf{z}) = W^{1/k}(N,\mathbf{z})$, for $N$ of the form $N = \alpha\sum_i \delta_{\mathbf{z}_i}$
- Offspring distribution: For parent with trait $\mathbf{z}$ in population $N$, Poisson with mean $W_k(N,\mathbf{z})$

Define $m_k(N,\mathbf{z}) = (W_k(N,\mathbf{z}) - 1)\rho_k$ and $v_k(N,\mathbf{z}) = (W_k^2(N,\mathbf{z}) - W_k(N,\mathbf{z}) + 1)\alpha_k\rho_k$. More generally, the following assumptions are made:

- (A1) **Mutation:** Individuals mutate independently with generator $\mathcal{A}$.
- (A2) **Convergence of growth rate:** Given $N^k \to \mathcal{N}$, there exists $m(\mathcal{N}, \mathbf{z})$ such that, for each $u = 1, 2, \ldots$,

$$m_k(N^k, \mathbf{z}) \to m(\mathcal{N}, \mathbf{z}) \quad \text{as } k \to \infty,$$

uniformly on the set $\{(\mathcal{N}, \mathbf{z}) : \mathcal{N}(w_4) \le u,\ |\mathbf{z}| \le u\}$, and $|m(\mathcal{N}, \mathbf{z})| \le C(1 + \mathcal{N}(1) + |\mathbf{z}|^2)$.
- (A3) **Convergence of reproductive variance:** Given $N^k \to \mathcal{N}$, there exists a constant $v > 0$ such that $v_k(N^k, \mathbf{z}) \to v > 0$ (with convergence locally uniformly).
- (A4) **Initial conditions:** $N_0^k(x) \to \mathcal{N}_0(x)$ with $\mathcal{N}_0 \in \mathfrak{M}^4$ for all $x \in \mathcal{D}$, and

$$\sup_k \mathbb{E}[N_0^k(w_4)^p] < \infty \quad \text{for some } p \ge 1.$$

- (A5) **Uniform fourth-moment bound:** For every finite time horizon $T > 0$,

$$\sup_{k\ge1} \mathbb{E}\left[\sup_{0\le t\le T} N_t^k(w_4)\right] < \infty, \qquad w_4(z) = 1 + |z|^4.$$

This controls all localization times $\tau_u^k$ and provides the uniform polynomial moment bounds needed later.

. . . . . . . . . . . . . . . . . . . . . . . . . . . . . . . . . . . . . . . . . . . . . . . . . . . . . . . . . . . . . . . . .

##### *7.3.5. A Martingale Characterization of the Approximating k-System*

For $x \in \mathcal{D}$, we have $N_t^k(x) = \int_{\mathbb{R}^d} x(\mathbf{z})\, N_t^k(d\mathbf{z}) = \alpha_k \sum_{i=1}^{n_t^k} x\big(\mathbf{z}_{i,k}(t)\big)$. I derive a (semi)martingale decomposition for $N_t^k(x)$ and identify the quadratic covariation.

**Brownian mutation:** Between branching events the population size $n_t^k$ is constant and for each $i, k$, $d\mathbf{z}_{i,k} = \sqrt{\mathbf{M}} d\mathbf{B}_{i,k}$. Applying Itô's formula to $x(\mathbf{z}_{i,k})$ gives

$$dx\big(\mathbf{z}_{i,k}\big) = \mathcal{A}x\big(\mathbf{z}_{i,k}\big)\, dt + \nabla x\big(\mathbf{z}_{i,k}\big)^\top \sqrt{\mathbf{M}}\, d\mathbf{B}_{i,k}. \tag{186}$$

Summing over individuals and multiplying by $\alpha_k$ gives

$$dN^{k,\text{mut}}(x) = \alpha_k \sum_{i=1}^{n_k} dx\big(\mathbf{z}_{i,k}\big) = \alpha_k \sum_{i=1}^{n_k} \mathcal{A}x\big(\mathbf{z}_{i,k}\big) dt + dM^{k,\text{mut}}(x), \tag{187}$$

with continuous martingale part

$$M_t^{k,\text{mut}}(x) := \alpha_k \sum_{i=1}^{n_t^k} \int_0^t \nabla x\big(\mathbf{z}_{i,k}(s)\big)^\top \sqrt{\mathbf{M}}\, d\mathbf{B}_{i,k}(s). \tag{188}$$

The deterministic bias of $dN^{k,\text{mut}}$ due to mutation can be written as

$$\alpha_k \sum_i \mathcal{A}x\big(\mathbf{z}_{i,k}\big) = \int_{\mathbb{R}^d} \mathcal{A}x(\mathbf{z})\, N^k(d\mathbf{z}). \tag{189}$$

The quadratic covariation for $x, y \in \mathcal{D}$ in differential notation is

$$\begin{aligned} d[M^{k,\text{mut}}(x), M^{k,\text{mut}}(y)] &= \alpha_k^2 \sum_i \nabla x\big(\mathbf{z}_{i,k}\big)^\top \mathbf{M} \nabla y\big(\mathbf{z}_{i,k}\big)\, ds \\ &= \alpha_k \int_{\mathbb{R}^d} \nabla x(\mathbf{z})^\top \mathbf{M} \nabla y(\mathbf{z})\, N^k(d\mathbf{z})\, ds. \end{aligned} \tag{190}$$

**Branching events:** Now consider the jumps in $N_t^k(x)$ due to branching. Take a population state $N = \alpha_k \sum_{i=1}^n \delta_{\mathbf{z}_i}$ and fix an individual with trait $\mathbf{z}$. When this individual branches:

- The event occurs at rate $\rho_k$.
- The individual is removed and replaced by $K \sim \text{Poisson}(W_k(N, \mathbf{z}))$ offspring.
- Neglecting mutation during the instantaneous branching, all offspring have trait $\mathbf{z}$ at the branching time.

The jump in $N(x)$ from this event is

$$\Delta N(x) = \alpha_k \left( \sum_{j=1}^{K} x(\mathbf{z}) - x(\mathbf{z}) \right) = \alpha_k (K-1)\, x(\mathbf{z}). \tag{191}$$

Conditionally on $N$ and the trait $\mathbf{z}$,

$$\mathbb{E}[\Delta N(x) \mid N, \mathbf{z}] = \alpha_k \big(\mathbb{E}[K] - 1\big)\, x(\mathbf{z}) = \alpha_k \big(W_k(N,\mathbf{z}) - 1\big) x(\mathbf{z}), \tag{192}$$

and

$$\mathbb{E}[(\Delta N(x))(\Delta N(y)) \mid N, \mathbf{z}] = \alpha_k^2\, \mathbb{E}[(K-1)^2]\, x(\mathbf{z}) y(\mathbf{z}). \tag{193}$$

For $K \sim \mathrm{Poisson}(W_k)$,

$$\mathbb{E}[(K-1)^2] = \mathrm{Var}\, K + (\mathbb{E}K - 1)^2 = W_k + (W_k - 1)^2 = W_k^2 - W_k + 1. \tag{194}$$

Summing (192) over all individuals and multiplying by the branching rate $\rho_k$ gives the deterministic bias of $dN^k$ due to branching:

$$\begin{aligned} dN^{k,\mathrm{br}}(x) &= \sum_{i=1}^{n} \rho_k\, \mathbb{E}[\Delta N(x) \mid N, \mathbf{z}_i] \\ &= \sum_i \rho_k\, \alpha_k \big(W_k(N,\mathbf{z}_i) - 1\big) x(\mathbf{z}_i) \\ &= \int_{\mathbb{R}^d} (W_k(N,\mathbf{z}) - 1) \rho_k\, x(z)\, N(d\mathbf{z}). \end{aligned} \tag{195}$$

We have $m_k(N,\mathbf{z}) = (W_k(N,\mathbf{z}) - 1)\rho_k$, which give $dN^{k,\mathrm{br}}(x) = \int_{\mathbb{R}^d} m_k(N,\mathbf{z})\, x(\mathbf{z})\, N(d\mathbf{z})$. Similarly, the quadratic covariation due to branching is given in differential notation by

$$\begin{aligned} d[M^{k,\mathrm{br}}(x), M^{k,\mathrm{br}}(y)] &= \sum_i \rho_k\, \mathbb{E}[\Delta N(x)\, \Delta N(y) \mid N, \mathbf{z}_i]\, ds \\ &= \sum_i \rho_k\, \alpha_k^2 \big(W_k^2(N,\mathbf{z}_i) - W_k(N,\mathbf{z}_i) + 1\big)\, x(\mathbf{z}_i) y(\mathbf{z}_i)\, ds \\ &= \alpha_k \rho_k \int_{\mathbb{R}^d} \big(W_k^2(N,\mathbf{z}) - W_k(N,\mathbf{z}) + 1\big)\, x(\mathbf{z}) y(\mathbf{z})\, N(d\mathbf{z})\, ds. \end{aligned} \tag{196}$$

We have $v_k(N,\mathbf{z}) := \big(W_k^2(N,\mathbf{z}) - W_k(N,\mathbf{z}) + 1\big)\alpha_k \rho_k$. Thus $d[M^{k,\mathrm{br}}(x), M^{k,\mathrm{br}}(y)] = \int_{\mathbb{R}^d} v_k(N^k,\mathbf{z})\, x(\mathbf{z}) y(\mathbf{z})\, N^k(d\mathbf{z})\, ds$.

**Branching and mutation:** Combining the above results we find that, for each $x \in \mathcal{D}$, the process

$$\begin{aligned} M_t^k(x) &:= N_t^k(x) - N_0^k(x) \\ &\quad - \int_0^t \int_{\mathbb{R}^d} \big(\mathcal{A}x(\mathbf{z}) + m_k(N_s^k,\mathbf{z})\, x(\mathbf{z})\big)\, N_s^k(d\mathbf{z})\, ds \end{aligned} \tag{197}$$

is a martingale. Furthermore, because the mutation component of $M_t^k$ is continuous and the branching component is entirely made of jumps, the quadratic covariation of $M_t^k$ for $x, y \in \mathcal{D}$ simplifies to

$$\begin{aligned} [M^k(x), M^k(y)]_t &= [M^{k,\mathrm{mut}}(x), M^{k,\mathrm{mut}}(y)]_t + [M^{k,\mathrm{br}}(x), M^{k,\mathrm{br}}(y)]_t \\ &= \alpha_k \int_0^t \int_{\mathbb{R}^d} \nabla x(\mathbf{z})^\top \mathbf{M} \nabla y(\mathbf{z})\, N_s^k(d\mathbf{z})\, ds \\ &\quad + \int_0^t \int_{\mathbb{R}^d} v_k(N_s^k,\mathbf{z})\, x(\mathbf{z}) y(\mathbf{z})\, N_s^k(d\mathbf{z})\, ds. \end{aligned} \tag{198}$$

(more generally, for martingales $M, N$, $[M+N, M+N] = [M,M] + 2[M,N] + [N,N]$.)

. . . . . . . . . . . . . . . . . . . . . . . . . . . . . . . . . . . . . . . . . . . . . . . . . . . . . . . . . .

##### 7.3.6. *Tightness and Identification of Limit Points of the Approximating k-system on $\bar{\mathfrak{M}}^4$*

To show that the $k$-system converges in $k$ to a solution of the localize $(\mathbf{M}, m, v)$-martingale problem on $\bar{\mathfrak{M}}^4$, we need to first show that limit points of the sequence $\{N^k\}_{k=1}^\infty$ exist. To do so, I outline how one may show that the probability distributions for the $k$-systems in the sequence $\{N^k\}_{k=1}^\infty$ do not spread out too much. That is, the probability distribution of their outcomes remains sufficiently concentrated in $D_{\bar{\mathfrak{M}}^4}[0,\infty)$ so that as $k$ increases the associated $k$-systems do not *wander off to infinity*. This property is called *tightness*. After tightness of $\{N^k\}_{k=1}^\infty$ is established (i.e., that limit points exist), I then provide some details one may use to show that each limit point of $\{N^k\}_{k=1}^\infty$ solves the localized $(\mathbf{M}, m, v)$-martingale problem on $\bar{\mathfrak{M}}^4$. An introduction to the formal theory this approach is based on is given by Billingsley (1999).

**Localization and moment bounds**

For $u = 1, 2, \ldots$, define the $k$-system stopping times

$$\tau_u^k := \inf\{t \geq 0 : N_t^k(w_4) \geq u\}, \qquad \tau_u := \liminf_{k\to\infty} \tau_u^k, \tag{199}$$

where $w_4(\mathbf{z}) := 1 + |\mathbf{z}|^4$. Define the stopped processes $N_t^{k,u}(x) := N_{t\wedge\tau_u^k}^k(x)$, for $x \in \mathcal{D}$ (note that it is *stopped* and not *killed* because it is constant for $t \geq \tau_u^k$ instead of equal to $\partial$). By definition then $N_t^{k,u}(w_4) \leq u$ for all $t \geq 0$. Hence,

$$\sup_k \mathbb{E}\Big[\sup_{0\leq t\leq T} N_t^{k,u}(w_4)^p\Big] \leq u^p, \qquad T > 0. \tag{200}$$

This will be leveraged to apply Lebesgue's Dominated Convergence theorem, and to demonstrate that $\{M^k\}_k$ has a subsequence with a martingale limit point.

**Tightness of one-dimensional projections**

Fix $x \in \mathcal{D}$. From the expression for $M_t^k$ obtained above, define $M_t^{k,u} := M_{t\wedge\tau_u^k}^k$. Then $M_t^{k,u}(x)$ is also a martingale for each $x \in \mathcal{D}$. Setting

$$F_t^{k,u}(x) := \int_0^{t\wedge\tau_u^k} \int_{\mathbb{R}^d} \big(\mathcal{A} + m_k(N_s^k, \mathbf{z})\big)\, x(\mathbf{z})\, N_s^k(d\mathbf{z})\, ds, \tag{201}$$

we can write $N_t^{k,u}(x) = N_0^{k,u}(x) + F_t^{k,u}(x) + M_t^{k,u}(x)$. Initial condition assumption (A4) already gives convergence of $N_0^{k,u}(x)$. Tightness of $\{N_t^{k,u}(x)\}_k$ for fixed $x \in \mathcal{D}$ can be shown by demonstrating tightness of $\{F_t^{k,u}(x)\}_k$ and $\{M_t^{k,u}(x)\}_k$ separately.

For $F_t^{k,u}(x)$, the growth bounds $|x| + |\nabla x| + |\nabla^2 x| \leq C_x(1 + |\mathbf{z}|^2)$ in combination with assumptions (A2) and (A5) can be used to demonstrate $F_t^{k,u}(x) \leq C_{x,u}(1+u)t$ for $0 \leq t \leq T$ and any finite $T > 0$. This implies $F_t^{k,u}(x)$ is Lipschitz in $t$:

$$|F_t^{k,u}(x) - F_s^{k,u}(x)| \leq C_{x,u}(1+u)|t-s| \quad \forall\, s,t \in [0,T], \quad T > 0. \tag{202}$$

The uniform Lipschitz bound together with the growth control places the family $\{F_t^{k,u}(x)\}_k$ in a setting where standard relative-compactness arguments for processes with continuous sample paths apply. Such compactness results are classical in the weak-convergence literature (e.g., Ethier and Kurtz, 1986; Billingsley, 1999).

The quadratic variation associate with $M_t^{k,u}(x)$ is given by

$$\begin{aligned}[M^{k,u}(x)]_t = \alpha_k \int_0^{t\wedge\tau_u^k} \int \nabla x(\mathbf{z})^\top \mathbf{M} \nabla x(\mathbf{z})\, N_s^k(d\mathbf{z})\, ds \\ + \int_0^{t\wedge\tau_u^k} \int v_k(N_s^k, \mathbf{z})\, x^2(\mathbf{z})\, N_s^k(d\mathbf{z})\, ds.\end{aligned} \tag{203}$$

For the subset of sample space where the process has not yet left the localization region by time $s$ (i.e., its $w_4$-moment is still below $u$), denoted by $\{s \leq \tau_u^k\}$, the integrands are bounded by $C_x(1+u)$ using (A1)-(A3) and the growth condition on $x$. Hence, for any fixed $T > 0$,

$$\sup_k \mathbb{E}\big[[M^{k,u}(x)]_T\big] \leq C_{x,u} T. \tag{204}$$

Using the compact-containment estimate together with the martingale moment bounds established above, the family $\{M^{k,u}(x)\}_k$ falls within a setting where standard relative-compactness results for llrc processes apply, and is therefore tight in $D_{\mathbb{R}}[0,T]$ (see, for example, Chapter 3 of Ethier and Kurtz, 1986; or Billingsley, 1999, for general weak-convergence arguments).

Combining the bounds for $F^{k,u}(x)$ and $M^{k,u}(x)$ with (200) and the fact that the jumps of $N^{k,u}(x)$ at branching events are of size $\alpha_k x(\mathbf{z})$ (hence uniformly small in $k$) yields: for each $x \in \mathcal{D}$ and $u \geq 1$, the sequence $\{N^{k,u}(x)\}_k$ is tight in $D_{\mathbb{R}}[0,\infty)$.

**Separation and Convergence-Determining Test Functions**

The space of compactly supported and infinitely differentiable functions $C_c^\infty(\mathbb{R}^d)$ has the property that finite measures are uniquely determined by their integrals against these functions. That is, if $\mu_1(x) = \mu_2(x)$ for all $x \in C_c^\infty(\mathbb{R}^d)$, then $\mu_1 = \mu_2$. This also means that convergence of finite measures $\mu_k \to \mu$ is fully determined by convergence of $\mu_k(x) \to \mu(x)$ for all $x \in C_c^\infty(\mathbb{R}^d)$. Because $C_c^\infty(\mathbb{R}^d) \subset \mathcal{D}$, it follows that our test function class $\mathcal{D}$ is sufficiently rich to distinguish elements of $\mathfrak{M}^4 \subset \mathfrak{M}_F$. This observation motivates the next step.

**Application of Roelly's Tightness Criterion**

Roelly's criterion (Theorems 2.1-2.2 in Roelly-Coppoletta, 1986) requires a *countable convergence-determining family* of test functions. Using the fact above, fix a countable collection $\{x_j\}_{j\geq1} \subset C_c^\infty(\mathbb{R}^d) \subset \mathcal{D}$ constructed, for example, by taking infinitely differentiable bump functions with rational centers, rational radii, and rational coefficients. This family has the property that, for any $y \in C_c^\infty(\mathbb{R}^d)$, there exists a subsequence such that $x_{j_\ell} \to y$ as $\ell \to \infty$. This family is therefore convergence-determining for finite measures, and hence for measures in $\mathfrak{M}^4$.

For each fixed $u \geq 1$, we already saw that every real-valued projection $\{N^{k,u}(x_j)\}_k$, $j \geq 1$, is tight in $D_{\mathbb{R}}[0,\infty)$. Roelly's criterion then implies that the sequence of stopped processes $\{N^{k,u}\}_k$ is tight in $D_{\bar{\mathfrak{M}}^4}[0,\infty)$. This establishes the tightness of the full measure-valued stopped processes, not merely their scalar projections.

**Localization from stopped to full processes**

For each fixed $u \geq 1$, the argument above and Roelly's criterion show that the stopped processes $\{N^{k,u}\}_k$ form a tight family in $D_{\bar{\mathfrak{M}}^4}[0,\infty)$. Moreover, assumption (A5) implies that the localization times

$$\tau_u^k := \inf\{t \geq 0 : N_t^k(w_4) \geq u\} \tag{205}$$

tend to $\infty$ in probability as $u \to \infty$, uniformly in $k$. That is, for every $T > 0$ and $\varepsilon > 0$ there exists $u$ such that

$$\sup_{k \geq 1} \mathbb{P}(\tau_u^k \leq T) \leq \varepsilon. \tag{206}$$

This is a *compact containment* condition that can be used to apply Theorem 7.2 on page 128 of Ethier and Kurtz (1986). That theorem also requires a *modulus-of-continuity* control, which is satisfied here by equation (204) above. Then, as a consequence of Theorem 7.2 in Ethier and Kurtz (1986), we have that the full sequence $\{N^k\}_k$ is tight in $D_{\bar{\mathfrak{M}}^4}[0,\infty)$. Note that Theorem 7.2 in Ethier and Kurtz (1986) is a statement about relative compactness. But in their context relative compactness is the same as tightness.

**Identification of Limit Points as Solutions of the Localized Martingale Problem**

Having established tightness of the full sequence $\{N^k\}_{k\geq 1}$ in $D_{\bar{\mathfrak{M}}^4}[0,\infty)$, there exists a subsequence $\{N^{k_\ell}\}_{\ell \geq 1}$ and an $\bar{\mathfrak{M}}^4$-valued llrc process $\mathcal{N} \in D_{\bar{\mathfrak{M}}^4}[0,\infty)$ such that $N^{k_\ell}(x) \to \mathcal{N}(x)$ in $D_{\mathbb{R}}[0,\infty)$ for each $x \in \mathcal{D}$ (where convergence is a.s. by Skorokhod's representation theorem). Define stopping times for the limit by $\tau_u := \inf\{t \geq 0 : \mathcal{N}_t(w_4) \geq u\}$, for $u = 1, 2, \ldots$, and stopped process $\mathcal{N}_t^u := \mathcal{N}_{t\wedge\tau_u}$. Similarly, for each $k_\ell$, define

$$\tau_u^{k_\ell} := \inf\{t \geq 0 : N_t^{k_\ell}(w_4) \geq u\}, \qquad N_t^{k_\ell,u} := N^{k_\ell}_{t\wedge\tau_u^{k_\ell}}. \tag{207}$$

For any fixed $u \geq 1$ and $x \in \mathcal{D}$, the stopped processes admit the semimartingale decomposition

$$N_t^{k_\ell,u}(x) = N_0^{k_\ell}(x) + F_t^{k_\ell,u}(x) + M_t^{k_\ell,u}(x), \tag{208}$$

where

$$F_t^{k_\ell,u}(x) = \int_0^{t\wedge\tau_u^{k_\ell}} \int_{\mathbb{R}^d} \left(\mathcal{A}x(\mathbf{z}) + m_{k_\ell}(N_s^{k_\ell}, \mathbf{z})\, x(\mathbf{z})\right) N_s^{k_\ell}(d\mathbf{z})\, ds, \tag{209}$$

and the martingale $M_t^{k_\ell,u}(x)$ has predictable quadratic variation

$$\begin{aligned}[M^{k_\ell,u}(x)]_t = \alpha_{k_\ell} &\int_0^{t\wedge\tau_u^{k_\ell}} \int \nabla x(\mathbf{z})^\top \mathbf{M} \nabla x(\mathbf{z})\, N_s^{k_\ell}(d\mathbf{z})\, ds \\ &+ \int_0^{t\wedge\tau_u^{k_\ell}} \int v_{k_\ell}(N_s^{k_\ell}, \mathbf{z})\, x^2(\mathbf{z})\, N_s^{k_\ell}(d\mathbf{z})\, ds.\end{aligned} \tag{210}$$

The localization event $\{s \leq \tau_u^{k_\ell}\}$ is the subset of sample space where $N_t^{k_\ell,u}(w_4) \leq u$ for $t \leq s$. So on this event, we have $N_s^{k_\ell}(w_4) \leq u$, so by assumptions (A1)-(A3) and the growth bounds on $x \in \mathcal{D}$, all drift integrands and bracket integrands are uniformly bounded in $\ell$. Together with (A5), this yields uniform integrability on finite intervals. Since $N^{k_\ell} \to \mathcal{N}$ (a.s.), Lebesgue's dominated convergence theorem gives

$$F_t^{k_\ell,u}(x) \;\to\; \int_0^{t\wedge\tau_u} \int_{\mathbb{R}^d} \left(\mathcal{A}x(\mathbf{z}) + m(\mathcal{N}_s, \mathbf{z})\, x(\mathbf{z})\right) \mathcal{N}_s(d\mathbf{z})\, ds, \tag{211}$$

and

$$[M^{k_\ell,u}(x)]_t \;\to\; v \int_0^{t\wedge\tau_u} \int_{\mathbb{R}^d} x^2(\mathbf{z})\, \mathcal{N}_s(d\mathbf{z})\, ds. \tag{212}$$

To obtain the limit of the quadratic covariation, apply the same argument to $x$, $y$, and $x + y$, and use bilinearity of the quadratic covariation.

Each $M^{k_\ell,u}(x)$ is a square-integrable martingale (that is, $\mathbb{E}[(M_t^{k_\ell,u}(x))^2] < \infty$). The convergence of quadratic covariation on time $[0,T]$ for any $T > 0$, together with uniform integrability and convergence of the finite-variation parts, implies that the limit

$$\mathcal{M}_t^u(x) := \mathcal{N}_{t\wedge\tau_u}(x) - \mathcal{N}_0(x) - \int_0^{t\wedge\tau_u} \int_{\mathbb{R}^d} \left(\mathcal{A}x(\mathbf{z}) + m(\mathcal{N}_s, \mathbf{z})\, x(\mathbf{z})\right) \mathcal{N}_s(d\mathbf{z})\, ds \tag{213}$$

is a martingale with quadratic covariation

$$[\mathcal{M}^u(x), \mathcal{M}^u(y)]_t = v \int_0^{t\wedge\tau_u} \int_{\mathbb{R}^d} x(\mathbf{z}) y(\mathbf{z})\, \mathcal{N}_s(d\mathbf{z})\, ds. \tag{214}$$

Define $\tau := \lim_{u\to\infty} \tau_u$ and set $\mathcal{N}_t = \partial$ for all $t \geq \tau$. This matches the killed-process construction used to define the localized martingale problem. In conclusion, for every $u \geq 1$ and every test function $x \in \mathcal{D}$, $\mathcal{M}_t^u = \mathcal{M}_{t\wedge\tau_u}$ is a martingale with quadratic covariation given by (185). Because $u$ was arbitrary, the above calculations offer compelling evidence that the diffusion-limit $\mathcal{N}$ exists and is a solution of the localized $(\mathbf{M}, m, v)$-martingale problem on $D_{\bar{\mathfrak{M}}^4}[0,\infty)$.

This outlines one potential route to formally construct the framework in the main text via a diffusion-limit of an individual-based model. The calculations above use an underlying model in which mutation is represented by multivariate Brownian motion. Replacing this with an individual-based model where mutation occurs only through Gaussian jumps at branching events alters some local details of the semimartingale decomposition (mutation then enters through the jump mechanism rather than the continuous part), but it does not change the overall limiting argument. That is, after the usual small-jump/high-rate rescaling, the same generator $\mathcal{A}$ and the same limiting process $\mathcal{N}$ appear. Likewise, models that decouple death and branching (so individuals may branch several times before death), once rescaled appropriately, converge to the same limit $\mathcal{N}$. Taken together, these observations show that a broad class of individual-based population models fall into the same diffusion-limit, reinforcing the utility and robustness of the framework developed in the main text for analyzing diverse biological population dynamics.

#### 8. The Reproductive Variance Parameter as Gillespie's Variance in Offspring Number

To show that the reproductive variance $v > 0$ is equal to Gillespie's (1974; 1975; 1977) variance in offspring number (especially when $v$ does not covary with genotype), start by considering the case where $v = v(\mathbf{z})$ depends on trait value. For the univariate trait case, a stochastic extension of DAGA was introduced by Week et al., (2021) that accounts for $v(z)$. In the absence of mutation, this equation reads $\dot{\nu}_t(z) = m(z)\nu_t(z) + \sqrt{v(z)\nu_t(z)}\dot{W}_t(z)$, with $\dot{W}_t(z)$ a space-time white noise. For the sake of comparison with the martingale approach used here, the results in Méléard and Roelly (1993) demonstrate that solutions to this stochastic partial differential equation can also be characterized by solutions to a martingale problem:

$$\mathcal{M}_t(x) = \int_{\mathbb{R}} x(z)\mathcal{N}_t(dz) - \int_{\mathbb{R}} x(z)\mathcal{N}_0(dz) - \int_0^t \int_{\mathbb{R}} m(z)x(z)\mathcal{N}_s(dz)ds \tag{215}$$

is a martingale with quadratic covariation

$$[\mathcal{M}_t(x), \mathcal{M}_t(y)]_t = \int_0^t \int_{\mathbb{R}} v(z)x(z)y(z)\mathcal{N}_s(dz)ds. \tag{216}$$

It turns out that the solution to this martingale problem, $\mathcal{N}_t(z)$, admits a density $\nu_t(z)$ that solves the stochastic extension of DAGA without mutation such that $\mathcal{N}_t(dz) = \nu_t(z)dz$ (Li, 1998).

Gillespie's (1974; 1975; 1977) primary result regarding the covariance of genotype with variance in offspring number states that, when two genotypes confer the same average number of offspring, selection favors the one with less variance in offspring number. Here I show how to recover this result from the stochastic extension of DAGA without mutation. In particular, when $v(z)$ depends on trait value, a form of selection emerges due to the covariance $\mathrm{Cov}(v,z) = \int_{\mathbb{R}}(v(z) - \bar{v})(z - \bar{z})p(z)dz$, where $\bar{v} = \int_{\mathbb{R}} v(z)p(z)dz$ and $p(z) = \nu(z)/n$, $n = \int_{\mathbb{R}} \nu(z)dz$. To demonstrate this, I start by deriving dynamics of the trait distribution $p$ from $\dot{\nu}$.

From hereon, I write $\mathcal{M}(x)$ in place of $\int_{\mathbb{R}} x(z)\mathcal{M}(dz)$, and I use stochastic differential notation so that $d\mathcal{N}(x) = \mathcal{N}(mx)dt + d\mathcal{M}(x)$ communicates the same statement as equation (215) above. With this notation, the heuristics presented in the main text are extended here using equation (216). For example, abundance dynamics are given by setting $x = 1$. Because $n = \mathcal{N}(1)$, this gives $dn = \bar{m}\,n\,dt + \sqrt{\bar{v}\,n}\,dB_n$ for some Brownian motion $B_n := \mathcal{M}(\hat{1})$.

For each $x$, set $\mathcal{P}(x) := \mathcal{N}(x)/n$ so that $\mathcal{P}(x) = \int_{\mathbb{R}} x(z)p(z)dz$. For fixed $x$, apply Itô to $\mathcal{N}(x)/n$:

$$d\mathcal{P}(x) = \frac{1}{n}d\mathcal{N}(x) - \frac{\mathcal{N}(x)}{n^2}dn + \frac{\mathcal{N}(x)}{n^3}d[\mathcal{M}(1)]_t - \frac{1}{n^2}d[\mathcal{M}(x), \mathcal{M}(1)]_t. \tag{217}$$

Substitute $d\mathcal{N}(x) = \mathcal{N}(mx)dt + d\mathcal{M}(x)$, $dn = \mathcal{N}(m)dt + d\mathcal{M}(1)$, $d[\mathcal{M}(x), \mathcal{M}(1)]_t = \mathcal{N}(vx)dt$, and $d[\mathcal{M}(1)]_t = \mathcal{N}(v)dt$:

$$d\mathcal{P}(x) = \frac{1}{n}(\mathcal{N}(mx)dt + d\mathcal{M}(x)) - \frac{\mathcal{N}(x)}{n^2}(\mathcal{N}(m)dt + d\mathcal{M}(1)) + \frac{\mathcal{N}(x)}{n^3}\mathcal{N}(v)dt - \frac{1}{n^2}\mathcal{N}(vx)dt. \tag{218}$$

Rewrite using $\mathcal{N}(x) = n\mathcal{P}(x)$:

$$d\mathcal{P}(x) = \mathcal{P}(mx)dt - \mathcal{P}(x)\mathcal{P}(m)dt - \frac{1}{n}(\mathcal{P}(vx) - \mathcal{P}(x)\mathcal{P}(v))dt + \frac{1}{n}d\mathcal{M}(x) - \frac{\mathcal{P}(x)}{n}d\mathcal{M}(1). \tag{219}$$

Define $\mathrm{Cov}_{\mathcal{P}}(a,x) := \mathcal{P}(ax) - \mathcal{P}(a)\mathcal{P}(x)$ and $d\mathcal{M}^{\mathcal{P}}(x) := \frac{1}{n}d\mathcal{M}(x) - \frac{\mathcal{P}(x)}{n}d\mathcal{M}(1)$. Then

$$d\mathcal{P}(x) = (\mathrm{Cov}_{\mathcal{P}}(m,x) - \frac{1}{n}\mathrm{Cov}_{\mathcal{P}}(v,x))dt + d\mathcal{M}^{\mathcal{P}}(x). \tag{220}$$

##### 8.1. Gillespie as a special case: two discrete types

Fix a subset of trait space $A \subset \mathbb{R}$ with positive initial frequency $0 < \mathcal{P}_0(A) < 1$ and define the indicator $x_A(z) := \mathbf{1}_{\{z \in A\}}$. The frequency of $A$ at time $t$ is

$$p_t := \mathcal{P}_t(x_A) = \mathcal{P}_t(A). \tag{221}$$

Define conditional means of $m$ and $v$ on $A$ and its complement $A^c$:

$$m_+ := \frac{\mathcal{P}_t(mx_A)}{p_t}, \quad m_- := \frac{\mathcal{P}_t(m(1 - x_A))}{1 - p_t}, \tag{222}$$

$$v_+ := \frac{\mathcal{P}_t(vx_A)}{p_t}, \quad v_- := \frac{\mathcal{P}_t(v(1 - x_A))}{1 - p_t}. \tag{223}$$

Then

$$\mathcal{P}_t(m) = p_t m_+ + (1 - p_t)m_-, \quad \mathcal{P}_t(mx_A) = p_t m_+, \tag{224}$$

so the covariance term becomes

$$\mathrm{Cov}_{\mathcal{P}_t}(m, x_A) = \mathcal{P}_t(mx_A) - \mathcal{P}_t(m)\mathcal{P}_t(x_A) = p_t(1 - p_t)(m_+ - m_-). \tag{225}$$

Similarly,

$$\mathcal{P}_t(v) = p_t v_+ + (1 - p_t)v_-, \quad \mathcal{P}_t(vx_A) = p_t v_+, \tag{226}$$

so

$$\mathrm{Cov}_{\mathcal{P}_t}(v, x_A) = p_t(1 - p_t)(v_+ - v_-). \tag{227}$$

Now plug $x = x_A$ into the general SDE:

$$dp_t = d\mathcal{P}_t(x_A) = (\mathrm{Cov}_{\mathcal{P}_t}(m, x_A) - \frac{1}{n_t}\mathrm{Cov}_{\mathcal{P}_t}(v, x_A))dt + d\mathcal{M}_t^{\mathcal{P}}(x_A). \tag{228}$$

Using the covariance expressions above, the deterministic term of $dp_t$ is

$$\det(dp_t) = p_t(1 - p_t)\big[(m_+ - m_-) - \frac{v_+ - v_-}{n_t}\big]. \tag{229}$$

Thus the Gillespie effective selection coefficient associated with the partition $(A, A^c)$ is

$$s_{\mathrm{eff}} = (m_+ - m_-) - \frac{v_+ - v_-}{n_t}. \tag{230}$$

For the stochastic term, use the quadratic variation of $\mathcal{M}^{\mathcal{P}}(x)$:

$$\frac{d}{dt}[\mathcal{M}^{\mathcal{P}}(x)]_t = \frac{1}{n_t}\int (x(z) - \mathcal{P}_t(x))^2 v(z)\mathcal{P}_t(dz). \tag{231}$$

With $x = x_A$ and $\mathcal{P}_t(x_A) = p_t$,

$$\int (x_A(z) - p_t)^2 v(z)\mathcal{P}_t(dz) = \int_A (1 - p_t)^2 v(z)\mathcal{P}_t(dz) + \int_{A^c} p_t^2 v(z)\mathcal{P}_t(dz). \tag{232}$$

Using the definitions of $v_+$ and $v_-$ this simplifies to

$$\int (x_A - p_t)^2 v\, d\mathcal{P}_t = p_t(1 - p_t)\big[(1 - p_t)v_+ + p_t v_-\big]. \tag{233}$$

Hence

$$\frac{d}{dt}[p]_t = \frac{p_t(1 - p_t)}{n_t}\big[(1 - p_t)v_+ + p_t v_-\big]. \tag{234}$$

Collecting deterministic and stochastic terms, the sde for the frequency of the general subset $A$ is

$$dp_t = p_t(1 - p_t)\big[(m_+ - m_-) - \frac{v_+ - v_-}{n_t}\big]dt + \sqrt{\frac{p_t(1 - p_t)}{n_t}\big[(1 - p_t)v_+ + p_t v_-\big]}\, dB_+, \tag{235}$$

which is exactly the Gillespie form for the two "types" defined by $A$ and $A^c$, for any measurable $A$ with initial $\mathcal{P}_0(A) \in (0, 1)$.

Now write the population means as

$$\bar{m}_t := \mathcal{P}_t(m) = p_t m_+ + (1 - p_t)m_-, \quad \bar{v}_t := \mathcal{P}_t(v) = p_t v_+ + (1 - p_t)v_-. \tag{236}$$

For the deterministic term, note

$$p_t(1 - p_t)(m_+ - m_-) = p_t(m_+ - \bar{m}_t) = (1 - p_t)(\bar{m}_t - m_-), \tag{237}$$

and likewise

$$p_t(1 - p_t)(v_+ - v_-) = p_t(v_+ - \bar{v}_t) = (1 - p_t)(\bar{v}_t - v_-). \tag{238}$$

Using these identities, the deterministic term can be rewritten as

$$\det(p_t) = p_t(m_+ - \bar{m}_t) - \frac{p_t}{n_t}(v_+ - \bar{v}_t), \tag{239}$$

or equivalently

$$\det(p_t) = (1 - p_t)(\bar{m}_t - m_-) - \frac{1 - p_t}{n_t}(\bar{v}_t - v_-). \tag{240}$$

For the stochastic term, use

$$(1 - p_t)v_+ + p_t v_- = \bar{v}_t + (1 - 2p_t)(v_+ - v_-), \tag{241}$$

so the variance term becomes

$$\frac{d}{dt}[p]_t = \frac{p_t(1 - p_t)}{n_t}\big[\bar{v}_t + (1 - 2p_t)(v_+ - v_-)\big]. \tag{242}$$

Thus the sde can be expressed in terms of $\bar{m}_t$ and $\bar{v}_t$ as

$$dp_t = \Big(p_t(m_+ - \bar{m}_t) - \frac{p_t}{n_t}(v_+ - \bar{v}_t)\Big)dt + \sqrt{\frac{p_t(1 - p_t)}{n_t}\big[\bar{v}_t + (1 - 2p_t)(v_+ - v_-)\big]}\, dB_+. \tag{243}$$

Recall the abundance dynamics $dn = \bar{m}\, n\, dt + \sqrt{\bar{v}\, n}\, dB_n$. Compare this with the abundance dynamics in the main text, $dn = \bar{m}\, n\, dt + \sqrt{v\, n}\, dB_n$, where $v > 0$ is constant. This identifies the reproductive variance parameter $v$ with Gillespie's variance in offspring number. This identification should not be taken literally: "variance in offspring number" is only one possible interpretation. The diffusion-limit construction above shows that $v$ can equally be interpreted as an effective reproduction rate. In particular, in standard birth–death models one has $v = b + d$, the sum of per-capita birth and death rates (see Week et al., 2021).

B. Øksendal. *Stochastic differential equations: an introduction with applications.* Springer Science & Business Media, 2013.
